# Polθ is a functional dependency and therapeutic vulnerability in DNMT3A deficient leukemia cells

**DOI:** 10.1101/2024.09.15.613155

**Authors:** Bac Viet Le, Umeshkumar Vekariya, Monika M. Toma, Margaret Nieborowska-Skorska, Marie-Christine Caron, Malgorzata Gozdecka, Zayd Haydar, Martin Walsh, Jayashri Ghosh, Elaine Vaughan-Williams, Paulina Podszywalow-Bartnicka, Anna-Mariya Kukuyan, Sylwia Ziolkowska, Jessica Atkins, Emir Hadzijusufovic, Gurushankar Chandramouly, Reza Nejati, Katarzyna Piwocka, Richard Pomerantz, George S. Vassiliou, Brian J.P. Huntly, Peter Valent, Mariusz Wasik, Alfonso Bellacosa, Jean-Yves Masson, Gaorav P. Gupta, Grant A. Challen, Tomasz Skorski

## Abstract

Myeloid malignancies carrying somatic *DNMT3A* mutations (*DNMT3Amut*) may be refractory to standard therapy. *DNMT3Amut* leukemia cells accumulate toxic DNA double strand breaks (DSBs) and stalled replication forks, rendering them dependent on DNA damage response (DDR). We report here that DNA polymerase theta (Polθ), a key element in DSB repair by end-joining (TMEJ) and in fork restarting, is essential for survival and proliferation of *DNMT3Amut* leukemia cells. Polθ is overexpressed in *DNMT3Amut* leukemia cells due to abrogation of PARP1 PARylation-dependent UBE2O E3 ligase-mediated ubiquitination and proteasomal degradation of Polθ. In addition, PARP1-mediated recruitment of the SMARCAD1-MSH2/MSH3 repressive complex to DSBs was diminished in *DNMT3Amut* leukemia cells which facilitated loading of Polθ on DNA damage and promoting TMEJ and replication fork restart. Polθ inhibitors enhanced the anti-leukemic effects of standard drugs such as FLT3 kinase inhibitor quizartinib, cytarabine +/- doxorubicin, and etoposide *in vitro* and in mice with *DNMT3Amut* leukemia. Altogether, Polθ is an attractive target in *DNMT3Amut* hematological malignancies.

## Introduction

DNA (cytosine-5)-methyltransferase 3A (DNMT3A) loss of function either by somatic mutation, allelic loss and epigenetic-dependent abrogation of expression has been found in solid tumors (e.g., lung and colon cancers, melanoma) and in hematological malignancies such as acute myeloid leukemia (AML), myeloproliferative neoplasm (MPN) and chronic myeloid leukemia (CML) blast phase (AACR, 2017; Kim et al., 2013; Sato et al., 2016; Zhang et al., 2020a). DNMT3A adds methyl groups to the cytosine residues in DNA, generating 5-methylcytosine (5-mC) (Okano et al., 1999). Somatic *DNMT3A* mutations lead to a loss of protein function and are associated with reduced sensitivity of hematological malignancies to standard treatments such as anthracyclines (including daunorubicin, the component of standard “7+3” induction chemotherapy for AML), interferon alpha (IFNα), BCL-2 inhibitor venetoclax, and ABL1 tyrosine kinase inhibitor imatinib (Guryanova et al., 2016; Knudsen et al., 2022; Mohanty et al., 2024; Nteliopoulos et al., 2019), which may result in unfavorable outcomes and adverse prognosis (Knudsen et al., 2022; Ley et al., 2013; Liang et al., 2019; Nangalia et al., 2015; Nteliopoulos et al., 2019; Rinke et al., 2020). Also, *DNMT3A*-mutated clones are commonly detected during remission, likely contributing to disease relapse (Klco et al., 2015; Knudsen et al., 2022). Altogether, these observations suggest that malignant clones carrying *DNMT3A* mutations are difficult to eliminate. Thus, novel therapeutic strategies against *DNMT3A* mutated hematological malignancies need to be developed.

AML, MPN and CML cells expressing oncogenic tyrosine kinases (OTKs) such as FLT3(ITD), JAK2(V617F), MPL(W515L) and BCR-ABL1 accumulate spontaneous DNA damage, including highly toxic DNA double strand breaks (DSBs), induced by metabolic products and replication stress (Chen et al., 2014; Esposito and So, 2014; Nieborowska-Skorska et al., 2017a; Vekariya et al., 2023). To repair abundant DSBs and protect the cells from pro-apoptotic effect of DSBs, leukemia cells might activate the DNA damage response (DDR) involving the pathways sensing DSBs (ATM and ATR kinases), repairing DSBs (RAD51-mediated homologous recombination = HR, RAD52-mediated transcription associated homologous recombination = TA-HR and single strand annealing = SSA, DNA-PK-mediated non-homologous end-joining = NHEJ, Polθ-mediated end-joining = TMEJ) as well as those activating checkpoints to arrest cell cycle (via CHK1 and CHK2 kinases) (Jackson, 2002; Le et al., 2022). In addition, PARP1 is required in various aspects of DDR (Ray Chaudhuri and Nussenzweig, 2017).

We reported that somatic mutations in *TET2* (ten-eleven translocation 2) caused HR and NHEJ deficiencies resulting in sensitivity of AML/MPN/CML cells to PARP1 inhibition; conversely, leukemia cells carrying *DNMT3A* mutations were resistant (Maifrede et al., 2021). Here we show that DNA polymerase theta (Polθ), a key element in TMEJ, is essential for survival of AML/MPN/CML cells carrying *DNMT3A* mutations, but not these with *TET2* mutations.

Polθ, encoded by *POLQ* gene, is a large protein (290 kDa), and includes an amino-terminal helicase domain and a carboxy-terminal polymerase domain (Ramsden et al., 2021). Polθ helicase domain captures of two ssDNA strands with microhomologies, bringing them in close proximity (Fijen et al., 2024). Then, Polθ polymerase domain binds to form a synaptic complex and initiate repair synthesis. In addition to playing a key role in TMEJ, Polθ also works in restarting stalled replication forks and in ssDNA gap filling (Belan et al., 2022; Mann et al., 2022; Schrempf et al., 2022; Wang et al., 2019; Wyatt et al., 2016). Polθ is emerging as a therapeutic target in cancer because of the synthetic lethal effect of inhibition of Polθ in HR-deficient tumors (Ramsden et al., 2021).

Although DNMT3A deficiency did not cause detectable changes in HR in tumor cells (Maifrede et al., 2021; Venugopal et al., 2022), we show that these cells are highly sensitive to pharmacological and genetic inactivation of Polθ. Remarkably, we discovered that DNMT3A inactivation promoted Polθ overexpression due to inhibition of PARP1-mediated PARylation-dependent ubiquitination (PARdU) of Polθ by UBE2O E3 ligase resulting in reduced proteasomal degradation of Polθ. DNMT3A-deficient leukemia cells displayed increased loading of Polθ on DNA damage, limited end resection, and enhanced TMEJ and replication fork restart. We also demonstrated that Polθ inhibitors combined with standard drugs could be used to eradicate *DNMT3A*-mutated malignant clones, which may not be eliminated by current therapeutics.

## Results

### DNMT3A deficient malignant hematopoietic cells are addicted to Polθ

FLT3(ITD)-positive murine hematopoietic 32Dcl3 cells and isogenic cells lacking TET2 [FLT3(ITD);*Tet2^KD^*] or DNMT3A [FLT3(ITD);*Dnmt3a^KD^*] were screened for their sensitivity to doxorubicin and DDR inhibitors targeting various aspects of DSB repair (Figure 1A). As expected, FLT3(ITD);*Dnmt3a^KD^* cells were resistant to doxorubicin and PARPi when compared to FLT3(ITD) and FLT3(ITD);*Tet2^KD^* counterparts (Figure 1B) (Guryanova et al., 2016; Maifrede et al., 2021). Moreover, FLT3(ITD);*Dnmt3a^KD^* cells displayed resistance to most of the tested DDR inhibitors in concordance with our recent report involving AML patient cells (Toma et al., 2024). The high overall toxicity of the RAD51 inhibitor most likely resulted from inhibition of HR, a key DSB repair pathway in proliferating cells (Jackson, 2002).

**Figure 1.**
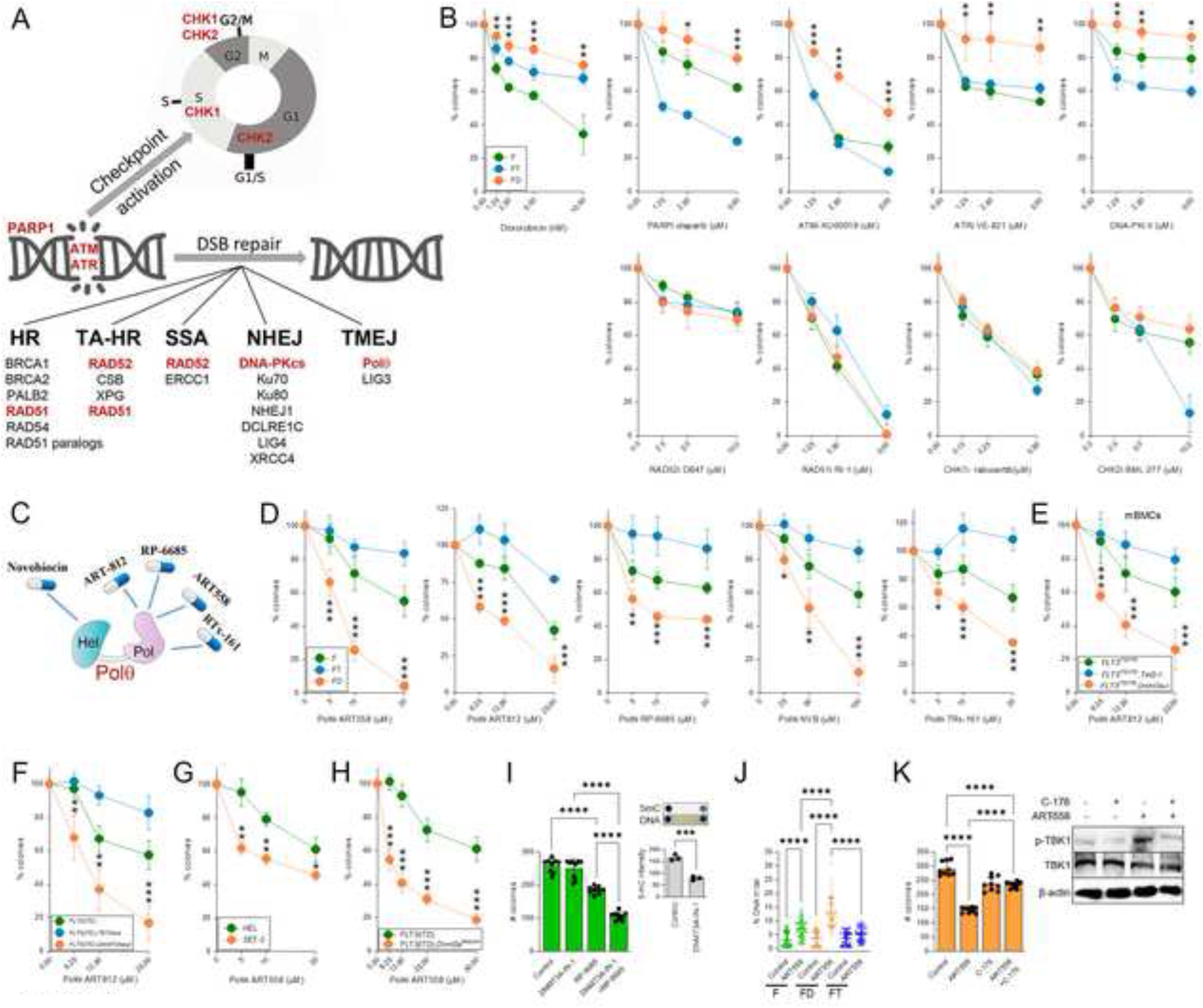
Inactivation of DNMT3A selectively hypersensitized leukemia cells to Polθi. (**A**) A diagram illustrating DSB repair pathways. HR = homologous recombination, TA-HR = transcription associated HR, SSA = single strand annealing, NHEJ = non-homologous end-joining, TMEJ = Polθ-mediated end-joining. Proteins highlighted in red were targeted with small molecule inhibitors. Inset: a representative accumulation of γH2AX foci (DSBs) in leukemia cells. (**B**) FLT3(ITD)-positive 32Dcl3 cells (F) and isogenic cells lacking DNMT3A [FLT3(ITD);*Dnmt3a^KD^* = FD) or TET2 (FLT3(ITD);*Tet2^KD^* = FT] were treated with doxorubicin and the indicated inhibitors. Results represent mean % of colonies relative to untreated control. (**C**) Polθ inhibitors tested here; Hel = helicase domain, Pol = DNA polymerase domain. (**D-H**) F, FD ad FT 32Dcl3 cells (**D**), *Flt3^ITD/ITD^*, *Flt3^ITD/ITD^*;*Dnmt3a^-/-^* and *Flt3^ITD/ITD^*;*Tet2^-/-^*Lin^-^cKit^+^ mBMCs (**E**), Lin^-^CD34^+^ AML patient cells (n=3 of each) carrying FLT3(ITD), FLT3(ITD);*DNMT3A^mut^*and FLT3(ITD);*TET2^mut^* (**F**), JAK2(V617F) human HEL cells and JAK2(V617F);DNMT3A(R882H) human SET-2 cells (**G**), and *Flt3^ITD/+^*and *Flt3^ITD/+^;Dnmt3a^R882H/+^* Lin^-^cKit^+^ mBMCs (**H**) were treated with the indicated Polθ inhibitors. Results represent mean % of colonies relative to untreated control. (**I**) Mean number of colonies of F 32Dcl3 cells treated, or not, with 4 μM DNMT3A-IN-1 and/or 10 μM RP-6685. *Right panels*: Detection of DNMT3A-IN-1-mediated inhibition of 5-mC. (**J**) F (green), FD (orange), and FT (blue) 32Dcl3 cells were untreated (-) or treated with 20 μM ART558 for 24 hrs. Results represent mean % of DNA in the tail. (**K**) FD 32Dcl3 cells were untreated or treated with 5 μM ART558, 0.5 μM C-176 and the combination. *Left panel:* Mean number of colonies. *Right panel*: Representative Western blot detecting phospho-TBK1 kinase, total TBK1 and β-actin (loading control). Error bars represent the SD; p values were determined using one-way Anova (**B**, **D**-**F**, **I**-left panel, **J** and **K**) and Student’s t test (**G, H, I**-Right panel).

Remarkably, FLT3(ITD);*Dnmt3a^KD^* 32Dcl3 cells were selectively highly sensitive to five different Polθ inhibitors which reduced TMEJ: ATPase inhibitor novobiocin (NVB), and DNA polymerase inhibitors ART558, ART812, RP-6685 and RTx-161, when compared to FLT3(ITD) and FLT3(ITD);*Tet2^KD^*counterparts (Figure 1C, D) (Bubenik et al., 2022; Fried et al., 2024; Stockley et al., 2022; Vekariya et al., 2023; Zatreanu et al., 2021; Zhou et al., 2021). In addition, murine Lin-cKit+ *Flt3^ITD/ITD^;Dnmt3a^-/-^*bone marrow cells were more sensitive to Polθi ART812 when compared to their *Flt3^ITD/ITD^*counterparts (Figure 1E). Importantly, human Lin-CD34+ *FLT3^ITD^;DNMT3A^R882H^*AML primary cells were highly sensitive to ART812 when compared to *FLT3^ITD^* patient cells (Figure 1F). In addition, *JAK2^V617F^;DNMT3A^R882H^* SET-2 cells and murine Lin^-^cKit^+^ *Flt3^ITD+/-^;Dnmt3a^R882H/+^*cells displayed sensitivity to ART558 when compared to JAK2^V617F^ HEL and *Flt3^ITD+/-^* murine Lin^-^cKit^+^ cells, respectively (Figure 1G, H). Also, inhibition of DNMT3A activity in FLT3(ITD) 32Dcl3 cells by a selective small molecule inhibitor DNMT3A-IN-1 [validated by reduction of 5-mC (Figure 1I, right panel)] resulted in increased sensitivity to Polθi (Figure 1I, left panel). Of note, FLT3(ITD);*Tet2^KD^* 32Dcl3 cells, *Flt3^ITD/ITD^;Tet2^-/-^* mBMCs, and human FLT3(ITD);*TET2mut* AML primary cells were resistant to NVB, ART558, ART812 and RP-6685 (Figure 1D-F).

ART558 treatment of FLT3(ITD);*Dnmt3a^KD^* 32Dcl3 cells resulted in accumulation of DSBs detected by neutral comet assay (Figure 1J) supporting the hypothesis that inhibition of TMEJ led to accumulation of toxic DSBs. Polθi ART558 exerted its anti-leukemia activity, at least partially, by activating DNA damage-dependent pro-apoptotic cGAS-STING pathway because STING inhibitor (STINGi) C-176 reduced the growth inhibitory effect of Polθi ART558 in FLT3(ITD);*Dnmt3a^KD^* 32Dcl3 cells (Figure 1K, left panel) (Zheng et al., 2023). As expected, treatment with ART558 induced phosphorylation of TBK1 [a marker of active STING (Haag et al., 2018)] in FLT3(ITD);*Dnmt3a^KD^* 32Dcl3 cells, which was inhibited by C-176 (Figure 1K, right panel). However, treatment with ART558 and/or C-176 did not modulate phosphorylation of TBK1 and did not enhance the proliferation of FLT3(ITD);*Tet2^KD^* 32Dcl3 cells which are resistant to Polqi (Supplemental Figure S1).

Next, we determined the source of spontaneous DSBs in FLT3(ITD);*Dnmt3a^KD^*32Dcl3 cells. We reported that in leukemia cells enhanced production of formaldehyde and reactive oxygen species (ROS) resulted in increased numbers of formaldehyde-induced DNA protein crosslinks (DPCs) and oxidative DNA lesions such as 8-oxoguanine (8-oxoG), respectively; unresolved DPCs and 8-oxoG may produce DSBs (Nieborowska-Skorska et al., 2012; Vekariya et al., 2023). We show here that FLT3(ITD);*Dnmt3a^KD^* 32Dcl3 cells and Lin^-^cKit^+^ *Flt3^ITD/ITD^;Dnmt3a^-/-^*mBMCs accumulate more formaldehyde and/or DPCs when compared to FLT3(ITD) and also FLT3(ITD);*Tet2^KD^* counterparts (Supplemental Figure S2A,B). Since *Dnmt3a^KD^* did not enhance the expression of SPRTN protease and TDP1 and TDP2 hydrolases (Supplemental Figure S2C), the proteins directly involved in DPCs removal (Stingele et al., 2017), it is unlikely that they provide sufficient protection against abundant DPCs in DNMT3A-deficient leukemia cells. Unresolved DPCs will ultimately generate DSBs likely contributing to the hypersensitivity to Polθ inhibitors (Chandramouly et al., 2021; Vekariya et al., 2023). In keeping with this, FLT3(ITD);*Dnmt3a^KD^* 32Dcl3 cells were resistant to Polθ inhibitor when cultured in glucose-, serine-, and glycine-free medium (No GSG) which abrogated serine/1C cycle metabolism-mediated generation of formaldehyde-induced DPCs (Supplemental Figure S2D, left) (Vekariya et al., 2023). Conversely, FLT3(ITD);*Dnmt3a^KD^* 32Dcl3 cells remained hypersensitive to Polθ inhibitor in high glucose-, serine-, and glycine-culture medium (High GSG) which facilitated production of formaldehyde and accumulation of DPCs (Supplemental Figure S2D, right). FLT3(ITD);*Dnmt3a^KD^* 32Dcl3 cells, however, did not accumulate more ROS and 8-oxoG lesions when compared to FLT3(ITD) and FLT3(ITD);*Tet^KD^* counterparts (Supplemental Figure S2E and F, respectively). Thus, it is likely that Polθ plays a major role in the repair of DPCs - but not ROS-induced DSBs to promote survival of FLT3(ITD);*Dnmt3a^KD^* 32Dcl3 cells.

To obtain additional evidence about the role of Polθ in *DNMT3A*-mutated hematological malignancies, Polθ was inactivated by introducing a point mutation in *Polq* exon 1 to create a premature stop codon (*Polq*^-/-^ mice) and by expression of Polθ D2230A+Y2231A polymerase inactive mutant. While deletion of *Polq* gene did not affect clonogenic potential of *Dnmt3a^-/-^*mBMCs, it had a detrimental effect on the clonogenic activity of the cells transformed with FLT3(ITD), JAK2(V617F) and BCR-ABL1 oncogenic tyrosine kinases (Figure 2A). This effect was associated with elevated DSBs (γH2AX) and decreased engraftment of GFP^+^;FLT3(ITD);*Dnmt3a^-/-^;Polq^-/-^*cells in mice when compared to GFP^+^;FLT3(ITD);*Dnmt3a^-/-^*counterparts (Figure 2B and C, respectively).

**Figure 2.**
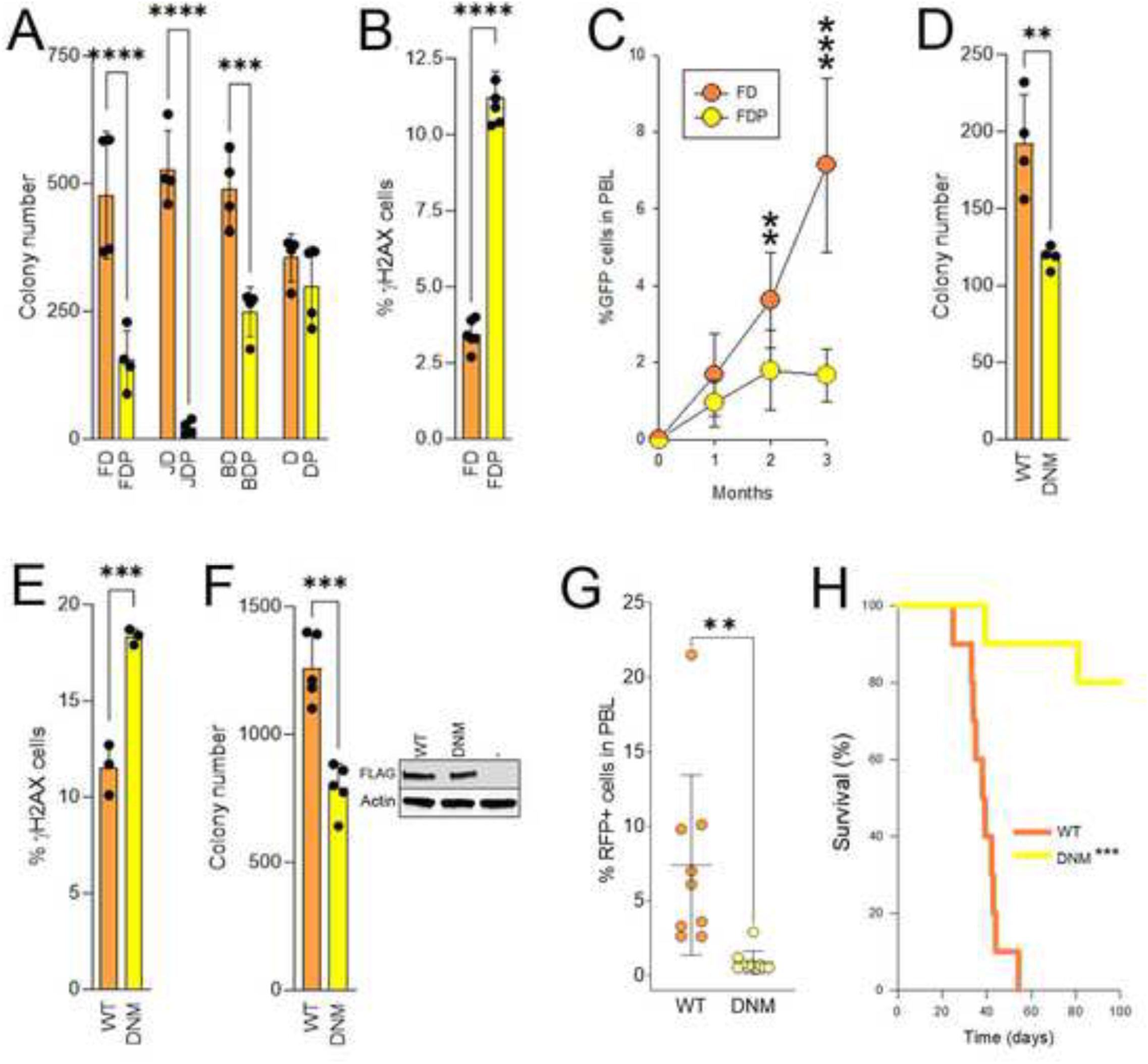
Detrimental effect of genetic inactivation of Polθ in DNMT3A deficient leukemia cells. (**A-C**) GFP^+^ Lin^-^cKit^+^GFP^+^ *Dnmt3a^-/-^* (D) and *Dnmt3a^-/-^;Polq^-/-^* (DP) mBMCs expressing FLT3(ITD) (FD and FDP, respectively), JAK2(V617F) (JD and JDP, respectively) and BCR-ABL1 (BD and BDP, respectively) were examined to detect mean (**A**) colony number, (**B**) % of γH2AX-positive cells, and (**C**) % of GFP^+^ cells in PBL of transplanted mice (n=5-9 mice/group). (**D, E**) *Flt3^ITD/+^;Dnmt3a^R882H/+^*Lin^-^cKit^+^RFP^+^ mBMCs transduced with Polθ wild-type (WT) and D2230A+Y2231A dominant-negative mutant (DNM) were tested to detect (**D**) mean colony number and (**E**) % of γH2AX-positive cells. (**F-H**) RFP^+^ FLT3(ITD);*Dnmt3a^KD^* 32Dcl3 cells transduced with Polθ wild-type (WT) and D2230A+Y2231A dominant-negative mutant (DNM) were tested to measure mean (**F**) number of colonies *in vitro* (*Inset-* Western blot), (**G**) % of RFP^+^ cells in PBL of transplanted SCID mice (n=10 mice/group), and (**H**) survival of the transplanted mice. Error bars represent the SD; p values were determined using Student’s t test (**A-G**); mice survival was evaluated using Kaplan-Meier LogRank test (**H**).

Polθ could be also inactivated by the expression of D2230A+Y2231A polymerase-deficient mutant which exerts a dominant-negative effect (Yousefzadeh et al., 2014). We found that clonogenic activity of Lin^-^cKit^+^ GFP^+^;*Flt3^ITD+/-^;Dnmt3a^R882H/+^*mBMCs transduced with Polθ(D2230A+Y2231A) mutant was decreased when compared to these transduced with wild-type Polθ (Figure 2D). This effect was accompanied by accumulation of DSBs (γH2AX) in the former cells (Figure 2E). In addition, reduction of clonogenic activity by Polθ(D2230A+Y2231A) mutant was also detected in RFP^+^;FLT3(ITD);*Dnmt3a^KD^* 32Dcl3 cells (Figure 2F). When injected into SCID mice, Polθ(D2230A+Y2231A) mutant abrogated the engraftment and leukemogenic capability of RFP^+^;FLT3(ITD);*Dnmt3a^KD^*32Dcl3 cells (Figure 2G, H).

Altogether, we present comprehensive evidence in orthogonal but complementary systems that inactivation of Polθ is selectively detrimental to DNMT3A-deficient myeloid hematological malignancies. This effect is associated with accumulation of formaldehyde-induced DPCs-triggered toxic DSBs and activation of cGAS/STING pathway.

### Polθ inhibitors combined with standard drugs eliminate *DNMT3A*-mutated AML

Since Polθ plays a key role in DNMT3A deficient leukemias, we tested the potential therapeutic value of targeting Polθ. We reported that the repair of etoposide-induced DNA-protein crosslinks (DPCs) is promoted by Polθ (Chandramouly et al., 2021). We show here that the combination of sub-optimal concentrations of etoposide and Polθi ART558, ART812 or RP-6685 eradicated FLT3(ITD);*Dnmt3a^KD^* 32Dcl3 clonogenic cells *in vitro* (Figure 3A, Supplemental Figure S3A). In concordance, the combination of RP-6685 + etoposide was particularly effective in facilitating the accumulation of DPCs and toxic DSBs in FLT3(ITD);*Dnmt3a^KD^* 32Dcl3 leukemia cells (Figure 3B and C, respectively).

**Figure 3.**
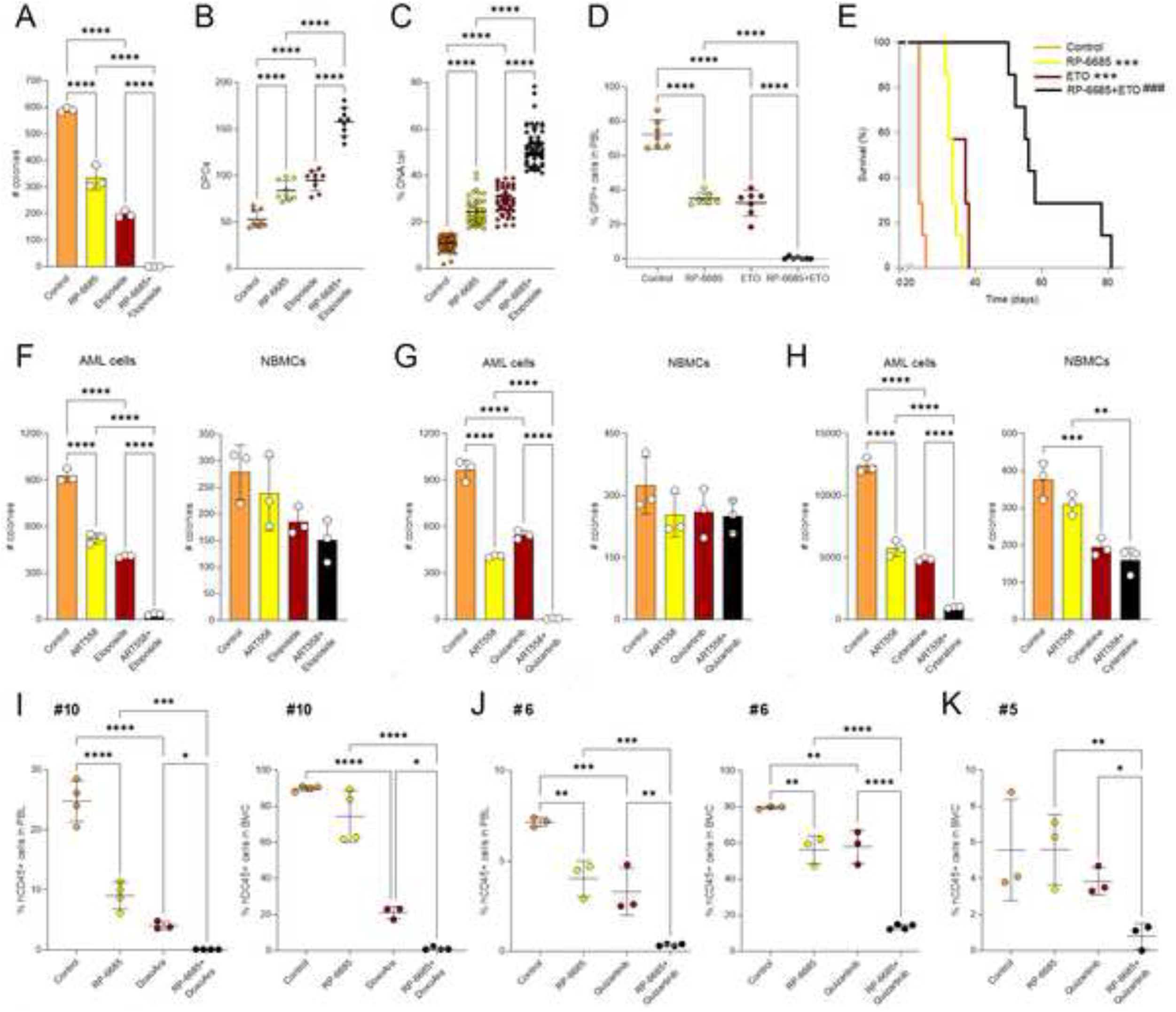
Polθi enhanced the sensitivity of DNMT3A deficient leukemia cells to standard drugs. (**A-C**) Sensitivity of FLT3(ITD);*Dnmt3a^KD^* 32Dcl3 cells to 4μM Polθi (RP-6685) ± 60 nM etoposide was tested *in vitro*. (**A**) Mean number of colonies. (**B**) Detection of DPCs. (**C**) % of DNA in the tail. (**D**) Detection of GFP^+^ FLT3(ITD);*Dnmt3a^KD^* 32Dcl3 cells (FD cells) in PBL of SCID mice untreated (Control) and treated with RP-6685, etoposide (ETO) and the combination (RP-6685+ETO). (**E**) Survival curves and MST of the mice. (**F-H**) Mean number of colonies from Lin^-^CD34^+^ FLT3(ITD);DNMT3A(R882H) AML primary cells (n=3 patients) and Lin^-^CD34^+^ normal bone marrow cells (NBMCs) (n=3 donors) treated with 25 μM Polθi ART558 combined with (**F**) 60 nM etoposide, (**G**) 2.5 nM quizartinib, and (**H**) 0.2 μM cytarabine. (**I-K**) Detection of hCD45^+^ *DNMT3Amut* AML primary cells in PBL and BMC of the NRGS mice untreated (Control) and treated with RP-6685, doxorubicin + cytarabine (DoxoAra), quizartinib, and indicated combinations. Error bars represent the SD; p values were calculated using one-way Anova; mice survival was evaluated by Kaplan-Meier LogRank test.

We tested the effectiveness of Polθi RP-6685, a potent, selective, and orally bioavailable Polθi (Bubenik et al., 2022) alone and in combination with etoposide against very aggressive GFP+ FLT3(ITD);*Dnmt3a^KD^* 32Dcl3 leukemia cells in NRG mice (MST 23.3± 0.3 days after i.v. injection of 10^3^ cells). In concordance, treatment of the leukemia – bearing mice with RP-6685 or etoposide diminished the percentage of GFP+ leukemia cells in peripheral blood by >2-fold. Remarkably, the combination of RP-6685 + etoposide reduced GFP+ cells below detectable levels in 6/7 mice (Figure 3D). The overall survival of GFP+ FLT3(ITD);*Dnmt3a^KD^* 32Dcl3 leukemic mice treated with individual RP-6685 or etoposide was prolonged to 34.5±1.2 and 32.6±0.7 days, respectively whereas the combination of RP-6685 + etoposide extended the survival to 58.6±5.0 days (Figure 3E). Despite elimination of leukemia cells from the peripheral blood shortly after the treatment, the latter mice succumbed to the disease perhaps due to persistence of leukemia cells in other organs, such as bone marrow (Bolandi et al., 2021).

Finally, we examined the effect of Polθi ART558 and/or etoposide against Lin^-^CD34^+^ FLT3(ITD);DNMT3A(R882H) AML primary cells characterized before (Maifrede et al., 2021; Toma et al., 2024; Toma et al., 2025). The combination of ART558 and etoposide exerted >12 times stronger effect against Lin^-^CD34^+^ FLT3(ITD);DNMT3A(R882H) AML cells when compared to individual treatments whereas Lin^-^CD34^+^ cells from healthy donors were only modestly affected (Figure 3F).

Inhibition of Polθ was synthetic lethal in cells with *BRCA1/2* mutations (Ceccaldi et al., 2015; Mateos-Gomez et al., 2015). Since FLT3 kinase inhibitor (FLT3i) induced acute BRCA1/2 deficiency in FLT3(ITD)-positive AML cells (Maifrede et al., 2018), we hypothesized that it would also sensitize FLT3(ITD)-positive DNMT3A deficient cells to Polθ inhibitors. Thus, the combinations of FLT3i quizartinib and Polθi ART558 or ART812 were tested against Lin^-^CD34^+^ FLT3(ITD);DNMT3A(R882H) AML primary cells and FLT3(ITD);*Dnmt3a^KD^* 32Dcl3 cells. We show that quizartinib when combined with ART558 or ART812 eradicated Lin^-^CD34^+^ FLT3(ITD);DNMT3A(R882H) AML primary cells and FLT3(ITD);*Dnmt3a^KD^* 32Dcl3 clonogenic cells *in vitro* (Figure 3G, Supplemental Figure S3B); again Lin^-^CD34^+^ human BMCs from healthy donors were not significantly affected by the treatment.

Moreover, *DNMT3A* mutations sensitized AML cells to a replication stalling agent cytarabine (Venugopal et al., 2022). Suboptimal concentrations of cytarabine combined with ART558 exerted 5x and 3x more potent effect against Lin^-^CD34^+^ FLT3(ITD);DNMT3A(R882H) AML cells and FLT3(ITD);*Dnmt3a^KD^* 32Dcl3 cells, respectively, when compared to individual inhibitors. Meanwhile, Lin^-^CD34^+^ human BMCs from healthy donors were modestly affected by the treatment (Figure 3H, Supplemental Figure S3C).

In addition, DNMT3A(R882H) AML cells were selectively susceptible to the hypomethylating agent azacytidine due to enhanced endogenous cell autonomous viral mimicry response (Scheller et al., 2021). However, azacytidine which selectively suppressed protein translation and induced apoptosis in *DNMT3Amut* cells via viral mimicry response (Scheller et al., 2021) did not substantially enhance the inhibitory effect of ART558 in FLT3(ITD);*Dnmt3a^KD^* 32Dcl3 cells (Supplemental Figure S3D).

Finally, we tested Polθi RP-6685 alone and in combination with standard drugs against *DNMT3A* mutated primary AML xenografts in NRGS mice. For this reason, we combined Polqi with doxorubicin + cytarabine (related to the standard consolidation treatment), etoposide (used to treat relapsed/refractory AML), and with FLT3 kinase inhibitor quizartinib, if FLT3(ITD) was detected. While RP-6685 reduced the number of AML #10 xenograft cells only in peripheral blood, doxorubicin + cytarabine displayed anti-AML effect in the peripheral blood and bone marrow (Figure 3I). Remarkably, the combination of RP-6685 + doxorubicin + cytarabine exerted exceptionally strong anti-AML #10 activity in both peripheral blood and bone marrow. RP-6685 and quizartinib, when used individually, diminished the number of FLT3(ITD)-positive AML #6 xenograft cells in peripheral blood and bone marrow, although the effect in blood seemed more pronounced than in bone marrow (Figure 3J). The combination of RP-6685 + quizartinib was highly efficient in eliminating AML cells from the peripheral blood and bone marrow. Low numbers of FLT3(ITD)-positive AML #5 xenograft cells were detected only in the bone marrow, thus mimicking remission/minimal residual disease (Figure 3K). RP-6685 and quizartinib were not particularly effective as individual agents, but the combination of RP-6685 + quizartinib significantly reduced the population of xenograft cells in the bone marrow.

Toxicology studies revealed that these treatments did not induce prolonged side effects. Body weight was not significantly affected (Supplemental Figure S4A). The analysis of peripheral blood parameters and other organs performed 2 and 17 days after the end of treatment did not reveal any significant toxic effects in mice treated with RP-6685 + quizartinib (Supplemental Figure S4B). On the other hand, the number of white blood cells, neutrophils and/or lymphocytes was diminished on day 2 in mice treated with RP-6685 + doxorubicin + cytarabine and RP-6685 + etoposide, but these parameters fully recovered on day 17. Moreover, no obvious toxic effects of the treatments were detected in other tissues analyzed on day 17 (Supplemental Figure S4C).

Collectively, we showed that Polθi might exert stronger anti-AML effect in peripheral blood than in bone marrow when administered as a single agent, but it was exceptionally efficient in eradicating AML cells in blood and bone marrow when used in combination with standard drugs. Therefore, combination of Polθi and standard treatments should be considered as novel therapeutic approach in DNMT3A mutated AML.

### Inactivation of DNMT3A enhances Polθ expression by downregulation of PARP1-mediated PARylation-dependent ubiquitination (PARdU) of Polθ by UBE2O E3 ligase

Next, we investigated the mechanistic aspects of the dominant role of Polθ in DNMT3Amut leukemia. Western blot analyses revealed abundant overexpression of Polθ protein in FLT3(ITD);*Dnmt3a^KD^* 32Dcl3 cells, and in Lin^-^ FLT3(ITD);*Dnmt3a^-/-^* and *Flt3^ITD+/-^;Dnmt3a^R882H/+^*mBMCs when compared to *Dnmt3a* wild-type counterparts (Figure 4A). Of note, Polθ was almost undetectable in FLT3(ITD);*Tet2^KD^*32Dcl3 cells and *Flt3^ITD/ITD^;Tet2^-/-^* mBMCs, explaining their resistance to Polθi (Figure 1C, D). In addition, incubation of FLT3(ITD) 32Dcl3 cells with DNMT3A inhibitor elevated Polθ expression (Figure 4B). Polθ was also elevated in DNMT3A-deficient cells expressing JAK2(V617F) and MPL(W515L) oncogenic tyrosine kinases (Supplemental Figure S5A) and in HCC-15 and A549 lung carcinoma cells treater for 6 days with DNMT3Ai (Supplemental Figure S5B), implicating a more general role of DNMT3A in regulating Polθ expression in hematological malignancies and solid tumors.

**Figure 4.**
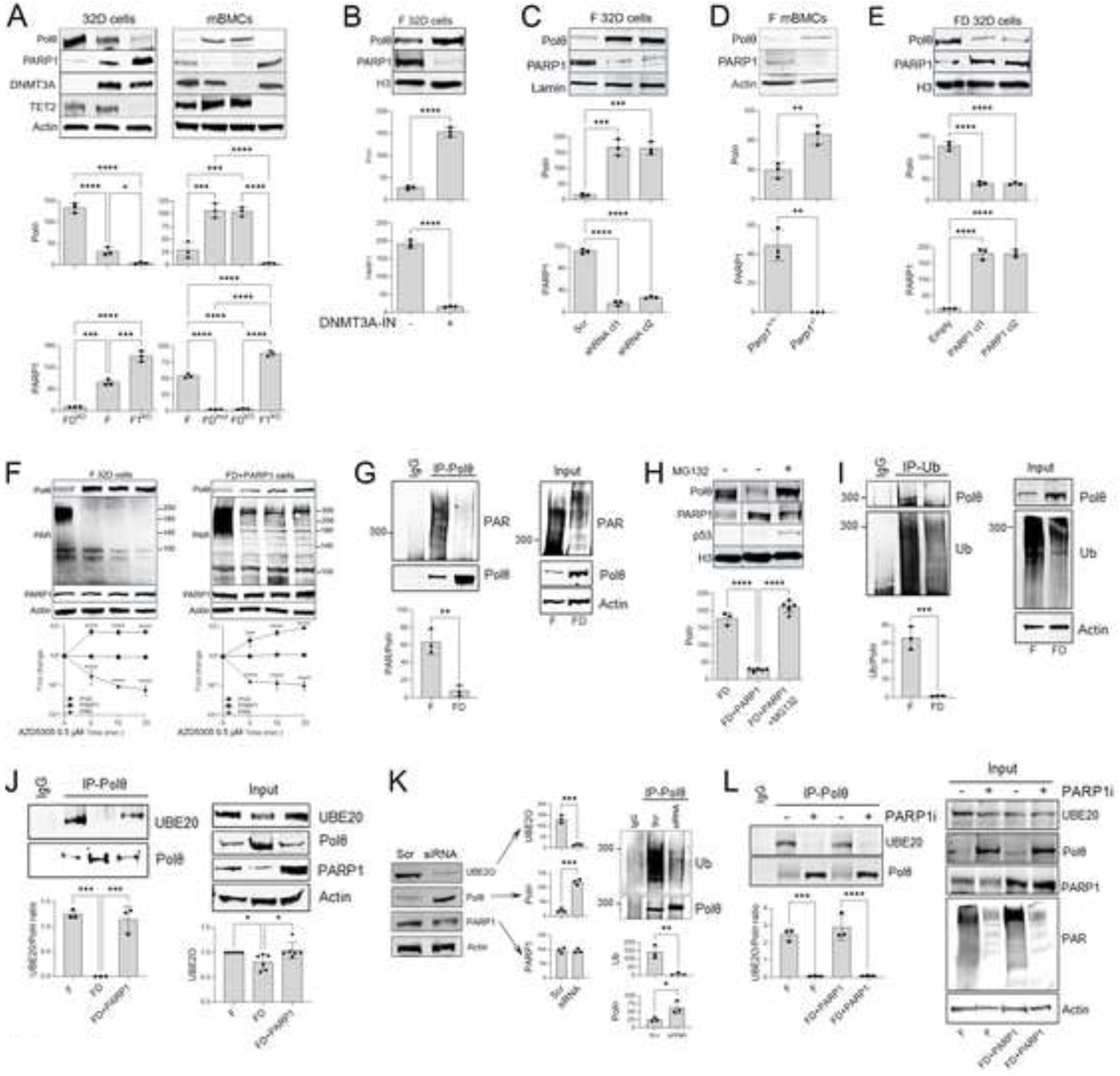
PARylation-dependent UBE2O-mediated ubiquitination of Polθ to elevate Polθ expression in leukemia cells. (**A-E**) Representative Western blots of the indicated proteins in: (**A**) FLT3(ITD) (F), FLT3(ITD);*Tet2^KD^* (FT) and FLT3(ITD);*Dnmt3a^KD^* (FD) 32Dcl3 cells and in Lin^−^ mBMCs from *Flt3^ITD/ITD^*(F), *Flt3^ITD+/-^; Dnmt3a^R882H/+^* (FD^mut^), *Flt3^ITD/ITD^;Dnmt3a^-/-^* (FD^KO^) and FLT3(ITD);*Tet2^KD^* (FT^KO^) mBMCs; (**B**) F 32Dcl3 cells treated (+) or not (-) with DNMT3A-IN-1; (**C**) F 32Dcl3 cells transduced with scrambled or *Parp1* specific shRNA; (**D**) *FLT3^ITD^;Parp1^-/-^* and *FLT3^ITD^;Parp1^+/+^* mBMCs; (**E**) FD 32Dcl3 cells and the clones transduced with PARP1 cDNA expression construct (FD-PARP1 cl1 and cl2). (**F**) Western analysis of F and FD-PARP1 32Dcl3 cells treated with 0.5 μM PARP1i AZD5305 for 0, 5 10 and 20 min. (**G**) Detection of PARylated Polθ in anti-Polθ immunoprecipitates (IP-Polθ) and total cell lysates (Input) from F and FD 32Dcl3 cells. (**H**) Western analysis of F and FD-PARP1 32Dcl3 cells treated (+) or not (-) with proteasome inhibitor MG132. (**I**) Detection of Polθ in anti-Ub immunoprecipitates (IP-Ub) and total cell lysates (Input) from F and FD 32Dcl3 cells. (**J**) Co-immunoprecipitation of Polθ and UBE2O in anti-Polθ immunoprecipitates (IP-Polθ) from F, FD and FD+PARP1 32Dcl3 cells; total cell lysates (Input) are shown, too. (**K**) Detection of Polq, PARP1 and UBE2O in total cell lysates (Input) and Ub-Polq and Polq in anti-Polq immunoprecipitates (IP-Polq) from FD+PARP1 32Dcl3 cells transduced with scrambled or *Ube2o* siRNA. (**L**) Co-immunoprecipitation of Polθ and UBE2O in anti-Polθ immunoprecipitates (IP-Polθ) from F and FD+PARP1 32Dcl3 cells treated (+) or not (-) with 0.5 μM AZD5305 for 60 min; total cell lysates (Input) are shown, too. Mean ± SD and statistical significance of the intensity of specific proteins/protein posttranslational modifications are shown; p values were determined using one-way Anova (**A**, **C, E, F, H, J**) and Student’s t test (**B, D, G, I, K, L**).

We did not detect any significant differences in *Polq* promoter region methylation between FLT3(ITD), FLT3(ITD);*Dnmt3a^KD^*, and FLT3(ITD);*Tet2^KD^* 32Dcl3 cells (Supplemental Figure S5C). In addition, DNMT3A-deficient leukemia cells displayed a modest 20-35% enhancement of *Polq* mRNA expression measured by real-time PCR (Supplemental Figure S5D) which was associated with increased *Polq* mRNA stability detected by analyzing mRNA half-life following transcription inhibition by actinomycin D (Supplemental Figure S5E), but not with enhanced transactivation of the reporter plasmid carrying the fragment of *Polq* promoter and/or alternative splicing determined by analyzing the size of RT-PCR-amplified *Polq* fragments (Supplemental Figure S5F and G, respectively). In addition, 30-40% enhancement of Polθ protein translation detected by *Polq* 3’UTR-mediated stimulation of luciferase activity was also detected in DNMT3A-deficient leukemia cells (Supplemental Figure S5H). These rather modest increases of the mRNA stability and protein translation were unlikely to explain 4-5-fold and a remarkable 10-34-fold overexpression of Polθ protein in DNMT3A deficient leukemia cells when compared to DNMT3A proficient and to TET2 deficient counterparts, respectively (Figure 4A, Supplemental Figure S5I).

Intriguingly, PARP1 expression was inversely proportional to Polθ levels in FLT3(ITD), JAK2(V617F) or MPL(W515L)-positive DNMT3A-deficient cells when compared to DNMT3A proficient and to TET2 deficient counterparts (Figure 4A, B, Supplemental Figure S5A), suggesting that PARP1 might repress Polθ expression. To test this hypothesis, PARP1 was downregulated by shRNA in FLT3(ITD)-positive 32Dcl3 cells, which caused upregulation of Polθ (Figure 4C). Moreover, *FLT3^ITD^;Parp1^-/-^* mBMCs displayed higher expression of Polq when compared to *FLT3^ITD^;Parp1^+/+^* counterparts (Figure 4D). Conversely, ectopic overexpression of PARP1 in FLT3(ITD);*Dnmt3a^KD^* 32Dcl3 (FLT3(ITD);*Dnmt3a^KD^* + PARP1) cells resulted in downregulation of Polq (Figure 4E).

Downregulation of PARP1 in FLT3(ITD);*Dnmt3a^KD^*32Dcl3 cells likely depended on decreased PARP1 translation detected using *Parp1* 3’UTR luciferase assay (Supplemental Figure S6A) and reduced protein stability suggested by cycloheximide chase assay (Supplemental Figure S6B) which was associated with proteasomal degradation (Supplemental Figure S6C). Among E3 ligases reported to be involved in PARP1 degradation (De Vos et al., 2014; Garzon et al., 2022; Gatti et al., 2020; Zhang et al., 2020b), HUWE1 was selectively upregulated in FLT3(ITD);*Dnmt3a^KD^* 32Dcl3 cells when compared to FLT3(ITD);*Tet2^KD^* and FLT3(ITD) counterparts (Supplemental Figure 6D). DNMT3Ai elevated the expression of HUWE1 in FLT3(ITD) 32Dcl3 cells suggesting that overexpression of HUWE1 in FLT3(ITD);*Dnmt3a^KD^*32Dcl3 cells depended on the lack of DNMT3A enzymatic activity (Supplemental Figure S6E). However, we could not detect any differences in *Huwe1* promoter region methylation between FLT3(ITD), FLT3(ITD);*Dnmt3a^KD^*, and FLT3(ITD);*Tet2^KD^* 32Dcl3 cells (Supplemental Figure S6F) thus implicating an indirect effect. HUWE1 co-immunoprecipitated with PARP1 (Supplemental Figure 6G) and shRNA-mediated downregulation of HUWE1 rescued PARP1 and abrogated Polθ proteins expression in FLT3(ITD);*Dnmt3a^KD^* 32Dcl3 cells (Supplemental Figure 6H). At the same time *PARP1* methylation, mRNA expression and stability were not affected by DNMT3A deficiency (Supplemental Figure S6I-K).

Downregulation of PARP1 in FLT3(ITD) 32Dcl3 cells did not increase *Polq* mRNA stability and Polθ translation (Supplemental Figure S6L and M, respectively). In concordance, *in silico* analysis did not detect PARP1 binding sites in any region of *POLQ* exon, intron, promoter, or 3’UTR sites, using the published ChIP and ATAC-Seq datasets deposited in the GeneHancer database (https://academic.oup.com/database/article/doi/10.1093/database/bax028/3737828). Thus, Polθ appeared to be regulated by another PARP1-dependent mechanism.

We reported that Polq is directly PARylated by PARP1 (Vekariya et al., 2024). PARP1-mediated downregulation of Polθ is a rapid and efficient process which depends on PARP1 catalytic activity, because PARP1 specific inhibitor enhanced the levels of Polθ in FLT3(ITD) and FLT3(ITD);*Dnmt3a^KD^* + PARP1 32Dcl3 cells in less than 5 min (Figure 4F), suggesting that PARylated Polq is targeted for degradation. In support of this hypothesis, PARylation of Polq immunoprecipitated from FLT3(ITD);*Dnmt3a^KD^*32Dcl3 cells was strongly reduced when compared to that immunoprecipitated from FLT3(ITD) counterparts (Figure 4G).

The speed and robustness of PARP1-mediated repression of Polθ expression implicated proteasomal degradation (Coll-Martínez and Crosas, 2019). In concordance, while Polθ was downregulated in FLT3(ITD);*Dnmt3a^KD^* + PARP1 32Dcl3 cells, proteasome inhibitor increased the expression of TP53 [positive control, (Lopes et al., 1997)] and rescued the overexpression of Polθ (Figure 4H). Moreover, Polθ was found in anti-Ub immunoprecipitates from FLT3(ITD) cells, but not from FLT3(ITD);*Dnmt3a^KD^*32Dcl3 counterparts (Figure 4I). Altogether, abrogation of PARP1-mediated PARylation of Polθ in FLT3(ITD);*Dnmt3a^KD^*32Dcl3 cells resulted in its decreased ubiquitination and reduced proteasomal degradation.

To identify the E3 ligase responsible for ubiquitination of Polθ, we analyzed a pull-down of Polθ followed by LC-MS/MS and detected UBE2O E3 ligase as a potential partner of Polθ (Vekariya et al., 2024). UBE2O was modestly downregulated in FLT3(ITD);*Dnmt3a^KD^* cells when compared to FLT3(ITD) and FLT3(ITD);*Dnmt3a^KD^* + PARP1 32Dcl3 cells (Figure 4J, Input). Remarkably, UBE2O was not detected in anti-Polθ immunoprecipitates from FLT3(ITD);*Dnmt3a^KD^* cells, while it was readily detectable in those from FLT3(ITD) and FLT3(ITD);*Dnmt3a^KD^* + PARP1 32Dcl3 counterparts (Figure 4J, IP-Polθ). To determine if UBE2O is responsible for PARP1-mediated degradation of Polθ in leukemia cells, expression of UBE2O was knocked down by siRNA in FLT3(ITD);*Dnmt3a^KD^* + PARP1 32Dcl3 cells (Figure 4K). This effect was associated with abrogation of Polq ubiquitination and upregulation of Polθ without affecting PARP1 expression. Moreover, inhibition of PARP1 activity resulted in dissociation of Polθ-UBE2O complex and elevated expression of Polθ (Figure 4L). Altogether, we postulate that DNMT3A deficiency in FLT3(ITD)-positive leukemia cells promoted overexpression of Polθ by abrogation of PARdU-dependent UBE2O E3 ligase-mediated ubiquitination and proteasomal degradation of Polθ.

### Polθ-dependent DNA repair is enhanced in DNMT3A deficient leukemia cells

Next, we examined if abundant overexpression of Polθ in *DNMT3A*mut leukemia cells results in enhanced DNA repair. Intracellular recruitment of Polθ to DNA damage could be detected in the form of nuclear foci (Ceccaldi et al., 2015; Kais et al., 2016; Mateos-Gomez et al., 2015). Polθ foci formation was tested in untreated cells [spontaneous DSBs caused by metabolic products: ROS and formaldehyde, and by replication stress (Chen et al., 2014; Esposito and So, 2014; Nieborowska-Skorska et al., 2017a; Vekariya et al., 2023)] and in cells receiving 2 Gy of γ-irradiation. Immunostaining followed by confocal microscopy revealed that FLT3(ITD);*Dnmt3a^KD^* cells accumulated 5x and 2x more spontaneous and irradiation-induced Polθ foci, respectively, when compared to FLT3(ITD) counterparts (Figure 5A), implicating enhancement of Polθ-mediated DNA repair in the former cells. As expected, Polθ foci were almost undetectable in FLT3(ITD);*Tet2^KD^* cells which express very low levels of Polθ (Figure 4A).

**Figure 5.**
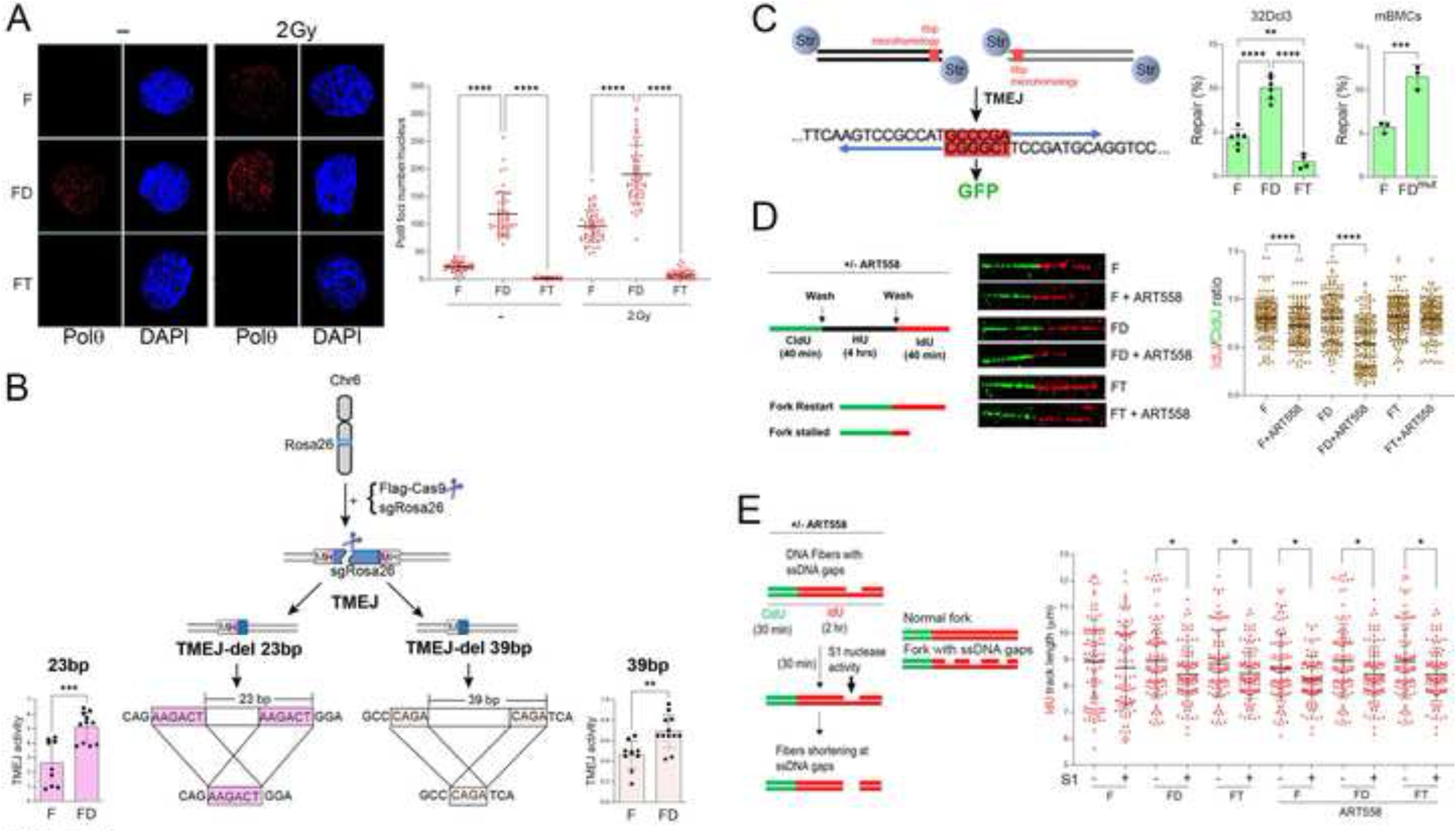
DNMT3A deficiency facilitates Polθ - mediated TMEJ and fork restart, but not ssDNA gap filling, in leukemia cells. (**A**) FLT3(ITD) (F), FLT3(ITD);*Dnmt3a^KD^* (FD) and FLT3(ITD);*Tet2^KD^* (FT) 32Dcl3 cells were irradiated (2Gy) or not. *Left –* representative immunofluorescent staining of Polθ nuclear foci; *Right* – number of foci/nucleus. (**B**) *Central panel*: A diagram illustrating chromosomal TMEJ-del 23bp and TMEJ-del 39bp assays; *Left and right bar panels*: Mean TMEJ-del 23bp and TMEJ-del 39bp activities, respectively. (**C**) *Left panel*: A diagram illustrating DPC-TMEJ (Str = streptavidin); *Right panels*: Mean % of DPC-TMEJ repair in F, FD and FT 32Dcl3 cells and in Lin^−^ mBMCs from *Flt3^ITD/ITD^* (F) and *Flt3^ITD+/-^;Dnmt3a^R882H/+^* (FD) mBMCs. (**D**) *Left panel:* Schematic diagram of replication fork restart. *Middle panel*: Representative DNA fibers from F, FD, and FT 32Dcl3 cells incubated with or without 12.5 μM ART558. *Right panel:* Mean DNA fiber lengths presented as IdU/CldU length ratio from ≥100 DNA fibers. (**E**) *Left panel:* Schematic representation of S1 nuclease treatment to quantify IdU track length. *Right panel:* Mean IdU track length from ≥100 DNA fibers from each F, FD and FT 32Dcl3 cells. Error bars represent the SD; p values were determined using one-way Anova (**A**, **C**-32Dcl3 panel) and Student’s t test (**B, C**-mBMCs panel, **D, E**).

We recently described a two-step spatiotemporal mechanism of Polθ regulation by PARP1 and PARG (Vekariya et al., 2024). First, PARP1 PARylates Polθ to promote liquid-liquid phase separation (LLPS) and facilitates its recruitment to DNA damage sites in an inactivated state. Second, PARG removes repressive PAR marks on Polθ to activate TMEJ. Since LLPS depends on the macromolecule concentration (Alberti et al., 2019), we postulate that the abundance of Polθ expression might alleviate the role of downregulated PARP1 in recruitment of Polθ to DNA damage sites in DNMT3A-deficient leukemia cells. However, it should be noted that reduced PARP1 loading on chromatin (Supplemental Figure S7A) and limited PARylation of Polθ (Figure 4G) in DNMT3A-deficient leukemia cells might provide enough signaling to recruit abundantly expressed Polθ to DNA damage where it is dePARylated and activated by PARG.

Polθ has been reported to play a key role in TMEJ and in restarting of stalled replication forks regardless of HR status, and also in ssDNA gap repair/filling during replication fork progression in HR-deficient cells (Belan et al., 2022; Mann et al., 2022; Schrempf et al., 2022; Wang et al., 2019; Wyatt et al., 2016). We examined these functions using specific experimental protocols to measure key Polθ-mediated DNA repair activities in *DNMT3A*mut leukemia cells.

To assess TMEJ, we took advantage of the intrachromosomal model whereby cells are transfected with Flag-Cas9 and sgRosa26 to generate a DSB in *Rosa26* murine locus on chromosome 6 (Feng et al., 2021). TMEJ uses 6-bp or 4-bp microhomology to generate 23-bp or 39-bp deletions, respectively, detected by quantitative droplet digital PCR assay (Figure 5B central panel, TMEJ-del 23bp and TMEJ-del 39bp). Both TMEJ-del 23bp and TMEJ-del 39bp amplicons of the DSB in *Rosa26* locus were enhanced in FLT3(ITD);*Dnmt3a^KD^* 32Dcl3 cells when compared to FLT3(ITD)-positive counterparts (Figure 5B, peripheral panels).

Moreover, we reported that Polθ-mediated TMEJ plays a key role in repairing DSBs resulting from DPCs (Chandramouly et al., 2021). Thus, DPC-associated DSB repair was measured by a reporter assay carrying a split green fluorescent protein (GFP) construct conjugated with 5′-terminal biotin-streptavidin linkages on both DNA ends (Figure 4C, left panel) (Chandramouly et al., 2021). TMEJ of the left and right 5′-adducted DNA constructs transduced into the cells is expected to use the 6-bp microhomology tract to restore *GFP* sequence resulting in GFP expression. Using this assay, we detected that FLT3(ITD);*Dnmt3a^KD^* 32Dcl3 cells and Lin^-^*Flt3^ITD+/-^;Dnmt3a^R882H/+^* mBMCs displayed >2-fold and >4-fold increased DPC-TMEJ activity when compared to FLT3(ITD)-positive *Dnmt3a* wild-type and FLT3(ITD);*Tet2^KD^* counterparts, respectively (Figure 5C, middle and right panels, respectively).

The impact of Polθ on replication fork restart and/or progression was examined by DNA fiber analysis (Figure 5D, left panel). Cells were labeled with CldU, followed by hydroxyurea (HU) treatment to stall the forks, and subsequently released into IdU in the presence or absence of Polθi ART558. While fork restart and/or progression activity did not differ between untreated FLT3(ITD), FLT3(ITD);*Tet2^KD^* and FLT3(ITD);*Dnmt3a^KD^* cells, ART558 exerted strong (∼1.6x) and modest (∼1.1x) inhibitory effects in FLT3(ITD);*Dnmt3a^KD^* and FLT3(ITD) cells, respectively (Figure 5D, middle and right panels). As expected, ART558 did not affect replication fork restart and/or progression in FLT3(ITD);*Tet2^KD^* cells which express very low levels of Polθ (Figure 4A) and are not sensitive to the inhibitor (Figure 1B).

Polθ-mediated ssDNA gap filling was investigated using modified DNA fiber assay where the gaps were detected by S1 nuclease activity (Figure 5E, left panel). Cells were labeled with the nucleoside analogs CldU and IdU followed by incubation with S1 nuclease to digest regions of ssDNA. S1 nuclease incubation modestly shortened the length of labeled tracks in FLT3(ITD), FLT3(ITD);*Tet2^KD^*and FLT3(ITD);*Dnmt3a^KD^* cells, indicating that ssDNA gaps are not abundant in these cells (Figure 5E, right panel). Importantly, ART558 did not affect the length of labeled tracks suggesting that ssDNA gaps are not processed by Polθ in these cells.

Altogether, we observed that overexpressed Polθ in DNMT3A-deficient cells promoted abundant TMEJ and fork restart, but not ssDNA gap filling.

### DNMT3A knockdown limits DSB end-resection to create favorable substrates for TMEJ

The extent of DNA end resection plays a key role in DSB repair choice and is usually determined by the interplay between the Shieldin complex restricting end resection and DNA nucleases promoting end resection (Setiaputra and Durocher, 2019; Zhao et al., 2020). TMEJ acts on short-range resected ends usually generated by MRE11 and CtIP nucleases, while HR prefers long-range resected ends created by EXO1 nuclease (van de Kamp et al., 2021) (Figure 6A).

**Figure 6.**
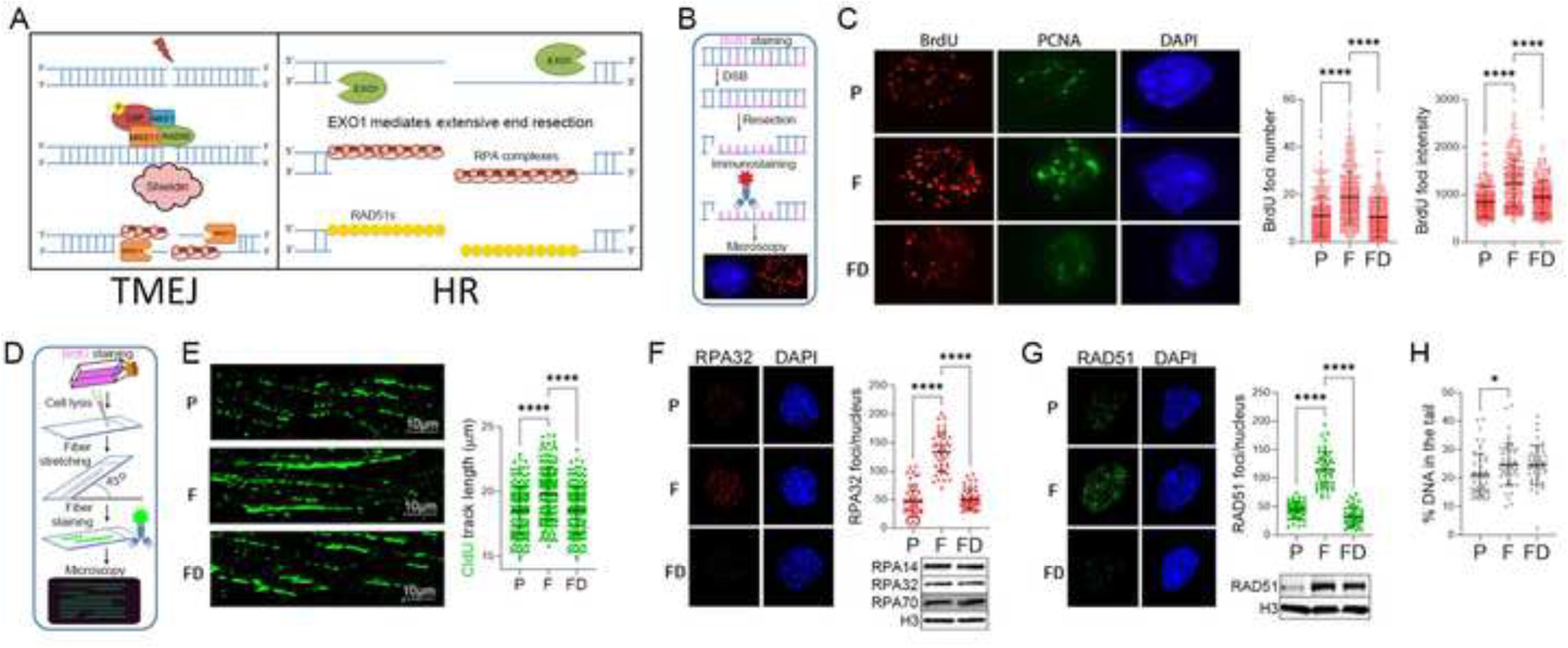
DNMT3A deficiency limited DNA end-resection in leukemia cells. (**A**) The model of DSB end resection favoring TMEJ or HR. (**B-H**) FLT3(ITD) (F), FLT3(ITD);*Dnmt3a^KD^*(FD) and parental (P) 32Dcl3 cells were irradiated (2Gy) and incubated for 2 hrs. (**B**) A scheme of BrdU/PCNA assay. (**C**) *Left panel* - Representative immunofluorescence BrdU and PCNA immunostaining are shown. *Middle and right panels* - mean number of BrdU foci and mean BrdU intensity, respectively, per PCNA-positive nucleus. (**D**) A scheme of SMART assay. (**E**) *Left panel -* representative CldU tracks were shown. *Right panel*: mean of DNA fibers length presented as the CIdU track length. (**F**) *Left panel* - representative RPA32 foci detected by immunofluorescence. *Upper right panel* - mean number of RPA32 foci/nucleus. *Lower right panel* - representative Western blots of the nuclear cell lysates detecting RPA14, RPA32 and RPA70; histone 3 (H3) was a loading control. (**G**) *Left panel –* representative RAD51 foci detected by immunofluorescence. *Upper right panel* - mean number of RAD51 foci/nucleus. *Lower right panel* – representative Western blots of the nuclear cell lysates detecting RAD51; histone 3 (H3) was a loading control. (**H**) Neutral comet assay; mean % of DNA in the tail. Error bars represent the SD; p values were determined using one-way Anova.

To evaluate end resection, we employed BrdU/PCNA assay where following DNA damage and DNA end resection, the length of single-stranded DNA is specifically estimated by an intensity of anti-BrdU immunostaining in the S/G2 phase of the cell cycle marked by PCNA staining (O’Sullivan et al., 2021) (Figure 6B). Using BrdU/PCNA approach, we observed that the presence of FLT3(ITD) promoted DSB-end resection (Figure 6C), consistent with our previous report (Vekariya et al., 2023). Remarkably, *Dnmt3a^KD^*was associated with reduced end resection in FLT3(ITD)-positive cells. In addition, we applied SMART assay, a modified DNA combing technique, to measure DNA resection at the level of individual DNA fibers (Cruz-García et al., 2014) (Figure 6D). In concordance with the cellular BrdU/PCNA assay, SMART assay detected that resection of individual DNA fibers was enhanced in FLT3(ITD) cells when compared to parental counterparts but decreased in FLT3(ITD);*Dnmt3a^KD^*cells (Figure 6E).

Moreover, we evaluated the accumulation of RPA32 into nuclear foci, a well-established resection marker (Sartori et al., 2007). In concordance with BrdU/PCNA and SMART assays, RPA32 foci formation/intensity was decreased in irradiated FLT3(ITD);*Dnmt3a^KD^*32Dcl3 cells when compared to FLT3(ITD) counterparts (Figure 6F). Finally, we detected decreased accumulation of RAD51 foci, a marker of HR (Graeser et al., 2010), in FLT3(ITD);*Dnmt3a^KD^* 32Dcl3 cells when compared to FLT3(ITD) counterparts (Figure 6G). Inhibition of RPA32 and RAD51 foci formation in FLT3(ITD);*Dnmt3a^KD^* 32Dcl3 cells were not due to differences in the protein expressions (Figure 6F and G, insets) and number of DSBs caused by irradiation (Figure 6H). Altogether, we concluded that DNMT3A deficiency restricted DNA end resection in FLT3(ITD)-positive cells.

### DNMT3A knockdown disassembles PARP1-SMARCAD1-MSH2/MSH3 repressive complex to facilitate Polθ loading on DSBs

Limited DNA end resection in FLT3(ITD);*Dnmt3a^KD^* 32Dcl3 cells was associated with decreased association of PARP1 with chromatin (Figure 7A). Using SMART assay, we detected that an extensive DNA end resection in HR-proficient FLT3(ITD) mBMCs was limited in FLT3(ITD);*Parp1^-/-^* counterparts (Figure 7B), implicating PARP1 in this process. PARP1 has been reported to either promote or inhibit DNA end resection by stimulating the recruitment of MRE11 and CtIP nucleases or preventing the recruitment of EXO1 nuclease, respectively, which play a critical role in DSB end processing (Figure 6A) (Caron et al., 2019; Luedeman et al., 2022). However, the amounts of all these proteins were similar in chromatin extracts of FLT3(ITD) and FLT3(ITD);*Dnmt3a^KD^* 32Dcl3 cells (Figure 7A). In addition, chromatin recruitment of REV7 (Xu et al., 2015), one of key factors of the shieldin complex is not affected by DNMT3A knockdown (Figure 7A). Therefore, it is rather unlikely that these factors contribute to limited DNA end resection in DNMT3A deficient cells.

**Figure 7.**
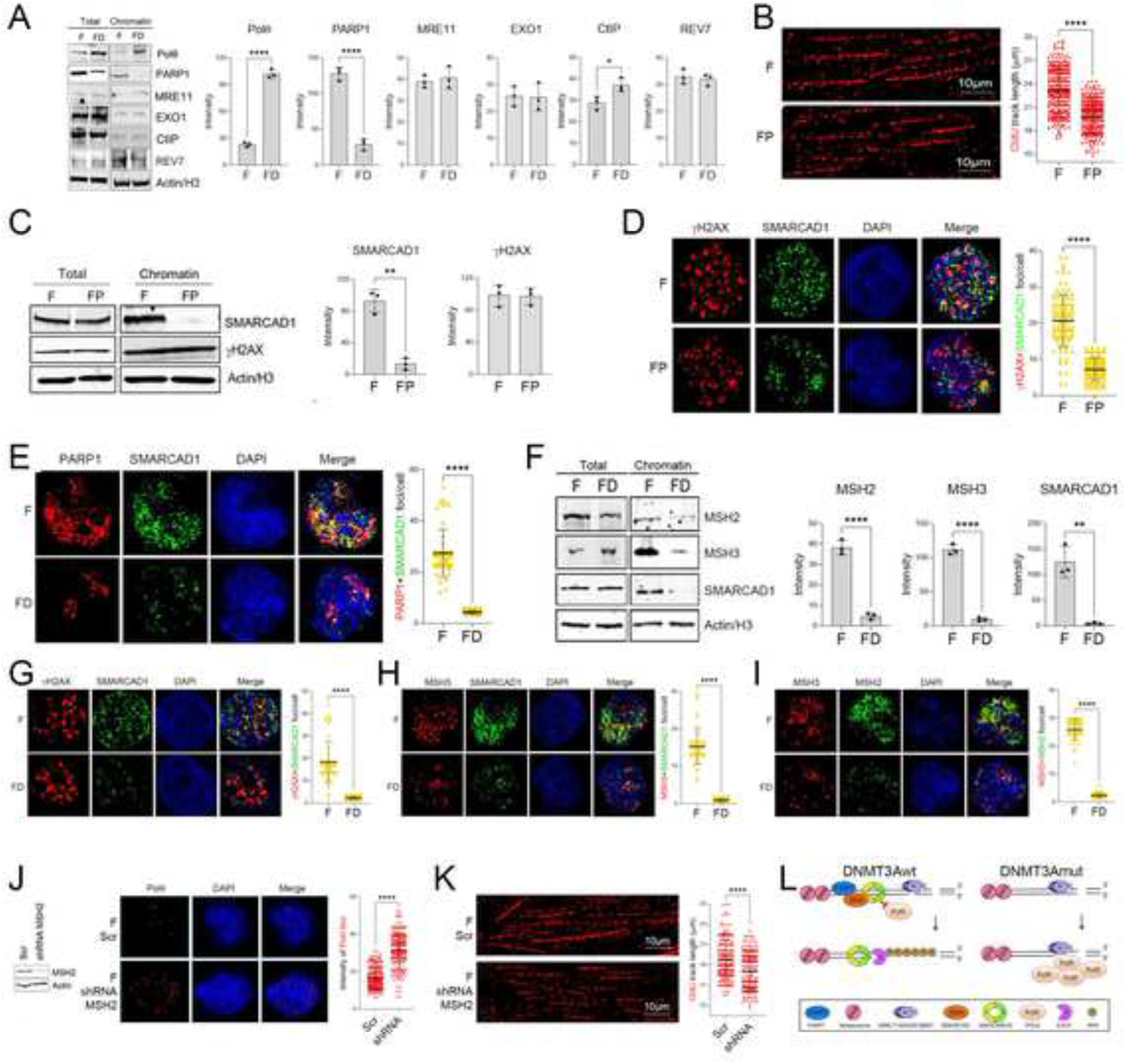
DNMT3A deficiency inhibited PARP1-SMARCAD1-MSH2/MSH3 repressive pathway to facilitate the recruitment of Polθ to DNA. (**A**) FLT3(ITD) (F) and FLT3(ITD);*Dnmt3a^KD^* (FD) 32Dcl3 cells were irradiated (2Gy) and incubated for 2 hrs. before the analyses. *Left panel –* representative Western blots detecting indicated proteins in total cell lysates and chromatin fractions. Actin and histone H3 served as loading controls. *Right panel* - mean of the indicated protein band intensity. (**B-D**) Lin^-^ *Flt3^ITD+/+^* (F) and Lin^-^ *Flt3^ITD+/+^;Parp1^-/-^* (FP) mBMCs were irradiated (2 Gy) 2 hrs. before the analyses. (**B**) SMART assay. *Left panel -* representative CldU tracks were shown. *Right panel* - mean of DNA fibers length presented as the CIdU track length. (**C**) *Left panels –* representative Western blots detecting indicated proteins in total cell lysates and chromatin fractions. Actin and histone H3 served as loading controls. *Right panels* - mean of the indicated protein band intensity in the chromatin fractions. (**D**) *Left panels –* representative SMARCAD1 and/or γH2AX foci; *Right panels* – mean numbers of co-localizing foci/nucleus. (**E-I**) F and FD 32Dcl3 cells were irradiated (2 Gy) 2 hrs. before the analyses. (**E, G-I**) *Left panels –* representative nuclear foci. *Right panels* – mean numbers of co-localizing foci/nucleus. (**F**) *Left panel –* representative Western blots detecting indicated proteins in total cell lysates and chromatin fractions. Actin and histone H3 served as loading controls. *Right panels* - mean band intensity of the indicated protein in the chromatin fractions. (**J**) F 32Dcl3 cells were transfected with the *Msh2* shRNA (F shRNA Msh2) or Scrambled (F Scr). *Left panel* - Western blot showing downregulation of MSH2 in the former cells; *Middle panel* – representative Polθ nuclear foci; *Right panel* – mean numbers of foci/nucleus. (**K**) SMART assay. *Left panels -* representative CldU tracks were shown. *Right panels* - mean length of DNA fibers presented as the CIdU track length. (**L**) A diagram illustrating how *DNMT3Amut* restrains DSB end resection by inhibiting PARP1-dependent repressive role of SMARCAD1-MSH2/MSH3 complex in the recruitment of Polθ to a DSB. Error bars represent the SD; p values were determined using Student’s t test.

PARP1 interacts with the SWI/SNF-like protein SMARCAD1 (Rowbotham et al., 2011) which recruits MSH2/MSH3 heterodimer to prolong end resection, engaging HR rather than TMEJ (Oh et al., 2023b). Thus, we hypothesized that impaired presence of PARP1 on the chromatin in FLT3(ITD);*Dnmt3a^KD^* cells might abrogate the recruitment of repressive SMARCAD1-MSH2/MSH3 complex and facilitate Polθ assembly on DNA damage resulting in limited DNA end resection. To test this hypothesis, SMARCAD1 recruitment to chromatin/DSBs was compared in irradiated FLT3(ITD) and FLT3(ITD);*Parp1^-/-^* mBMCs. The absence of PARP1 strongly decreased the assembly of SMARCAD1 on chromatin and on DSBs (marked by γH2AX) (Figure 7C and D, respectively). Immunofluorescence staining revealed inhibition of colocalization of PARP1 and SMARCAD1 in FLT3(ITD);*Dnmt3a^KD^* 32Dcl3 cells when compared to FLT3(ITD) counterparts (Figure 7E), further supporting the role of PARP1 as an anchor of SMARCAD1 on DSBs in leukemia cells.

Therefore, we tested the role of repressive PARP1-SMARCAD1-MSH2/MSH3 pathway in enhanced assembly of Polθ on DNA damage and limited DNA end resection in FLT3(ITD);*Dnmt3a^KD^* 32Dcl3 cells. Remarkably, while expressions of SMARCAD1, MSH2 and MHS3 proteins were similar in total cell lysates, a severe decrease of chromatin-bound SMARCAD1, MSH2 and MHS3 proteins was detected in irradiated FLT3(ITD);*Dnmt3a^KD^* 32Dcl3 cells when compared to FLT3(ITD) counterparts (Figure 7F). Moreover, immunofluorescent staining showed strong reduction of SMARCAD1 assembly on DSBs (marked by γH2AX foci), and abrogation of SMARCAD1-MSH3 and MSH2/MSH3 colocalization in FLT3(ITD);*Dnmt3a^KD^* cells when compared to FLT3(ITD) counterparts (Figure 7G-I, respectively). In addition, shRNA-induced downregulation of MSH2 in FLT3(ITD) 32Dcl3 cells (Figure 7J, left panel) was also associated with enhanced formation of Polθ nuclear foci (Figure 6J, middle and right panels). In concordance, SMART assay (Figure 6K, left panel) revealed decreased DNA end processing in FLT3(ITD) cells with MSH2 knockdown when compared to FLT3(ITD) counterparts (Figure 7K, right panel).

It has been also reported that SMARCAD1 preferentially associates with H3 citrullinated on arginine 26 (H3R26Cit) (Xiao et al., 2017). However, DNMT3A deficiency was not associated with any changes in H3R26Cit levels (Supplemental Figure S7B), highlighting the role of PARP1, but not H3R26Cit in SMARCAD1 recruitment to DSBs in leukemia cells.

Activation of PLK1 kinase [phosphorylates RHINO1 and Polθ to stimulate TMEJ in mitosis (Gelot et al., 2023)] assessed by phosphorylation on Thr210 (Tavernier et al., 2021) and expression of RHINO [interacts with Polθ enabling its recruitment to DSBs during mitosis (Brambati et al., 2023)] were upregulated by DNMT3A knockdown (Supplemental Figure S7C), and may enhance Polθ recruitment to DNA damage sites in mitosis. However, there were no significant differences in G2/M cell cycle phase and sensitivity to PLK1 inhibitor between FLT3(ITD);*Dnmt3a^KD^* and FLT3(ITD) 32Dcl3 cells (Supplemental Figure S7D and E, respectively), arguing against a major role of PLK1-RHINO axis in Polθ recruitment to DSBs and activation of TMEJ in DNMT3A deficient leukemia cells.

Altogether, we postulate that the inhibition of the assembly of repressive PARP1-SMARCAD1-MSH2/MSH3 complex, but not facilitation of PLK1-RHINO axis or H3R26Cit, is a major stimulator of enhanced recruitment of Polθ to DNA damage sites in FLT3(ITD);*Dnmt3a^KD^* cells (Figure 7L). This effect resulted in limited DNA end resection, stimulation of TMEJ and hypersensitivity to Polθ inhibition.

## Discussion

Accumulation of DNA damage and modulation of DDR are the hallmarks of cancer (Hanahan, 2022). DDR gene alterations are prevalent in many cancer types and led to development of novel therapeutic strategies, e.g., synthetic lethality triggered by PARP1 inhibitors in HR-deficient tumors including leukemia (Esposito and So, 2014; Knijnenburg et al., 2018; Kontandreopoulou et al., 2021; Murai and Pommier, 2023; Nieborowska-Skorska et al., 2017b). In particular, we reported that myeloid hematological malignancies carrying *TET2* mutations displayed HR deficiency and were sensitive to PARP1 inhibitors, whereas these with *DNMT3A* mutations were resistant to these inhibitors (Maifrede et al., 2021). Using a limited drug sensitivity screen, we discovered unexpected vulnerability of *DNMT3A* mutated leukemia cells – their unique dependence on Polθ which could be exploited therapeutically.

For example, Polθi in combination with FLT3 kinase inhibitor quizartinib and topoisomerase II inhibitor etoposide were very efficient in eliminating primary AML cells bearing FLT3(ITD);*DNMT3Amut in vitro* and *in vivo*. Quizartinib most likely hypersensitized FLT3(ITD);*DNMT3Amut* cells to Polθi due to induction of HR and D-NHEJ deficiencies (Maifrede et al., 2018). Similar effect would be expected in JAK2(V617F);*DNMT3Amut* MPN cells treated with JAK1/2 kinase inhibitor ruxolitinib (Nieborowska-Skorska et al., 2017a). On the other hand, the beneficial effect of etoposide and Polθi could depend on accumulation of DPC-mediated toxic DSBs (Chandramouly et al., 2021; Vekariya et al., 2023). Cytarabine also enhanced the effect of Polθi in FLT3(ITD);*DNMT3Amut* cells, probably due to a cumulative effect of cytarabine-mediated fork collapse (Venugopal et al., 2022) and Polθi-mediated delay of replication fork restart (this work). Moreover, the combination of Polθi and 5+3 regimen of doxorubicin+cytarabine was highly effective in eradication of *DNMT3Amut* primary AML xenografts.

However, it should be noted that, in addition to the activating mutations in FLT3 and JAK2 kinases, *DNMT3A* mutations are accompanied by other mutations (Ley et al., 2013; Maslah et al., 2023; Rinke et al., 2020), which might affect the response of leukemia clones to DDR inhibitors, including Polθi (Maifrede et al., 2021; Maifrede et al., 2018; Nieborowska-Skorska et al., 2017a; Nieborowska-Skorska et al., 2019; Toma et al., 2024). Therefore, it is paramount to determine if/how additional clinically relevant mutations may affect the sensitivity of *DNMT3A* mutated leukemia cells to Polθi +/- standard drugs.

The remarkable synthetic viability interdependence of *DNMT3Amut* leukemia cells on Polθ was associated with abundant overexpression of Polθ, which depended on the process called PARylation-dependent ubiquitination (PARdU) (Vivelo et al., 2019). In PARdU the targeted protein is initially PARylated, which triggers an E3 ligase-mediated ubiquitination, resulting in proteasomal degradation of the protein. Here we determined that DNMT3A deficiency repressed PARP1-mediated PARdU of Polθ by the UBE2O E3 ligase, resulting in protection of Polθ from proteasomal degradation. In addition, we reported that FLT3(ITD) kinase-mediated activation of ERK1/2 kinase might protect Polq from c-CBL E3 ligase-mediated ubiquitination and proteasomal degradation (Vekariya et al., 2023).

We also showed that DNMT3A deficiency not only enhanced Polθ expression but also modulated processing of the DNA damage sites to favor Polθ-mediated TMEJ and fork restart in leukemia cells. DNMT3A deficient leukemia cells displayed limited DNA end resection due to dysregulation of the assembly of PARP1-SMARCAD1-MSH2/MSH3 repressive complex, which in turn facilitated Polθ recruitment to DNA damage sites and promoted TMEJ. This conclusion is supported by the reports that the absence of SMARCAD1 promoted TMEJ (Oh et al., 2023b). Moreover, SMARCAD1 deficiency caused frequent fork stalling/inefficient fork restart, which require Polθ to restart the forks [(Lo et al., 2021) and this work]. These two Polθ-mediated activities, TMEJ and replication fork restart, are most likely required to counteract genotoxic effect of excessive formaldehyde-induced DPCs-triggered DSBs and stressed replication forks in DNMT3A deficient leukemia cells (Chandramouly et al., 2021; Mórocz et al., 2017; Stingele et al., 2017; Wang et al., 2019). These Polq-dependent mechanisms do not appear to be directly regulated by DNA methylation. Remarkably, HUWE1-mediated proteasomal degradation of PARP1 in DNMT3A deficient leukemia cells specifically abrogated the assembly of SMARCAD1-MSH2/MSH3 repressive complex and promoted Polq recruitment to chromatin. In support of our findings, DNMT3A deficiency-mediated modulation of specific proteins recruitment to chromatin to regulate the response to daunorubicin was described before (Guryanova et al., 2016).

It has been reported that survival of HR-deficient cells under genotoxic stress depends on Polθ-mediated DNA repair (Ceccaldi et al., 2015). Although DNMT3A-deficient leukemia cells did not display any detectable changes in the expression of HR proteins and in HR activity measured by extrachromosomal reporter cassette (Maifrede et al., 2021), diminished RPA and RAD51 foci formation and limited DNA end resection revealed a potential competition between Polθ-mediated TMEJ and RAD51-mediated HR in these cells. This unexpected observation is supported by the report of crosstalk between HR and Polθ in S phase (Gelot et al., 2023). Polθ recruitment to DSBs to facilitate TMEJ in S phase is reliant on HR intermediates whereby Polθ appears to displace RAD51, a key HR factor, from ssDNA and may also counteract RPA, another HR factor (Ceccaldi et al., 2015; Gelot et al., 2023; Mateos-Gomez et al., 2017; Schaub et al., 2022; Wyatt et al., 2016). Thus, the abundance of Polθ in DNMT3A-deficient leukemia cells might facilitate the eviction of RAD51 and/or RPA from DSBs to promote TMEJ (Mateos-Gomez et al., 2015).

The interplay between downregulated PARP1 and upregulated Polθ appears unique for *DNMT3A* mutated leukemia cells. For example, inactivating mutations in *TET2* gene [encodes methylcytosine dioxygenase 2 that catalyzes the conversion of methylcytosine to 5-hydroxymethylcytosine, (Ley et al., 2013; Maslah et al., 2023; Rinke et al., 2020)] exert an opposite effect on PARP1 and Polθ expression, TMEJ and fork restarting activities, and sensitivity to Polθ inhibition. Polθ was barely detectable in TET2 deficient leukemia cells coinciding with low TMEJ activity, lack of the impact on fork restart and resistance to Polθ inhibitors. Thus, it appears that the sensitivity of leukemia cells to Polθi was associated with the expression levels of Polθ; DNMT3A deficient cells - the highest expressors - were hypersensitive, whereas TET2 deficient cells - the lowest expressors - were resistant. This observation is in concordance with reports implicating high expression of Polθ with sensitivity to the inhibitors (Zatreanu et al., 2021; Zhou et al., 2021).

To date, there are no reported human cancers with defects in the TMEJ, thus *TET2* mutated malignancy is the first one identified with TMEJ deficiency. We reported that *TET2* mutated leukemia cells are also HR deficient, thus the double deficiency in TMEJ and HR explain enhanced sensitivity to PARP and DNA-PKcs inhibitors [(Maifrede et al., 2021) and this work].

Altogether, we postulate that the acquisition of DNMT3A deficiency in leukemia cells triggered the negative feedback loop between PARP1 and Polθ to promote abundant expression of Polθ and restrict DNA end resection, resulting in hyperactive TMEJ, fork restarting and hypersensitivity to Polθ inhibition (Supplemental Figure S8) which may be explored therapeutically. Somatic *DNMT3A* mutations are frequent in myeloid malignancies [for example in 23-37% of cytogenetically normal AML (Chen et al., 2023)], thus the unique dependence of *DNMT3A*-mutated malignancies on Polθ could be exploited therapeutically by applying Polθi alone and in combination with pre-selected standard modalities. However, since DNMT3A deficiency has been also reported in different types of solid tumors (e.g., lung and colon cancers, melanoma), pharmacological targeting of Polθ may have more broad application (AACR, 2017; Kim et al., 2013; Zhang et al., 2020a). This possibility is highlighted by recent clinical trials testing Polθ inhibitors against solid tumors (NCT05687110, NCT05898399, NCT04991480). Since inhibition of Polθ triggered cGAS/STING pathway [(Oh et al., 2023a) and this work], stimulation of immune response may also contribute to anti-tumor effect of Polθi (Zhang et al., 2024).

## Resource Availability

### Lead contact

Further information and requests for resources and reagents should be directed to and will be fulfilled by the lead contact, Tomasz Skorski.

### Materials availability

This study did not generate new unique reagents.

### Data and code availability

- Data reported in this paper will be shared by the llead contact upon request.
- This paper does not report original code.
- Any additional information required to reanalyze the data reported in this paper is available from the lead contact upon request.

## Acknowledgements

This work was supported by R01CA186238, R01CA237286, R01CA244044, R01CA244179, R01CA247707, R01CA283396 and LLS TRP 6628-21 to T.S. G.A.C. was supported by the National Institutes of Health (NIH; HL147978, CA236819 and DK124883), the Edward P. Evans Foundation, the Leukemia and Lymphoma Society (6667-23) and the American Cancer Society (CSCC-RSG-23-991417-01-CSCC). G.P.G. was supported by P01CA247773. G.S.V. and B.J.P.H. were supported by the Bloodwise Program Grant (17006) and Blood Cancer UK Grant (21006). J.Y.M. is a Canada Research Chair in DNA repair and Cancer Therapeutics and was supported by a CIHR Project Grant (PJT-526991). P.P-B. was supported by the grant 2018/30/M/NZ3/00274 from the National Science Center, Poland. We thank Dr. Sara H. Small (Fox Chase Cancer Center and Fels Cancer Institute for Personalized Medicine) for critical reading of the manuscript. We acknowledge Boris Bartholdy from Albert Einstein College of Medicine, Bronx, NY, USA for his involvement in analyzing PARP1 methylation and expression in the published database.

## Authors contribution

BVL, UV, MMT, MN-S, M-CC, MG, ZH, MW, JG, EV-W, PP-B, A-MK, SZ, JA, RN and GC performed investigation and formal analysis. EH and PV performed formal analysis and provided AML patient samples, KP supervised PP-B, RP supervised GC, GSV and BJP supervised MG, AB supervised MW, MW supervised RN, and J-YM supervised M-CC. GPG and GAC contributed to investigation. TS conceptualized the idea, acquired funding, supervised BVL, UV, MMT, MN-S, and ZH, and wrote the manuscript.

## STAR Methods

### Primary human leukemia cells and cell lines

Somatic mutations in Lin-CD34+ AML patient cells obtained at diagnosis are listed in the Key resources table and were described before (Maifrede et al., 2021; Toma et al., 2024; Toma et al., 2025). Lin^-^CD34^+^ cells were obtained from mononuclear fractions by magnetic sorting using the EasySep Lin negative selection followed by CD34 positive selection cocktail as described before (Nieborowska-Skorska et al., 2017b). Human AML primary cells were incubated in StemSpan®SFEM medium (Stem Cell Technologies, Vancouver, Canada) supplemented with the cocktail of growth factors (100 ng/ml SCF, 20 ng/ml IL-3, 100 ng/ml FLT-3 ligand, 20 ng/ml G-CSF, 20 ng/ml IL-6) as described before (Nieborowska-Skorska et al., 2017b). JAK2(V617F)-positive human HEL cells and JAK2(V617F);*DNMT3A*(R882H)-positive human SET-2 cells were described before (Quentmeier et al., 2006; Quentmeier et al., 2020).

### Primary murine cells and cell lines

Lin^-^cKit^+^ murine bone marrow cells were obtained from the following genetically modified mice: (1) *Flt3^ITD^*, *Flt3^ITD^;Tet2^-/-^* and *Flt3^ITD^;Dnmt3a^-/-^*, and *FLT3^ITD^;Parp1^-/-^* which were described before (Maifrede et al., 2021; Vekariya et al., 2023); (2) *Vav-Cre(+);Dnmt3a^(f/f)^;Polq^-/-^* were generated by crossing *Vav-Cre(+);Dnmt3a(f/f)* and *Polq^-/-^*, mice, and (3) *Mx-1-Cre;Dnmt3a^R882H/+^;Flt3^ITD/+^*mice were generated by crossing *Dnmt3a^floxR882H/+^* mouse model (Gozdecka et al., 2025) with *Flt3-ITD* (Lee et al., 2007) and *Mx1-Cre* mice (Kuhn et al., 1995). Cre expression was induced by intraperitoneal injection of 5- to 6-week-old mice with pIpC (Sigma P1530, 20mg/kg; five doses over a period of 10 days). Mouse ear snips were lysed with DirectPCR Lysis Reagent (Viagen, 401-E) according to the manufacturer’s instructions. Lysed DNA was used for genotyping of each allele with the primers listed in the Key resources table. PCR was performed with REDTaq ReadyMix PCR Reaction Mix (Sigma, R2523) with following conditions: initial denaturation at 95°C for 1 minute, followed by 35 cycles of 95 °C for 15 s, annealing at 57°C for 15s and elongation at 72°C for 15s. Final elongation was performed at 72°C for 10 minutes. PCR products were visualized on the 2.5% agarose gel (Supplemental Figure S9). Mice with proper genotypes were aged and sacrificed at signs of disease. Lin^-^cKit^+^ cells were obtained from mononuclear fractions by magnetic sorting using the EasySep Lin negative selection followed by cKit positive selection cocktail as described before (Nieborowska-Skorska et al., 2017b). Murine normal and malignant hematopoietic cells were maintained in StemSpan®SFEM medium supplemented with: (normal cells, and CML-like and AML-like: 100 ng/ml SCF, 20 ng/ml IL-3 and IL-6; MPN-like: 100 ng/ml SCF; 10 ng/ml FLT3 ligand; 20 ng/ml IL-3, IL-6, G-CSF and GM-CSF; 12 units/ml EPO; 2.5 ng/ml TPO] as described before (Maifrede et al., 2018; Nieborowska-Skorska et al., 2017a; Nieborowska-Skorska et al., 2017b). FLT3(ITD) (F), FLT3(ITD);*Tet2*-*KD* (FT-KD) and FLT3(ITD);*Dnmt3a*-*KD* (FD-KD) 32Dcl3 cells were characterized before (Maifrede et al., 2021).

### Expression constructs

pMIGR1-GFP was purchased from Addgene (cat# 27490). pMIG1-FLT3(ITD)-GFP, pMIG1-BCR-ABL1-GFP and pMIG1-JAK2(V617F)-GPF were described before (Maifrede et al., 2018). pCDH-EF1-FHC-POLQ (*POLQ*-wildtype) (#64875) and pCDH-EF1-FHC-POLQ-DY2230AA (*POLQ*-polymerase mutant) (#64878) were purchased from Addgene. *POLQ*-wildtype and *POLQ* polymerase mutant cDNAs were cloned into pCMMP-MCS-IRES-mRFP (Addgene #36972) by Fast Digest *BamH*I (#FD0054), *Not*I (#FD0596) and *Bgl*II (#FD0084) from ThermoFisher Scientific. In this experiment, after enzymatic digestion, DNA fragments were purified by GeneJET Gel Extraction Kit (ThermoFisher Scientific #K0691) and then ligated by Rapid DNA Ligation Kit (ThermoFisher Scientific # K1422). Ligation products were then heat-shock transformed into *E. coli* DH5α. Single bacterial colony which survived in ampicillin agar LB medium was picked up for DNA plasmid extraction by Macherey-Nagel NucleoSpin Plasmid (#74049950). To confirm the sequence of *POLQ*-wildtype and *POLQ*-polymerase mutant, pCMMP-MCS-IRES-mRFP-POLQ and pCMMP-MCS-IRES-mRFP-POLQ-DY2230AA were sequenced by Sanger sequencing.

### Viral infection

10 µg of the expression plasmid and 5 µg pCL-Eco (Addgene plasmid #12371) were co-transfected into Pheonix-Eco cells in the culture medium without antibiotics, to produce the retrovirus packaging these plasmids by Lipofectamin 2000 Transfection Reagent (ThermoFisher Scientific # 11668019). After 24 and 48 hours of the transfection, the retroviral production was obtained and purified by 0.45 µm pore filter. Next, 0.5 x 10^6^ cells were suspended in 1 mL retroviral production, supplemented with 8 µg polybrene. The spin (540 g) infection was performed in 90 minutes at 30°C, and the retroviral production was replaced by regular culture medium after 4 hours. GFP^+^ and RFP^+^ cell populations were obtained by flow cytometry. To produce the lentiviral production for the infection, 10 µg expression plasmid, 7 µg psPAX2 (Addgene cat# 12260), and 5 µg pMD2.G (Addgene #12259) were co-transfected into HEK 293T/17 cells (ATCC # CRL-11268) in the culture medium without antibiotics. The medium was refreshed every 24 hours. The lentiviral production was collected and purified by 0.45 µm pore-size filter after 24 and 48 hours. The spin infection was performed by resuspending 0.5 x 10^6^ cells in 1 mL lentiviral production with 8 µg polybrene supplement, followed by the spin at 2200 rpm/30°C in 60 minutes. After the incubation at 37°C/5% CO_2_ in 4 hours, the lentiviral production was replaced by regular medium. After 48 hours, GFP^+^ cells were sorted by flow cytometry.

### Knockdown of MSH2 and UBE2O

Cells were transfected with 2 μM shRNA of MSH2 (Dharmacon, cat# V3SM11241-234501644) or 200 nM siRNA-smart pool UBE2O [Dharmacon, cat# L-064151-01-0020, (J-064151-09, J-0654151-10, J-065151-11, J-064151-12)] co-transfected with 100 ng pCMMP-MCS-IRES-mRFP using Nucleofector^TM^ kit (cat#. VPA-1003, Lonza, Germany). GFP^+^ (MSH2 Knockdown) or RFP^+^ (UBE2O knockdown) cells were sorted 48 hours post-transfection; downregulation of MSH2 or UBE2O was confirmed by Western blot.

### Leukemogenesis in mice

Female SCID (C.B-17/IcrHsd-*Prkdc*^scid^) mice were purchased from Inotiv (formerly Envigo #182). The animals were sub-lethally total body irradiated (2 Gy) and injected i.v. with (1) 1 x 10^6^ GFP^+^ Lin^-^cKit^+^ GFP^+^ *Dnmt3a-/-* and *Dnmt3a-/-;Polq-/-* murine bone marrow cells (mBMCs) expressing FLT3(ITD), and (2) 1 x 10^3^ RFP^+^ FLT3(ITD);*Dnmt3a*-ko 32Dcl3 cells transduced with *POLQ* wild-type and DY2230AA dominant-negative mutant. GFP^+^ and RFP^+^ cells were detected in PBL by flow cytometry. Survival of the mice injected with 32Dcl3 cells was monitored.

### Inhibitors and cytotoxic drugs

The following DNA repair inhibitors were applied: PARPi olaparib (Selleckchem, cat# S1060), ATMi KU60019 (EMD Millipore, cat# 531978), ATRi VE-821 (Selleckchem, cat# S8007), DNA-PKi II (EMD Millipore, cat# 260961), RAD52i DI-03 (Huang et al., 2016), RAD51i RI-1 (Selleckchem, cat# S8077), CHK1i rabusetib: (Selleckchem, cat# S2626), CHK2i BML-277 (Selleckchem, cat# S8632), Polθi novobiocin (Selleckchem, cat# S2492), Polθi ART558 (MedChemExpress, cat# HY-141520), Polθi ART812 (MedChemExpress, cat# HY-139289), and Polθi RP-6685 (MedChemExpress, cat# HY-151462). Polθi RTx-161 was synthesized as described (Fried et al., 2024). Etoposide (cat# S1225), cytarabine (cat# U-19920A), doxorubicin (S1208), azacytidine (cat# S1782), C-176 (cat# S6575), PLK1i volasertib (cat# S2235), FLT3i quizartinib (cat# S1526) were purchased from Selleckchem. DNMT3A selective inhibitor DNMT3A-IN-1 (cat# HY-144433) was from MedChemExpress (Huang et al., 2021). MG132 (cat# M7449), and hydroxyurea (cat# H-8627) were from Sigma. All compounds were dissolved, aliquoted and stored following manufacturer’s instructions. Compounds were added for 1-3 days, as indicated, following by trypan blue exclusion viability test and/or plating in Methocult (Stemcell Technologies cat# 04320) when indicated; colonies were scored after 5-7 days.

### Detection of proteins in total and nuclear cell lysates and in chromatin fractions

Nuclear and total cell lysates were obtained as described before (Cramer-Morales et al., 2013). Chromatin bound proteins were isolated using a chromatin extraction kit (Abcam cat# ab117152) according to the manufacturer’s instructions. Briefly, 1 x 10^7^ cells were washed 2 times with PBS, followed by addition of 1x lysis buffer (1 x 10^6^ cells/100 μL) containing protease inhibitor cocktail and incubated on ice for 10 min. The samples were vortexed for 10 sec. followed by centrifugation at 2,800 g for 5 min. at 4° C. The supernatants were carefully removed (cytosolic fraction) and pellet were resuspended with 500 μL (1 x 10^6^ cells/50 μL) of working extraction buffer on ice for 10 min. and vortexed occasionally. The resuspended samples were sonicated 2 × 20 sec. The samples were cooled on ice between sonication pulses for 30 sec., and centrifuged at 16,000 g at 4°C for 10 min. The supernatants were transferred to a new tube, and chromatin buffer was added at a ratio of 1:1. The extracts were then used for SDS-PAGE and the proteins were detected by Western blotting.

### Immunoprecipitation

Immunoprecipitation experiments were performed as described before (Vekariya et al., 2019; Vekariya et al., 2018). Briefly, 15 x 10^6^ cells were collected upon completion of incubation. Cells were washed twice with 1X PBS and centrifuge at 3000 g for 5 min. Pellets were subjected to lysis using IP lysis buffer. Lysate were precleared using Dynabeads^TM^ Protein A (cat# 10001D, Invitrogen) and followed by incubation with Anti-HA Magnetic beads (cat# 88837, Pierce) for 16-18 hrs. at 4°C. Next, beads were washed twice with lysis buffer, once with PBS. Purified proteins were eluted SDS sample buffer by incubating for 5 minutes at 95°C. Proteins were resolved on SDS-PAGE and detected by Western blotting.

### Western Blotting

Total cell, nuclear or chromatin protein fractions were mixed with 5x sample loading buffer (Thermo Fisher cat# 39000) to dilute the buffer to 1x. The mixture was then heated at 95-100°C for 5 min. to denature protein and loaded onto 4-12% GenScript SurePAGE™ Precast Gels (GenScript cat# M00653). The electrophoresis was run in Tris-MOPS-SDS running buffer (GenScript cat# M00138) at 200V for 30-40 min. After SDS-PAGE electrophoresis, proteins were transferred into nitrocellulose membrane followed by washing twice for 5 min. with MilliQ water to completely remove transferring buffer and blocking in 5% BSA in PBST for one hour. The membranes were incubated overnight at 4°C with primary antibody diluted 1:1000 in 5% BSA/PBST. The membrane was then washed three times by PBST and incubated with the secondary antibody diluted (1:5000) in 5% BSA/PBST for one hour. The procedure was continued by washing membrane three times with 1x PBST and using Pierce ECL Western Blotting Substrate (Thermo Fisher cat# 32106). The protein bands were visualized by iBright.

### Immunofluorescence (IF) and confocal microscopy

Cells were processed and analyzed for IF as described before (Vekariya et al., 2023). Briefly, cells were fixed with 4% (v/v) paraformaldehyde for 20 min. at 4°C. Then, cells were washed with PBS, permeabilized with 0.5% (v/v) Triton X-100 for 10 min and blocked with PBS containing 3% BSA. Cells were incubated overnight at 4°C with 3% BSA in PBS containing respective primary antibodies followed by washing 3x with PBST buffer and 1 hour incubation with secondary antibodies. After incubation, cells were washed 5x in PBST for 3 min., followed by mounting on a coverslip with 20 μl mounting solution. Cells were visualized and imaged using a Leica SP8 Confocal microscope at a 63X objective magnification, and images were analyzed using ImageJ software. For quantification, > 50 cells were counted from two independent experiments/group. Z-stacking (5 slices per Z-stack with 1 μm) was performed to assess the protein-protein co-localization.

### Protein stability assay

Protein stability was analyzed as described before (Vekariya et al., 2023). Briefly, cells were treated with 20 μg/mL cycloheximide (CHX) (Selleckchem, cat# S7418) for the indicated time points. Total cell lysates were analyzed by SDS-PAGE electrophoresis followed by Western blotting.

### Neutral comet assay

DSBs were detected by neutral comet assay as described before (Maifrede et al., 2018). Briefly, comet assays were performed using the Oxiselect Comet Assay Kit (Cell Biolabs, cat# STA-355) according to the manufacturer’s instructions. Images were acquired by an inverted Olympus IX70 fluorescence microscope using an FITC filter, and the percentage of tail DNA of individual cells was calculated using the OpenComet plugin of ImageJ.

### TMEJ assays

DPC-TMEJ activity was detected as described before (Vekariya et al., 2023). Chromosomal TMEJ activity was measured as described before with modifications (Feng et al., 2021). Briefly, 3 x 10^4^ cells were transfected with 5μg Flag-Cas9, 5μg sgRosa26 and 1μg pEGFP-N2 with a Neon transfection kit (Invitrogen, cat# MPK 10025) using 1400 V, 10 ms pulse. Twenty-four hours after transfection, a portion of cells were analyzed for transfection efficiency, and genomic DNA was extracted from remaining cells. Droplet genomic PCR was performed to detect TMEJ products.

### BrdU/PCNA assay

DNA end resection was examined as described before (O’Sullivan et al., 2021).

### DNA fiber assay measuring replication fork restart/progression

DNA fiber assay performed as described previously with modifications (Bryant et al., 2009). Briefly, cells were treated or not with 12.5 μM ART558 for 24 hrs. to inhibit TMEJ (Vekariya et al., 2023) followed by stepwise incubation with 50 μM CldU (40 min.), 4 mM HU (4 hrs.) and 250 μM IdU (40 min.) in the presence of ART558. Cells were washed 3 x with PBS between each incubation step. Then, cells were harvested, resuspended in 1X PBS at concentration of 1000 cells/μl. 2 μl of cell solution was lysed on slides (Superfrost Plus microscope slides, Fisher Scientific, cat# 12-550-15) with 200 mM Tris–HCl, 50 mM EDTA, 0.5% SDS, pH 7.4 buffer for 8 min. Slides were tilted to a 15-degree angle to spread fibers followed by air drying and fixing in 3:1 methanol: acetic acid for 10 min and 1 wash with H_2_O followed by air dry. Then, slides were stored at -20° C until further use. Next, denaturation of DNA fibers was performed in 2.5 M HCl for 2.5 hrs. followed by two times washing with PBS for 5 min. before and after denaturation. Slides were then incubated with BSA solution (2% in PBS) for 40 min. at room temperature prior to incubation with primary antibodies recognizing CldU (Abcam, cat# ab6326, dilution 1:300) and IdU (BD Biosciences, cat# 347580, dilution 1:100) for 2.5 h at room temperature, 3 times washed, then incubated with AlexaFluor488 and AlexaFluor594 conjugated secondary antibodies (ThermoFisher Scientific, cat# A11062 and A21470, respectively, dilutions 1:300) for 1 hour at room temperature. Fibers were mounted after 3 washes with PBS. Images were acquired with a Leica SP8 LSM Confocal microscope and fiber lengths were measured using ImageJ software. A minimum of 100 replication forks were measured per group and DNA fiber lengths are presented as IdU/CldU length ratio.

### DNA fiber assay measuring ssDNA gaps

S1 nuclease assay was performed as described previously (Panzarino et al., 2021). Briefly, cells were treated or not with 12.5 μM ART558 for 24 hrs. to inhibit TMEJ (Vekariya et al., 2023) and incubated with 50 μM CIdU for 40 min to label replication forks, followed by 250 μM ldU 2 hrs. in the presence or absence of ART558. Subsequently, cells were permeabilized with CSK buffer (100 mM NaCl, 10 mM MOPS, 3 mM MgCl2, pH 7.2, 300 mM sucrose, and 0.5% Triton X-100) at room temperature for 8 minutes, followed by S1 nuclease treatment (20U/μL) in S1 buffer (30 mM sodium acetate, pH 4.6, 10 mM zinc acetate, 5% glycerol, and 50 mM NaCl) for 30 min. at 37^0^C. Finally, cells were collected, pelleted, and resuspended in 100–500 μL PBS at concentration 1-2 x 10^3^; 2 μl of cell suspension was spotted on a positively charged slide and lysed and processed as described in the DNA fiber assay section above.

### Single-Molecule Analysis of Resection Tracks (SMART)

SMART assay were performed as described previously (Altieri et al., 2020). Briefly, cells were treated with 100 μM CldU for 24 hrs followed by Irradiated at 3 Gy and incubated for 3 hrs. Cells were collected and washed with ice-cold PBS. Resuspend cells at concentration of 1 x 10^3^ cells/μL. 2 x 10^3^ cells were lysed on slides directly (Superfrost Plus microscope slides, Fisher Scientific, cat# 12-550-15) with 7.5 μL of lysis buffer (200 mM Tris–HCl, 50 mM EDTA, 0.5% SDS, pH 7.4). Slides were tilted 15-degree to spread fibers followed by air dried and fixed in 3:1 methanol: acetic acid for 10 min. Slides were dipped in H_2_O and air dried followed by store at - 20^0^ C overnight. Next day slides were washed with PBS for 5 min. and blocked with 2% bovine serum albumin (BSA) in PBS for 40 min at room temperature prior to incubation with primary antibodies recognizing CldU (Abcam, catalog# ab6326, dilution 1:300) for 2.5 hrs at room temperature. Slides washed with PBS three times, then incubated with AlexaFluor488 conjugated secondary antibodies (ThermoFisher Scientific, cat# A11062, dilutions 1:300) for 1 hr. at room temperature. Images were acquired with a Leica SP8 LSM Confocal microscope and fiber lengths measured using ImageJ software.

### DNA dot blot

This method was employed to detect the reduction in the formation of 5-methylcytosine (5-mC) after using DNMT3A inhibitor in the cells, and it was conducted as described before (Maifrede et al., 2021). Briefly, cells were treated with DNMT3A-IN-1 (MedChemExpress cat# HY-144433) for 6 days, followed by genomic DNA (gDNA) extraction. Then, gDNA samples were further diluted to 60 ng/µL. gDNA (2.0, 1.5, 1.0, and 0.5 µg) was prepared in total volume of 100 μL of nuclease-free water. Then, 100 μL of 2x DNA denature buffer was added into gDNA, before the denaturation step at 96-100^0^C for 10 minutes. The denaturation was terminated by adding 200 μL of ice-cold 2M ammonium acetate (pH 7), and chilled on ice for 5 minutes. The entire DNA mixture was loaded through BioRad Dot Blot apparatus until all the wells were completely vacuumed. The membrane was briefly washed in 2x Saline Sodium Citrate Buffer, and DNA-membrane crosslink was performed under UV on 2 minutes. The membrane was then blocked in 5% non-fat dry milk in PBST for 1 hour and incubated overnight with anti-5-mC primary antibody (Cell Signaling, cat# 28692) diluted (1:1000) in 5% BSA in PBST at 4^0^ C. The following day, the membrane was washed in PBST and incubated for 1 hr. with HRP-conjugated antibody diluted 1:5000 in 5% BSA in PBST before adding the Pierce ECL Western Blotting Substrate to visualize the dot blot using iBright.

### *Polq* RT-PCR

Total RNA was extracted from 10^7^ cells. The cells were lysed in 600 µL TRI-Reagent (MRC cat# TR-118) to obtain RNA, DNA and proteins. Then, the equal volume of 100% ethanol was added to precipitate RNA and DNA. Genomic DNA was removed, and total RNA was washed and purified by Zymo Research R2050 Direct-zol RNA MiniPrep (Genesee cat# 11-330). The purification and concentration of RNA were measured by NanoDrop Spectrophotometer (Thermo Scientific cat# ND-1000). Genomic DNA removal was validated by agarose (1% w/v) electrophoresis. Next, 3 µg total RNA was employed to synthesize cDNA by SuperScript™ IV First-Strand Synthesis System (Thermo Scientific cat# 18091050). Basically, RNA was annealed to Oligo d(T)_20_ primer, and cDNA was synthesized by reverse transcriptase enzyme. In the last step, the remaining RNA was removed by *E. coli* RNase H. In the PCR reaction, 1.5 µL of cDNA template was added to 2X DreamTaq Green PCR Master Mix (Thermo Scientific cat# K1082), 1 µM forward primer, 1 µM reverse primer, and nuclease-free water to obtain the total reaction volume up to 25 µL. Primers are listed in the Key resources table. Nuclease-free water was used to replace cDNA template as the negative control. Thermocycler included initial denaturation at 95^0^C in 3 min.; followed by 36 cycles of 95^0^C in 1 min., 58^0^C in 45 sec., and 72^0^C in 2 min.; and the final extension at 72^0^C in 10 min. Finally, PCR products were evaluated by agarose (1% w/v) electrophoresis.

### CpG bisulfite amplicon sequencing of murine *Polq*

Genomic DNA (250 to 500 ng) of the FD and FT cells was converted by sodium bisulfite using the EZ DNA Methylation-Lightning kit (Zymo Research) according to the manufacturer’s instructions. PCR was used to amplify the converted DNA using the following primer pairs: 5′-TTTAGTTAGGAGAATTTTGTGGTTT-3′ plus 5′-AATAACAACCCTACCTCCAACTATC-3′. PCR products were sent to the Massachusetts General Hospital Center for Computational and Integrative Biology DNA Core Facility for amplicon sequencing. After trimming adapter sequencing, the sequences of interest were aligned to the converted mm39 genome using Bowtie2 and Bismark; bedgraph files were then extracted to graph, in Integrative Genomics Viewer, the percent methylation corresponding to each of the CpG locations.

### Promoter transactivation assay

The cells were cultured in Iscove-modified Dulbecco medium growth medium supplemented with 10% FBS, 1% L-glutamine, 1x Penicillin/Streptomycin and 1% WEHI-conditioned medium. After 24 hrs. the cells were transfected by nucleofection (Lonza Amaxa kit V, cat# VCA-1003). To this end, 1 x 10^6^ of cells and 1.5 μg of plasmid obtained from GeneCopoeia with *Polq* promoter [NM_001159369 (cat# MPRM33982-PG04)] were used for nucleofection following the manufacturer protocol. After nucleofection, the cells were transferred to 3 mL of growth medium without antibiotics. Activity of Gaussia luciferase and Alkaline Phosphatase secreted to medium was measured after 48 hrs. using Secrete-Pair Dual Luminescence Assay Kit (GeneCopoeia, cat# LF032), following the manufacturer protocol.

### 3’UTR regulation luciferase assay

The assay was performed as described before (Vekariya et al., 2023). Briefly, cells were seeded in Iscove-modified Dulbecco medium supplemented with 10% FBS, 1% L-glutamine, Penicillin/Streptomycin, and 10% WEHI-conditioned medium. After 24h the cells were transfected by nucleofection (Amaxa SF Cell line 4D-Nucleofector X kit L, cat# V4XC-2024). To this end 1 x 10^6^ cells and 2 μg of plasmid obtained from GeneCopoeia: control (#CmiT000001-MT06), *Polq* 3’UTR of NM_029977.2 (#MmiT106208-MT06) or *Parp1* 3’UTR of NM_007415.3 (#MmiT090879-MT06), were used for nucleofection following the manufacturer protocol. After nucleofection the cells were transferred to 3 mL of growth medium supplemented with 10% WEHI and 10% FBS but without antibiotics. The activity of firefly and Renilla luciferases was checked after 18 hrs. using the Duo-Luciferase Assay kit (Promega, cat# E1910), following the manufacturer’s protocol.

### mRNA stability assay

Cells (3.5 x 10^6^) were simultaneously treated with 5 µg/mL actinomycin D (Selleckchem cat# S8964) and 2 µg/mL cycloheximide (Selleckchem cat# S7418) for 0, 1 and 2 hours followed by washing twice with ice-cold PBS. Cells were lysed in 350 µL TRI-Reagent (MRC, cat# TR-118), total RNA was purified, and cDNA was synthesized by RT-PCR. The quantitative RT-PCR of *Polq* was performed, following the procedure described in *Polq* RT-qPCR. The expression of *Parp1* (ThermoFisher Scientific, cat# Mm01321084_m1) was used as a control.

### Real-time RT-qPCR

RNA was extracted using RNeasy plus mini kit (Qiagen, cat# 74134), following the manufacturers guidelines. The isolated RNA was redissolved in Milli-Q water. Extracted RNA was converted to cDNA using SuperScript IV first-strand synthesis system (Invitrogen #18091050) using manufacturer’s protocol. Predesigned TaqMan assay (Polq Assay ID: Mm00712819_m1) and TaqMan gene expression master mix (Applied Biosystems Assay ID# 4369016) were used for quantitative real-time RT-PCR. Briefly, 5 μl of TaqMan gene expression master mix, 0.5μL TaqMan assay, and 1.5μL cDNA and 3μL nuclease-free water were used for qPCR. The housekeeping gene, ubiquitin C (*Ubc*) (Thermo Fisher Scientific, Assay ID# Mm01201237_m1) was used to normalize gene expression. The thermocycle was established by a default program for *Taqman* chemical reaction in the QuantStudio Real-Time PCR 96-well Instrument (ThermoFisher Scientific). The relative gene expression level was calculated by using comparative ΔCt (Ct*_Polq_* – Ct*_UBC_*) analysis.

### *In vivo* treatment of the FLT3(ITD);*Dnmt3a^KD^* 32Dcl3 leukemia

Ten-week old female NRG mice (The Jackson Laboratory, cat# 007799) were total body irradiated (4 Gy) and injected i.v. with 1 x 10^3^ GFP^+^ FLT3(ITD);*Dnmt3a^KD^*32Dcl3 cells. The treatment regimen began when the percentage of GFP^+^ cells in peripheral blood leukocytes (PBL) reached 5-10%. Mice (10 mice/group) were randomized and treated for 14 consecutive days with (1) vehicle, (2) RP-6685 [80 mg/kg oral gavage (Bubenik et al., 2022)], (3) etoposide [5 mg/kg every two days (Cheema et al., 2011)], and (4) the combination of etoposide and RP-6685. The efficacy of the treatment was evaluated two days after the end of treatment by measurements of GFP^+^ cells in PBL. The median survival time was determined.

### *In vivo* treatment of the primary AML xenografts

Doxorubicin was purchased as 10 mM solution in DMSO. Cytarabine was first solubilized in DMSO (50 mg/ml). Both drugs were further diluted in PBS prior to injection. Quizartinib was solubilized in DMSO (100 mg/ml) and prior to injection diluted freshly in 10% Kolliphor HS15 (Sigma 42966) in PBS. RP-6685 was diluted in DMSO (100 mg/ml) and freshly resuspended in 10% D-α-Tocopherol polyethylene glycol 1000 succinate (Sigma 57668) + 5% 1-Methyl-2-pyrrolidionone (Sigma 443778) before administration. NRGS mice (Jackson Laboratory Strain 024099) were sublethally irradiated at 500 Gy and the following day injected via tail vain with 1 x 10^7^ of Lin-primary cells from AML samples: #17550 carried DNMT3A(C537Y) detected in approximately 95% of the cells, and SMC1A(G1139W), CHEK2(K416Rfs*) and NRAS(G12D) (Toma et al., 2025); #13 carried DNMT3A(P569Afs*9) [VAF=44.3%], FLT3(ITD) [VAF=53%] and NPM1(W288Cfs*) [VAF=44.9%]; and #16 carried DNMT3A(R882H) [VAF=47%], FLT3(ITD) [VAF=52.5%] and NPM1(W288Cfs*) [VAF=42.3%]. After 2 weeks the mice were randomly assigned to the experimental groups. Mice injected with #13 and #16 AML cells were treated with vehicles, quizartinib [10 mg/kg for 14 days via intraperitoneal (i.p.) injection (Maifrede et al., 2021)], RP-6685 [80 mg/kg for 14 days via oral gavage (o.g.) (Sullivan-Reed et al., 2018)], or the combination of these drugs. Mice harboring #17550 AML cells were treated with vehicles, doxorubicin + cytarabine [5+3 chemotherapy consisting of: 3 days of 1.5 mg/kg doxorubicin and cytarabine 50 mg/kg via intravenous injection (i.v.), followed by another 2 days of cytarabine i.p. (Porazzi et al., 2022)], RP-6685 [80 mg/kg for 14 days via o.g.] or the combination of all 3 drugs. The mice were euthanized 2 months after the treatment and human CD45+ cells were detected in peripheral blood and bone marrow as described before with modifications (Toma et al., 2024; Vekariya et al., 2023). Briefly, erythrocytes were lysed with ACK lysis buffer (Fisher A1049201), cells were washed with PBS and blocked with Anti-Mo CD16/CD32 antibody (Thermo 14-0161-86) for 10 min, followed by 45 min. staining with PE-linked mouse anti-human CD45 antibody (BD Biosciences 555483). The cells were then washed with PBS and fixed in 2% paraformaldehyde. The samples were analyzed using BD Symphony A5 instrument and analyzed using FACSDiva software.

### Statistics

Data are expressed as mean ± standard deviation (SD) from at least 3 independent experiments unless stated otherwise. When conducting subgroup comparisons between two groups, two-tailed unpaired t-test was used for normally distributed variables. Multiple groups were compared using one-way Anova. Median survival time of the mice ± standard error (SE) was calculated by Kaplan-Meier and compared by Log-Rank test. p values less than 0.05 were considered statistically significant; *p<0.05, **p<0.01, ***p<0.001 and ****p<0.0001.

### Institutional approvals

The patient samples were collected after obtaining informed consent in concordance with the Declaration of Helsinki and were approved by the institutional review boards. Animal experiments were approved by the Institutional Animal Care and Use Committees review board at Temple University. Mice were housed in a temperature- and light-controlled animal facility under a 12-h light/dark cycle and were provided with standard food and water ad libitum.

## Key Resources Table

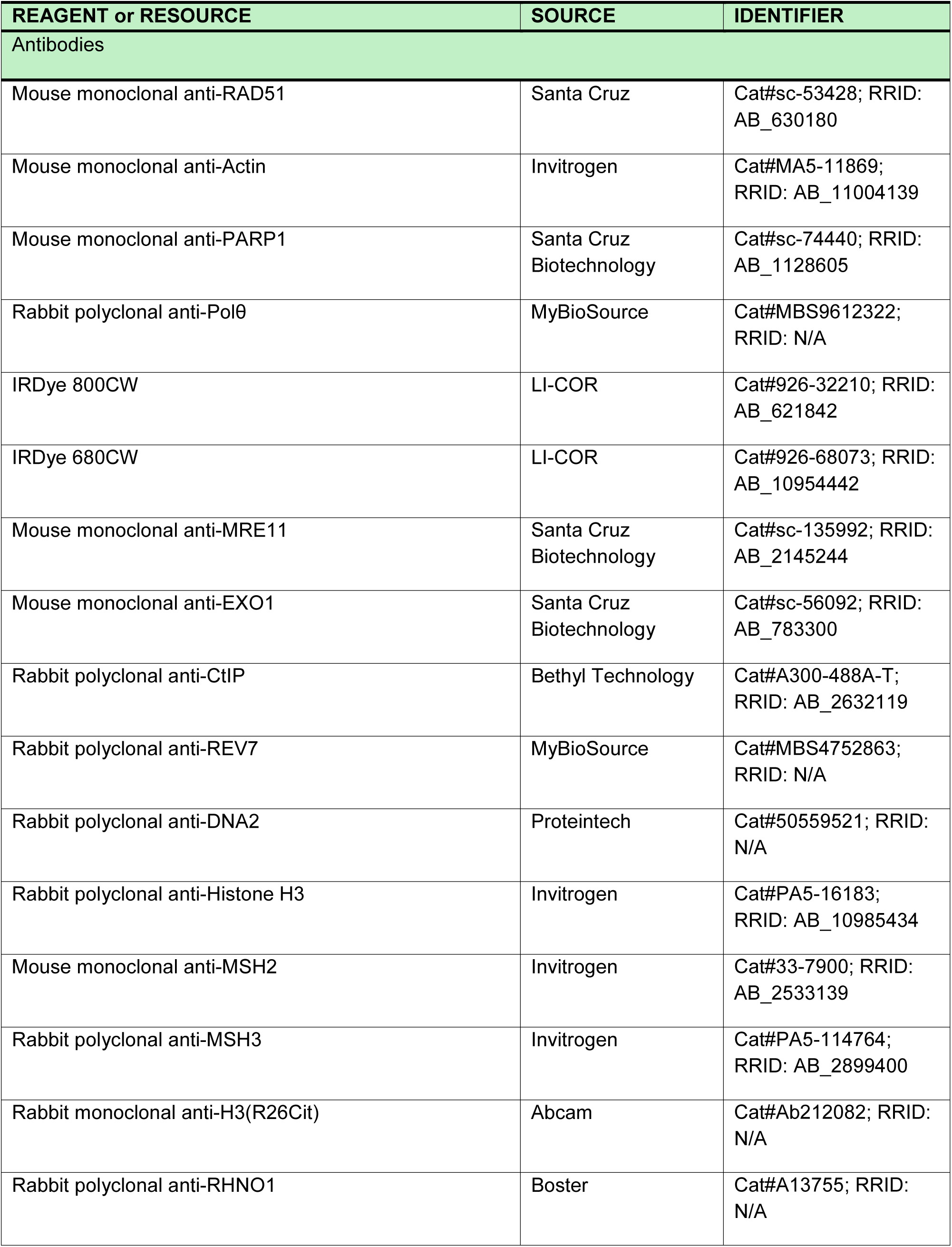
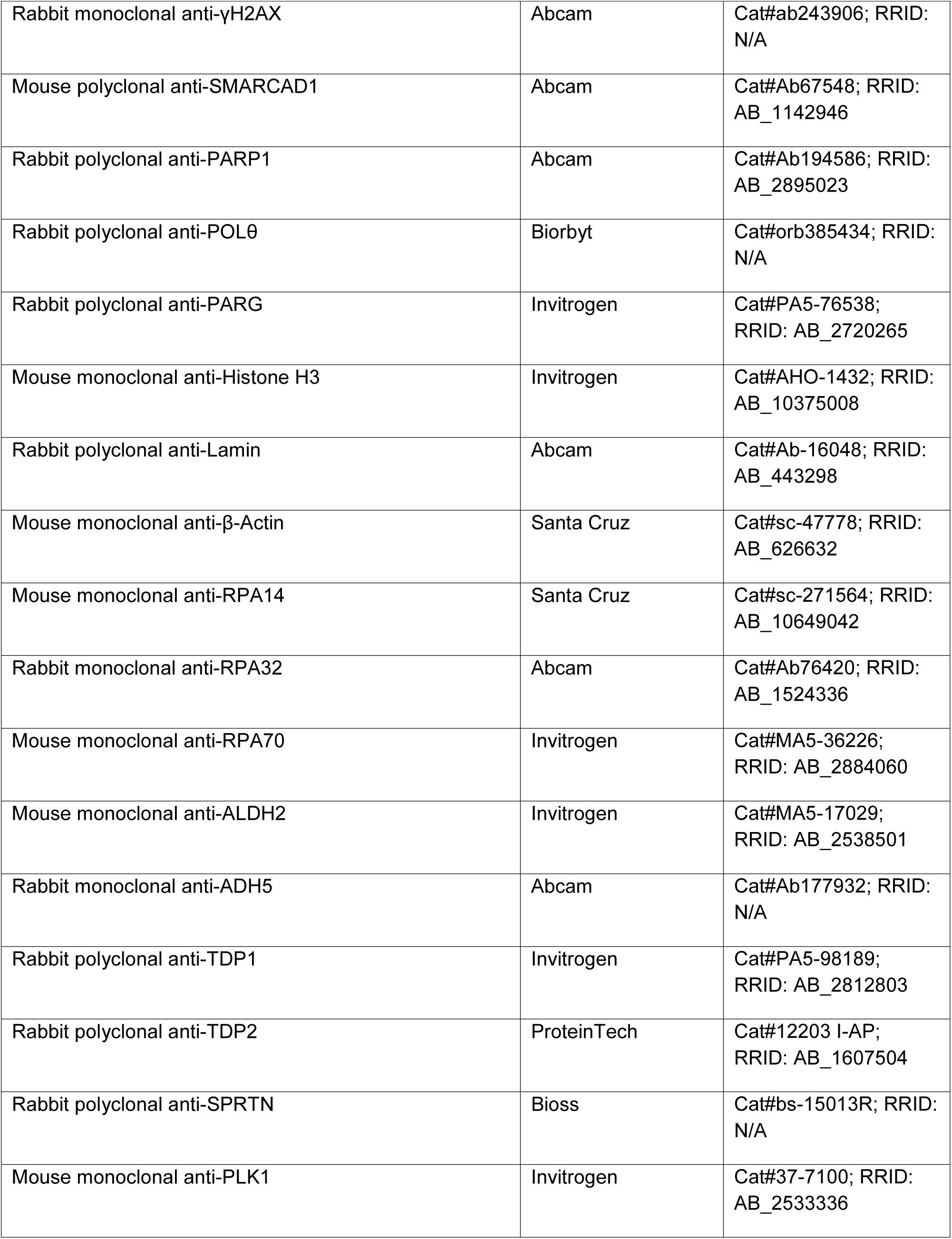
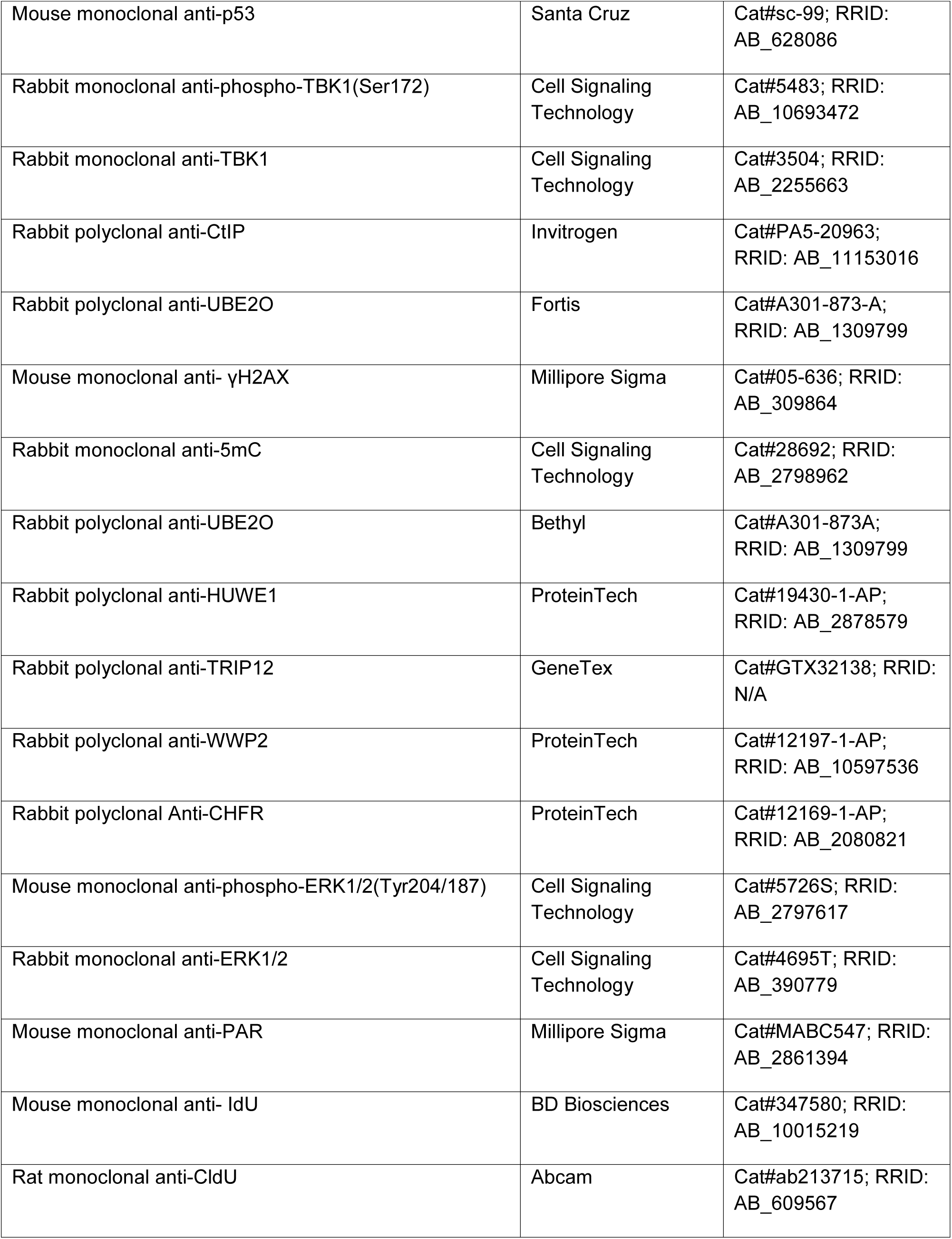
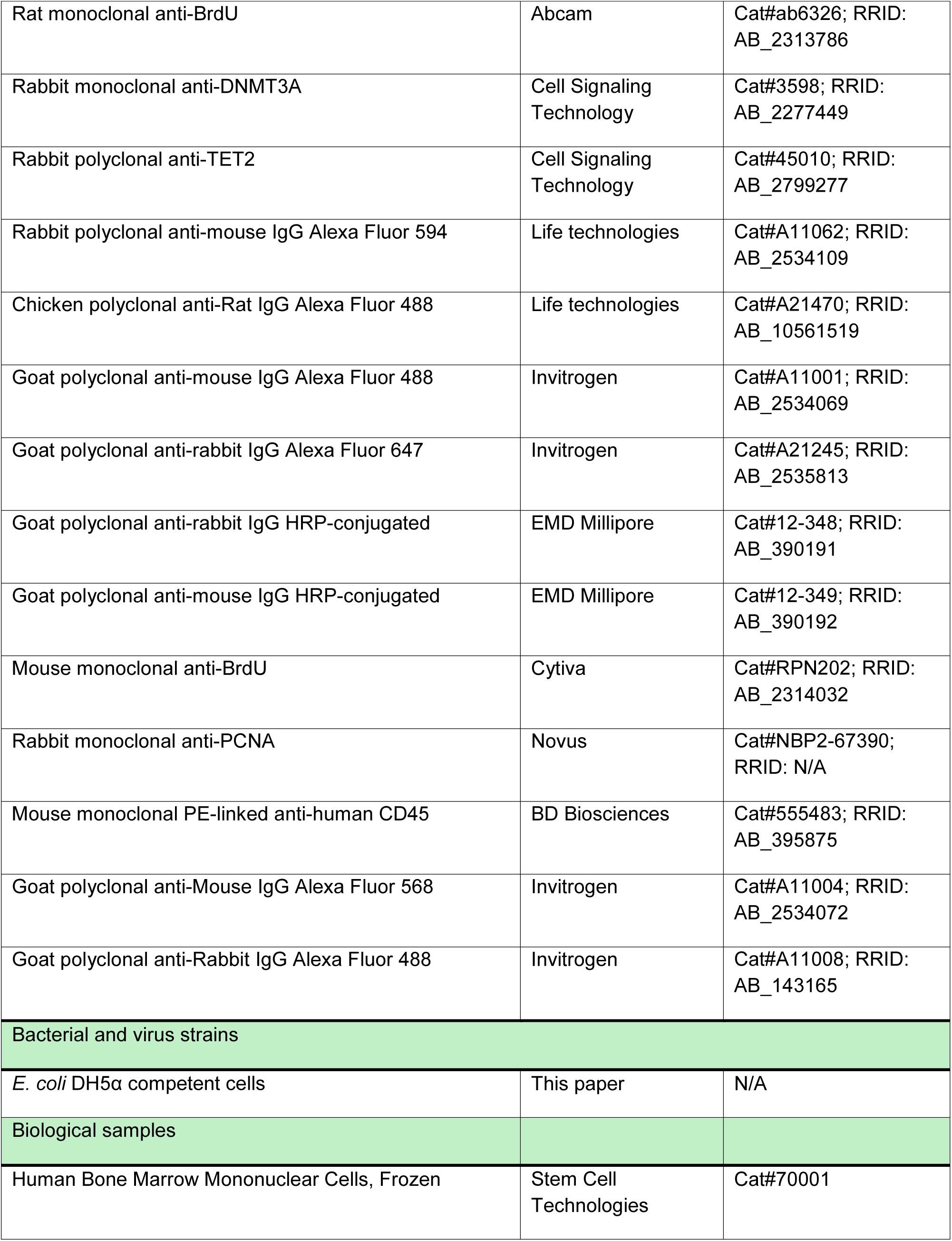
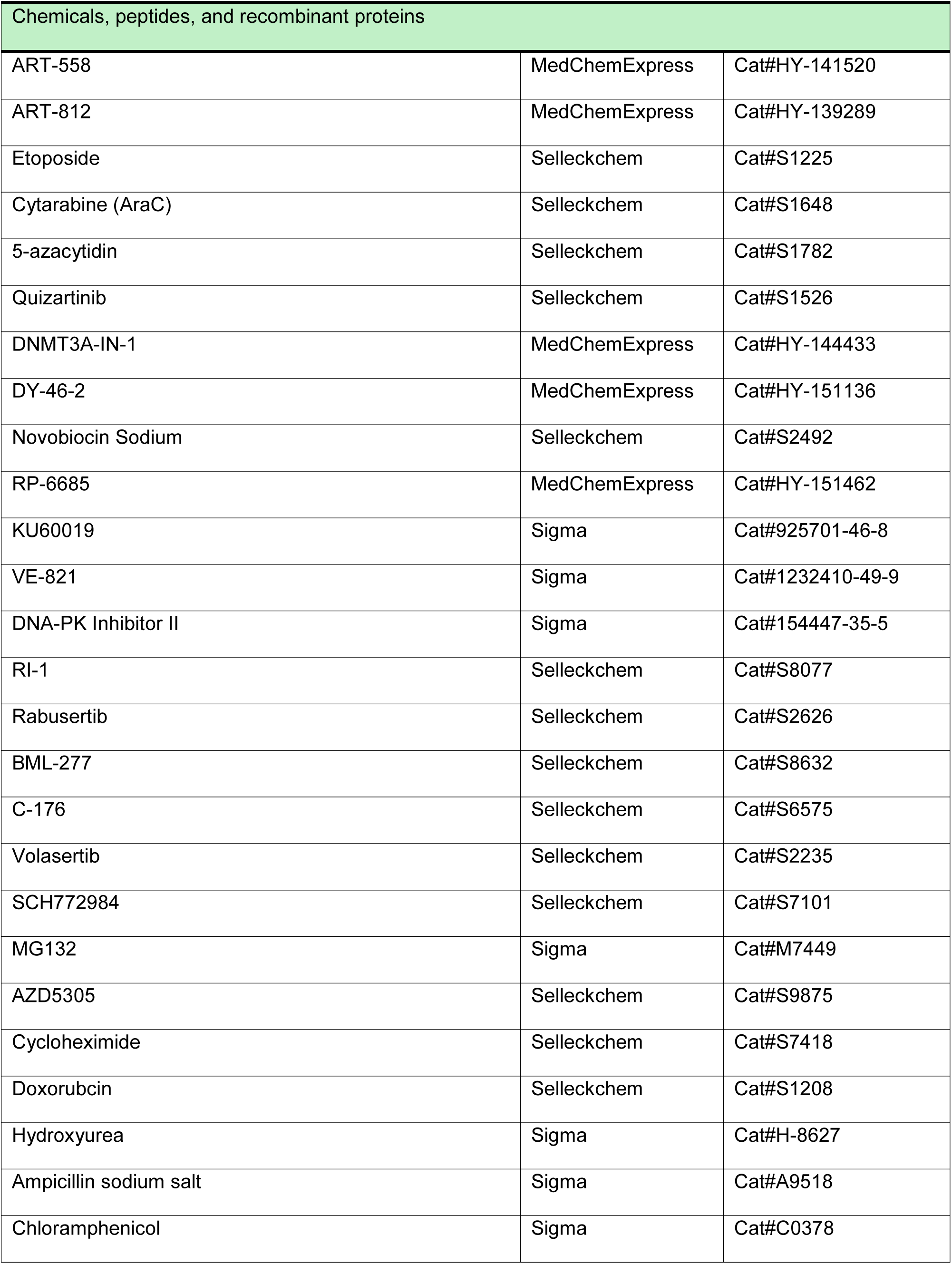
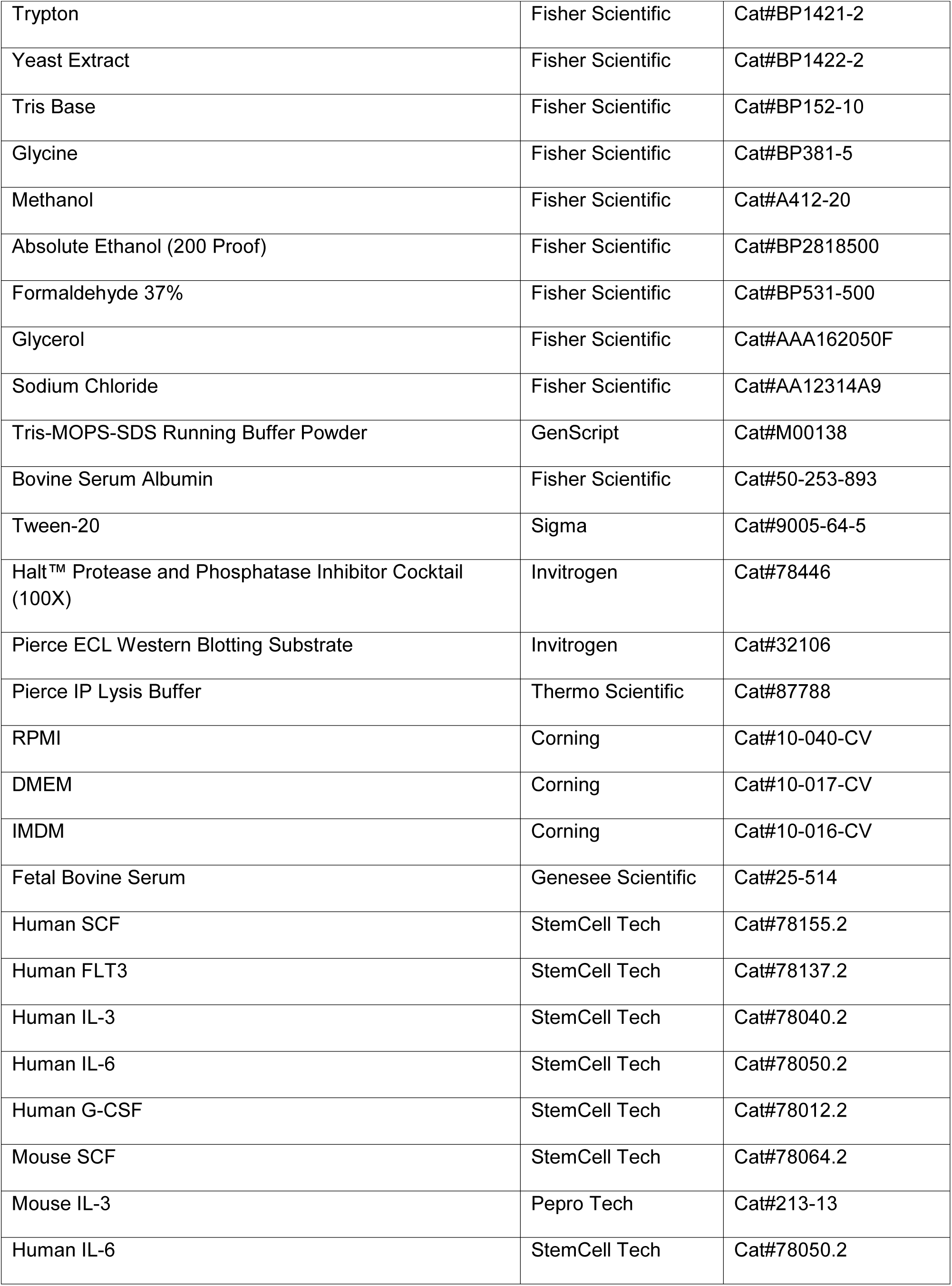
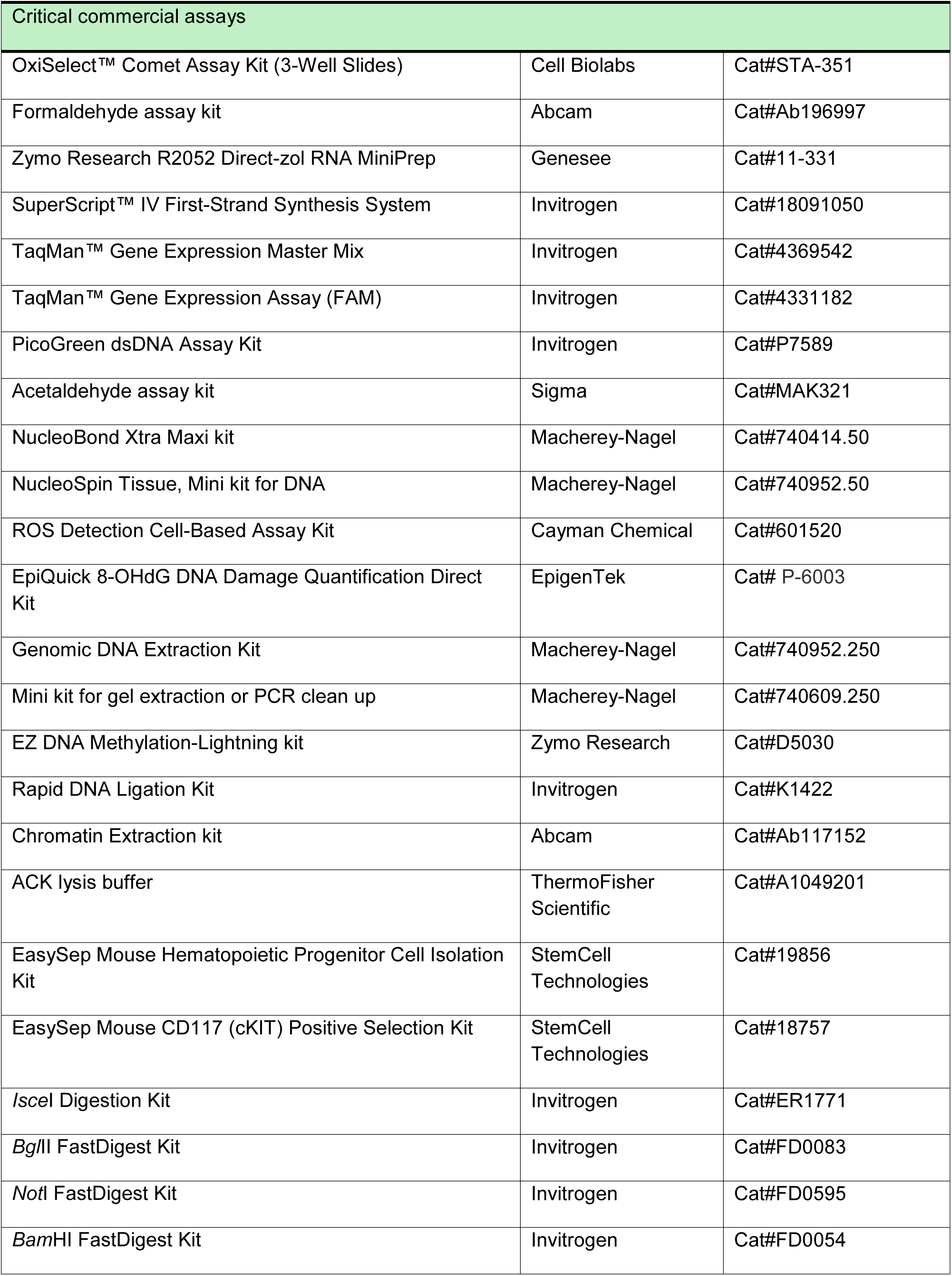
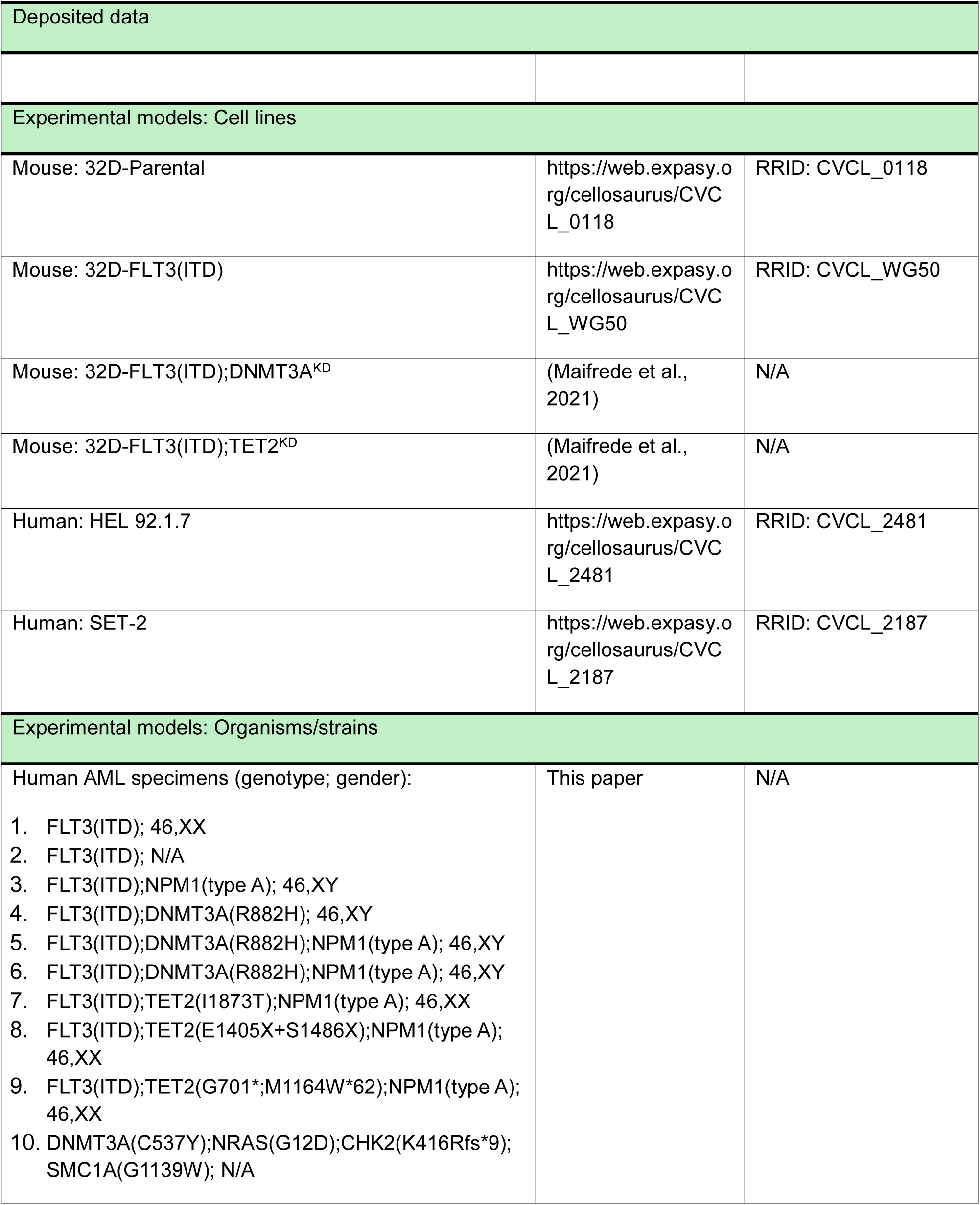
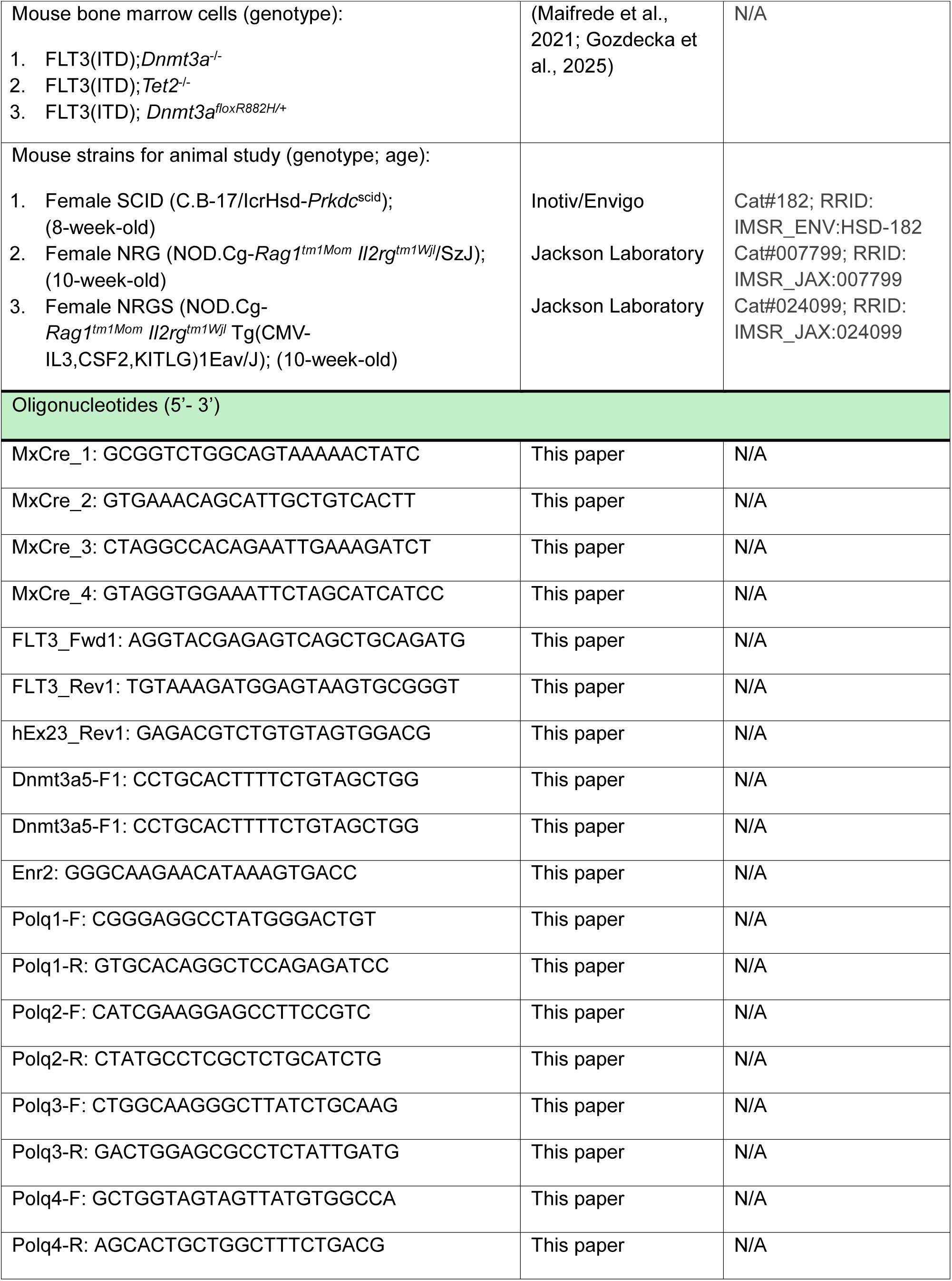
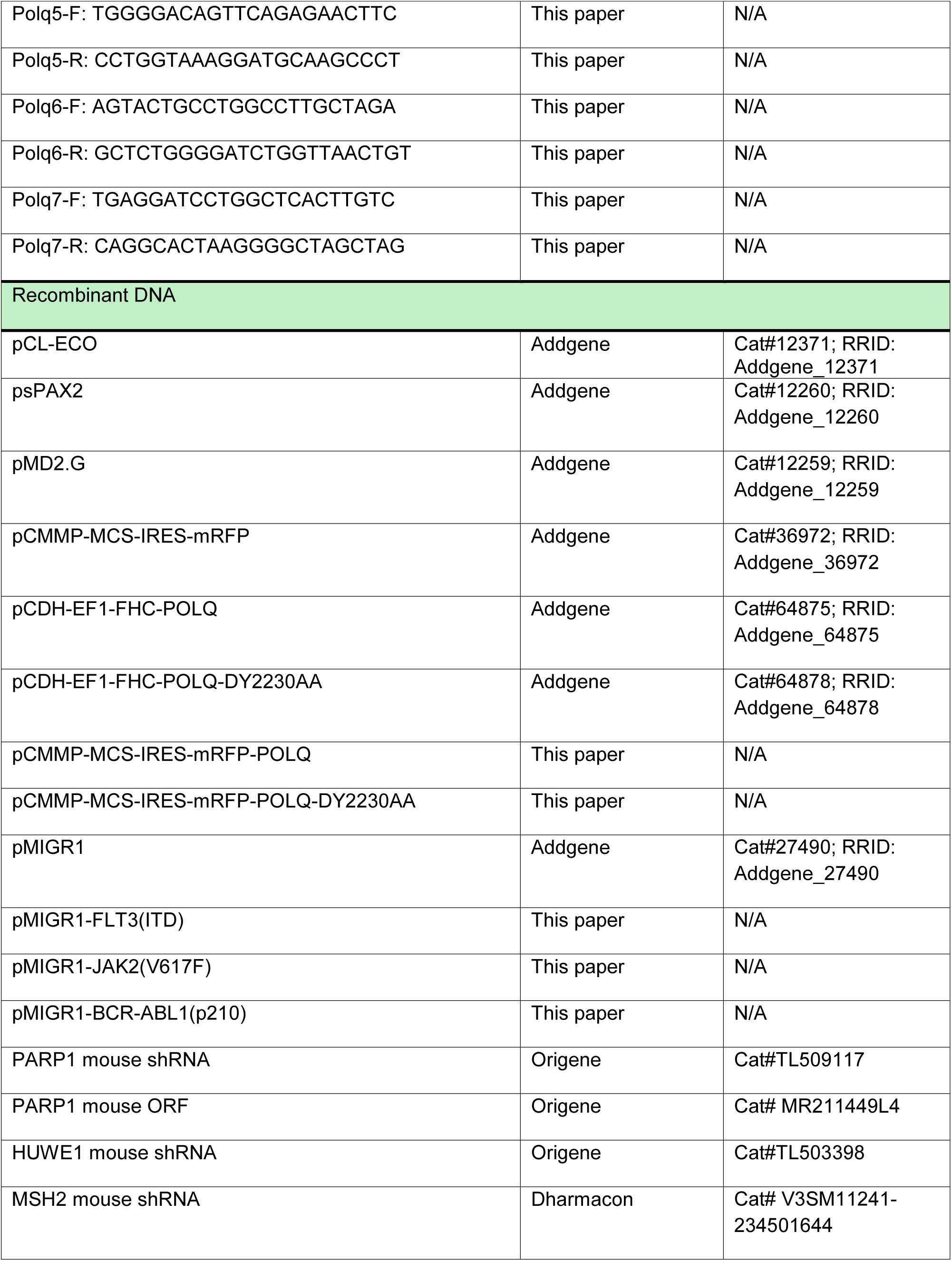
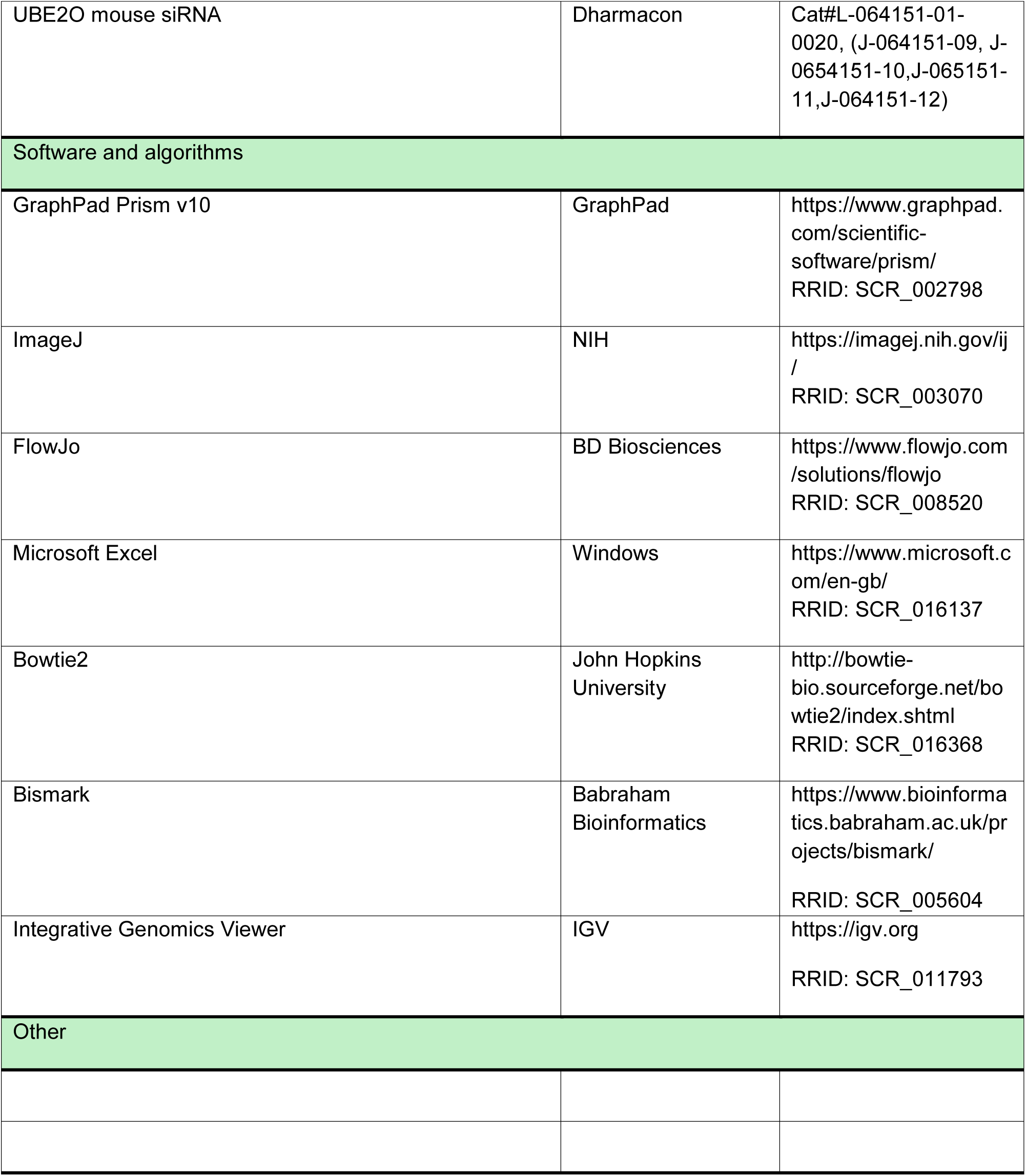

## Supplemental Text and Figures

**Supplemental Figure S1.**
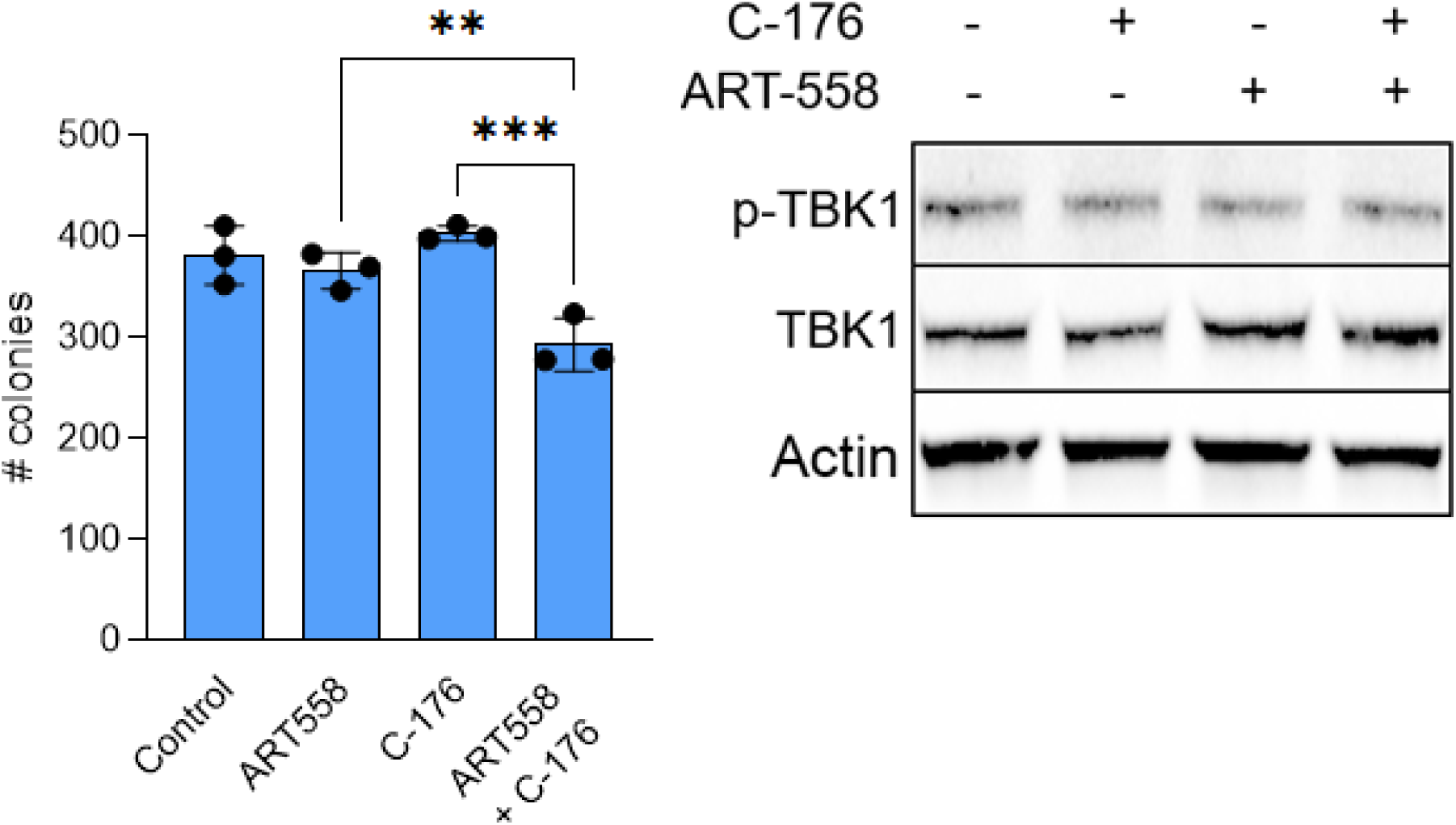
related to Figure 1. The effect of ART558 +/- C-176 in FLT3(ITD);*Tet2^KD^* 32Dcl3 cells. Cells were untreated or treated with 5 μM ART558, 0.5 μM C-176 and the combination. *Left panel:* Mean number of colonies. *Right panel*: Western blot detecting phospho-TBK1 kinase, total TBK1 and actin (loading control). Error bars represent the SD; p values were determined using one-way Anova.

**Supplemental Figure S2.**
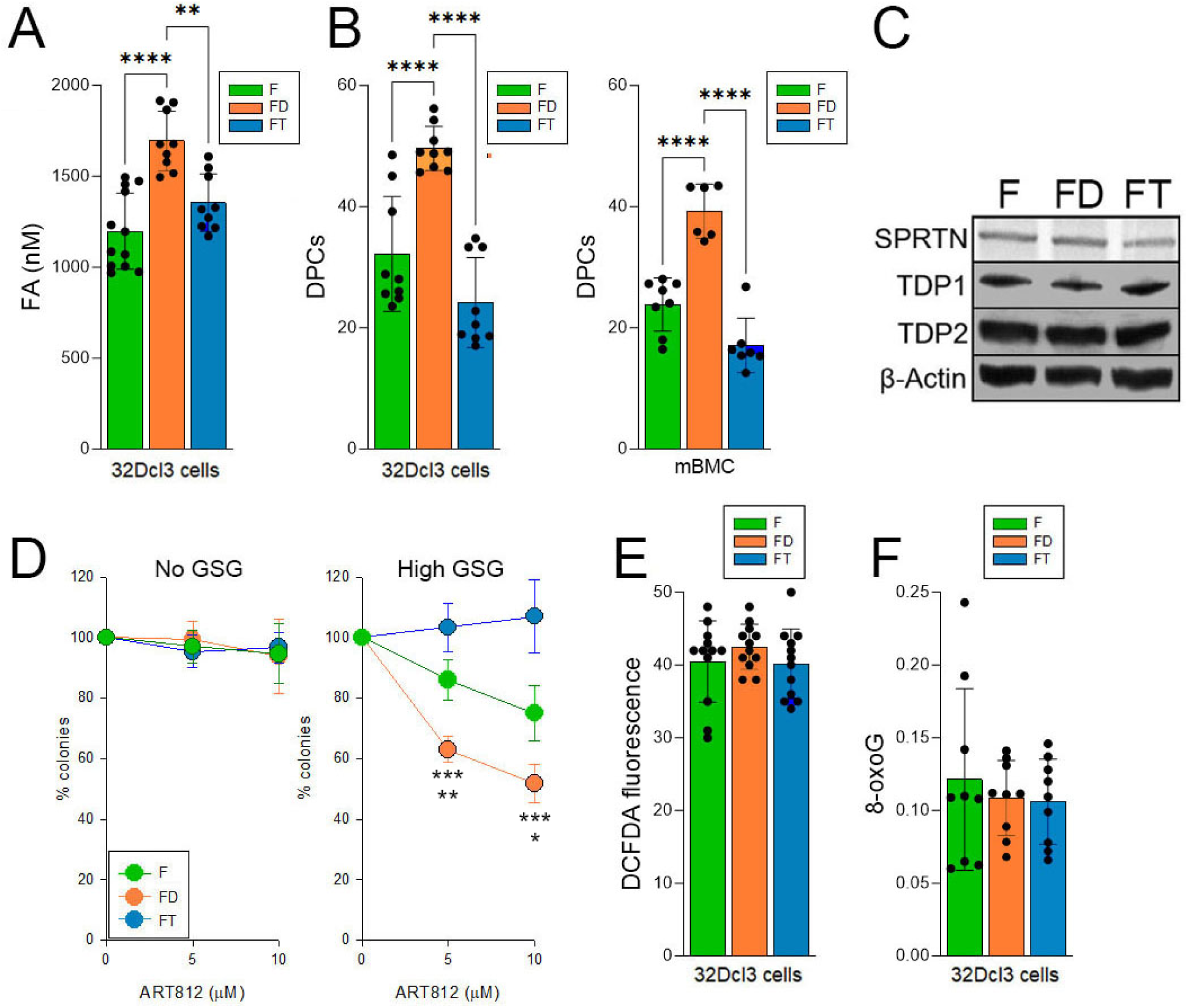
related to Figure 2. Detection of formaldehyde and ROS-induced DNA damage. (**A**) Mean formaldehyde (FA) levels in F, FD and FT 32Dcl3 cells. (**B**) Mean levels of DPCs in F, FD, and FT 32Dcl3 cells and in *Flt3^ITD/ITD^*(F), *Flt3^ITD/ITD^*;*Dnmt3a-/-* (FD) and *Flt3^ITD/ITD^*;*Tet2-/-* (FT) Lin-cKit+ mBMCs. (**C**) Western blot detecting SPRTN, TDP1 and TDP2 in total cell lysates obtained from F, FD, and FT cells. (**D**) Sensitivity of F, FD, and FT 32Dcl3 cells to ART558 in No GSG and High GSG medium. Mean % of colonies when compared to untreated counterparts; p values calculated versus FT (top) and F (bottom) cells. (**E**) Mean DCFDA fluorescence staining using the Detection Cell-Based Assay Kit (Cayman Chemical # 601520). (**F**) 8-oxoG was detected according to the protocol from the EpiQuick 8-OHdG DNA Damage Quantification Direct Kit (EpigenTek P-6003). Genomic DNA was extracted using DNA extraction kit (Macherey-Nagel 740952.250). Error bars represent the SD; p values determined using one-way Anova.

**Supplemental Figure S3.**
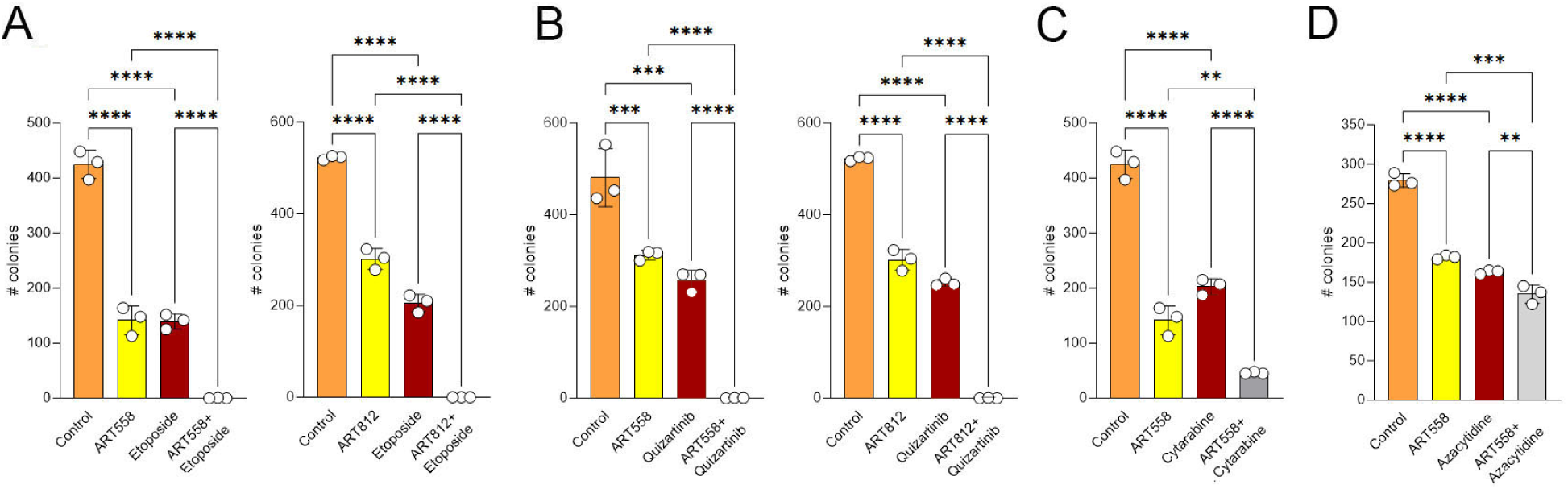
related to Figure 3. Polθi enhanced the sensitivity of DNMT3A deficient leukemia cells to standard drugs. Sensitivity of FLT3(ITD);*Dnmt3a^KD^*32Dcl3 cells to Polθi (25 μM ART558, 50μM ART812) +/- (**A**) 60 nM etoposide, (**B**) 2.5 nM quizartinib, (**C**) 0.2 μM cytarabine, and (**D**) 0.25μM azacytidine. Results represent mean number of colonies. Error bars represent the SD; p values were calculated using one-way Anova.

**Supplemental Figure S4.**
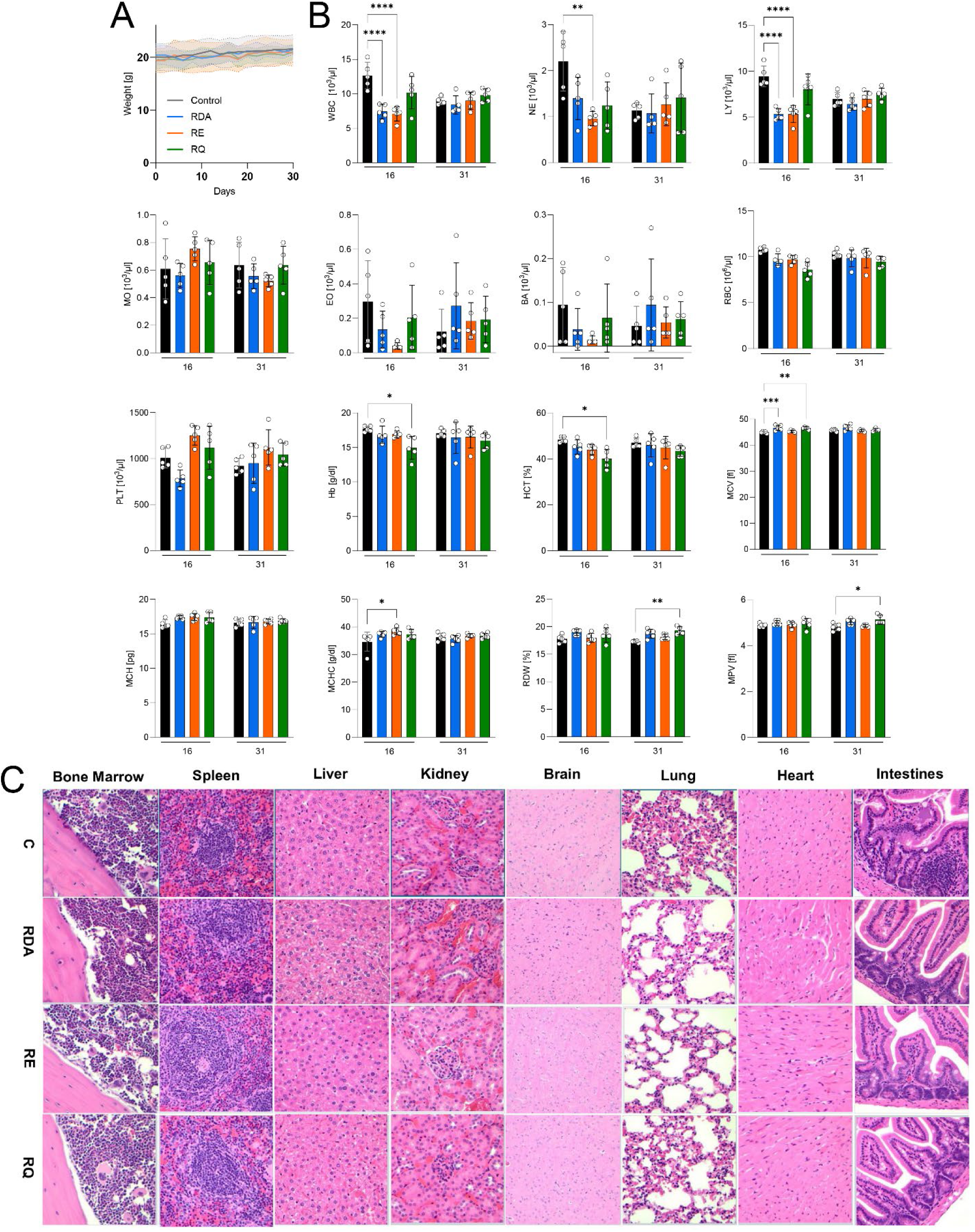
related to Figure 3. Toxicology studies of the combinations of Polθi and standard drugs. BALB/cJ mice (Jackson Lab stock 00651) were randomly assigned to 4 groups (5 mice/group) and treated with vehicles (C - black), and the combination of RP-6685 + doxorubicin + cytarabine (RDA - blue), RP-6685 + etoposide (RE - orange), or RP-6685 + quizartinib (RQ - green). All drugs were used in the same dosing as described in the Materials and Methods of the main text. (**A**) The weight of the animals was determined every other day starting from the first treatment day for a total of 30 days. The values are presented as mean value +/- SD represented by the shaded areas. (**B**) Mean ± SEM of the indicated peripheral blood parameters analyzed on day 16 (2 days after the last treatment) and day 31 (17 days after the last treatment): white blood cells (WBCs), red blood cells (RBCs), platelets (PLT), neutrophils (NE), lymphocytes (LY), monocytes (MO), eosinophils (EO), basophils (BA), hemoglobin (Hb), hematocrit (HCT), mean corpuscular volume (MCV), mean corpuscular hemoglobin (MCH), mean corpuscular hemoglobin concentration (MCHC), red cell distribution width (RDV), and mean platelet volume (MPV). Results represent mean ± SD; each dot represents a result from an individual mouse. * p<0.05, ** p<0.01, *** p<0.001, ****p<0.0001 using one-was Anova. (**C**) On day 31 the animals were humanely euthanized, and the indicated tissues were harvested to 4% paraformaldehyde. Representative (n=5) histopathology/H&E-stained tissue sections of the bone marrow, spleen, liver, kidney, brain, lung, heart, and intestine are shown (40x).

**Supplemental Figure S5.**
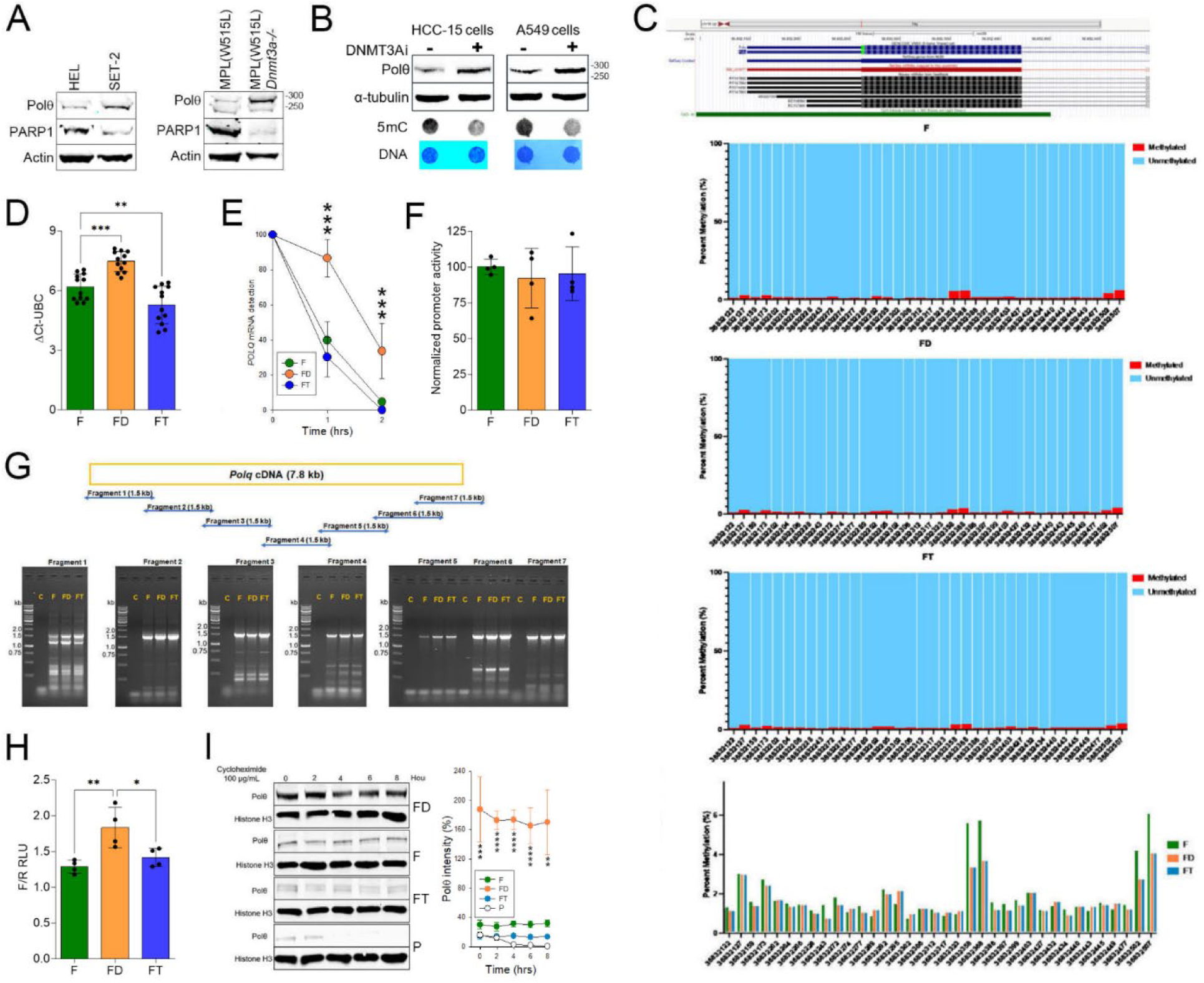
related to Figure 4. Mechanisms regulating Polθ overexpression in FLT3(ITD)-positive DNMT3A deficient leukemia cells. (**A**) Polθ and PARP1 were detected by Western blot in total cell lysates from: *Left panel*: human JAK2(V617F)-positive HEL and JAK2(V617F);DNMT3A(R882H) cells; *Middle panel:* MPL(W515L) and MPL(W515L);*Dnmt3a^-/-^*mBMC. (**B**) Polθ was detected by Western blot in total cell lysates from HCC-15 and A549 lung carcinoma cells treater for 6 days with DNMT3Ai (4 μM DNMT3A-IN-1 or 1 μM DY-46-2). Inhibition of DNMT3A was confirmed by dot-blot detecting downregulation of 5mC. Results represent 3 independent experiments. (**C**) CpG methylation status of the murine *Polq* promoter (coordinates: chr16:36,832,097-36,832,550) by bisulfite amplicon sequencing in FLT3(ITD) (F), FLT3(ITD);*Tet2*-ko (FT) and FLT3(ITD);*Dnmt3a*-ko (FD) 32Dcl3 cells. *Upper panel-* This region includes 35 of the 40 CpG sites of the *Polq* CpG island. *Middle panels-* Graphs depict the averaged percentage of methylation of the individual CpG sites (with their indicated coordinates) in this region of the promoter in the three cell lines, whereby methylation status is depicted by color as follows: red, methylated; blue, unmethylated. A schematic of mouse chromosome 16 and the 5’ region of the *Polq* gene (from UCSC Genome Browser) are shown above the graphs. *Bottom panel-* Bar graph summarizing averaged percent of methylation of the individual CpG sites (with their indicated coordinates) in F, FD and FT cells. (**D**) Mean values of *Polq* mRNA detected in F, FD and FT 32Dcl3 cells by real-time PCR. (**E**) Mean values of *Polq* mRNA stability; p values determined using one-way ANOVA. (**F**) Mean values of *Polq* transactivation assay in F, FD and FT 32Dcl3 cells. (**G**) Agarose gel electrophoresis of 7 RT-PCR-amplified *Polq* fragments in F, FD and FT cells; C – control. Locations and sizes of the expected RT-PCR fragments through *Polq* cDNA sequence is shown. (**H**) *Polq* 3’UTR luciferase assay in F, FD and FT 32Dcl3 cells. Mean activity of Firefly luciferase activity (synthesized under control of the 3’UTR) normalized to Renilla luciferase (constitutively synthesized) in shown. (**I**) Cycloheximide (CHX) chase assay: *Left* – Representative Western blot from nuclear cell lysates obtained from F, FD, FT and parental (P) 32Dcl3 cells after CHX treatment (hrs.). *Right* – Mean expression levels of Polθ. Error bars represent the SD; p values determined using one-way Anova (**D**, **E**, **H**, **I**).

**Supplemental Figure S6.**
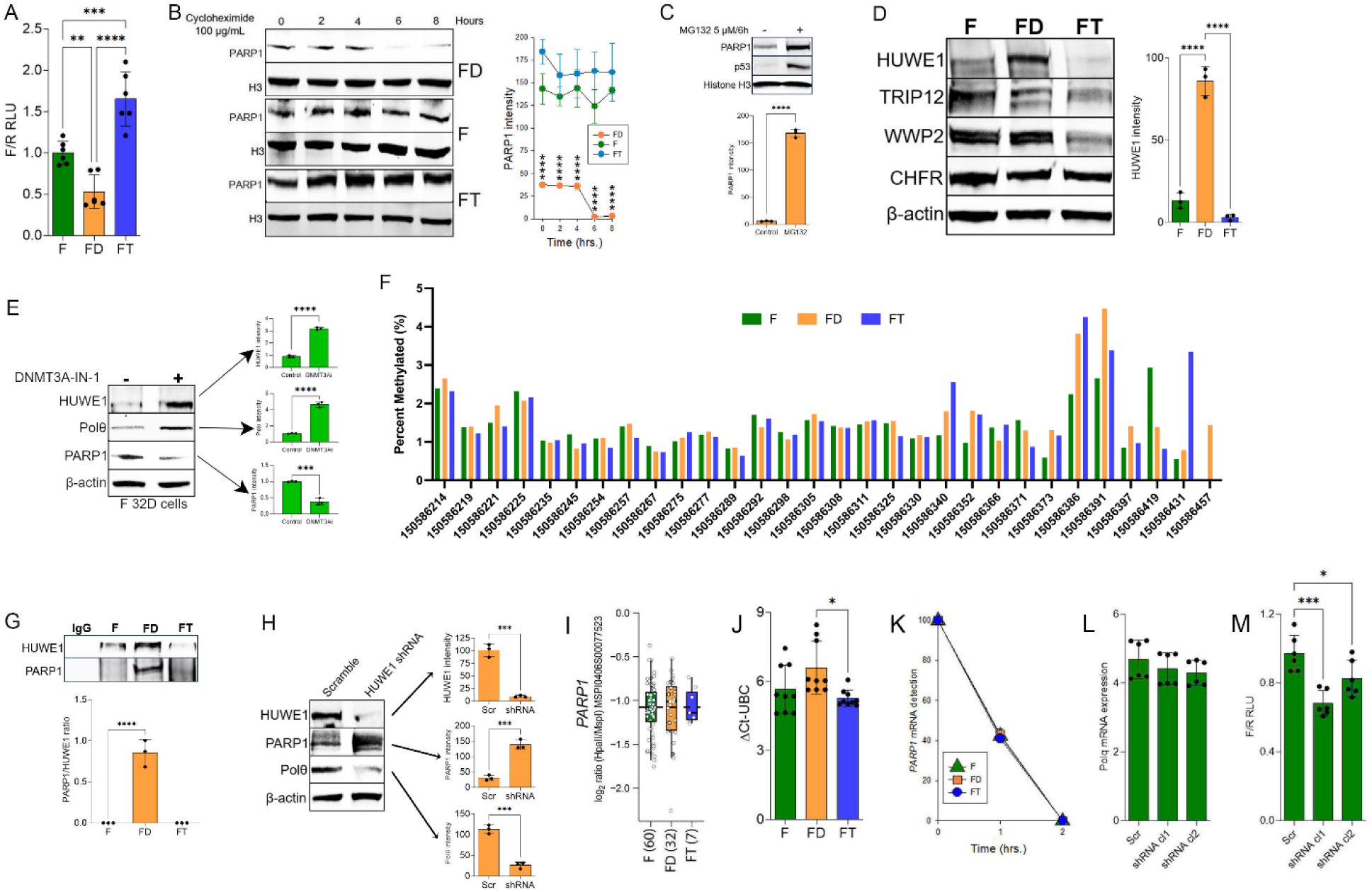
related to Figure 4. Regulation of PARP1 expression in DNMT3A deficient FLT3^ITD^-positive AML cells. (**A**) *Parp1* 3’UTR luciferase assay in FLT3(ITD) (F), FLT3(ITD);*Dnmt3a^KD^* (FD), and FLT3(ITD);*Tet2^KD^* (FT) 32Dcl3 cells. Mean ± SD of the Firefly luciferase activity (synthesized under control of the 3’UTR) normalized to Renilla luciferase (constitutively synthesized) in shown. (**B**) Cycloheximide (CHX) chase assay: *Left* – Representative Western blot from nuclear cell lysates obtained from F, FD and FT 32Dcl3 cells after CHX treatment (hrs.). *Right* – Mean expression levels of PARP1. (**C**) *Top –* Representative Western analysis of FD 32Dcl3 cells treated (+) or not (-) with proteasome inhibitor. *Bottom -* Mean of PARP1 intensity. (**D**) *Left* – Representative Western blot from total cell lysates obtained from F, FD and FT 32Dcl3 cells. *Right* – Mean expression levels of HUWE1. (**E**) *Left* – Representative Western blot from total cell lysates from FLT3(ITD) 32Dcl3 cells treated or not (Control) with 4 μM DNMT3A-IN-1 for 10 days. *Right* – Mean expression levels of the indicated proteins. (**F**) CpG methylation status of the murine *Huwe1* promoter (coordinates: ChrX: 150,586,214-150,586,457) by bisulfite amplicon sequencing in FLT3(ITD) (F), FLT3(ITD);*Tet2*-ko (FT) and FLT3(ITD);*Dnmt3a*-ko (FD) 32Dcl3 cells. Bar graph summarizes averaged percent of methylation of the individual CpG sites (with their indicated coordinates) in F, FD and FT cells. (**G**) *Top –* Representative Western analysis of anti-HUWE1 immunoprecipitates from F, FD and FT cells treated. *Bottom -* Mean of HUWE1/PARP1 intensity. (**H**) *Left* – Representative Western blot from total cell lysates from FLT3(ITD);*Dnmt3a^KD^* 32Dcl3 cells transduced with scrambled or *Huwe1* shRNA. *Right* – Mean expression levels of the indicated proteins. (**I**) Mean methylation of the *PARP1* promoter regions in human *FLT3^ITD^;TET2^mut^*(FT), *FLT3^ITD^;DNMT3A^mut^* (FD) and *FLT3^ITD^*(F) cells (number of patient samples in parentheses). DNA methylation was measured in patients from the ECOG E1900 trial using a HELP promoter array (GSE24505). (**J**) Mean mRNA levels of *Parp1* detected by real-time PCR detecting in F, FD and FT 32Dcl3 cells. (**K**) Mean of *PARP1* mRNA stability. (**L**) Mean *Polq* mRNA detected by real-time RT-PCR in F cells transduced with scrambled and *Parp1* shRNA (clone #1 and #2; see Figure 3C). (**M**) Mean *Polq* 3’UTR luciferase activity in F cells transduced with scrambled and *Parp1* shRNA (clone #1 and #2; see Figure 3C). Error bars represent the SD; p values determined using one-way Anova (**A, D, G, J, M**) and Student’s t-test (**C, E, H**).

**Supplemental Figure S7.**
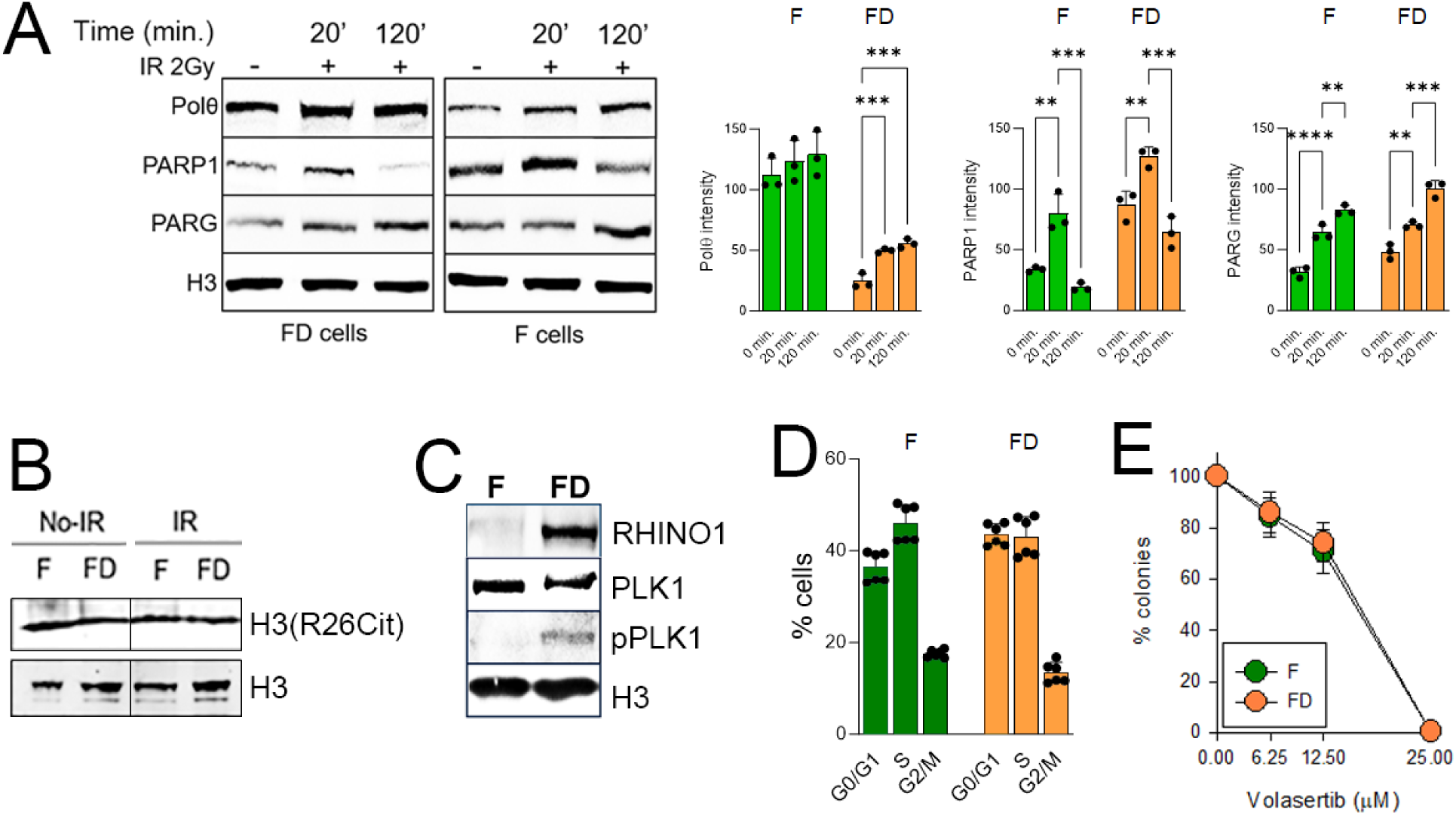
related to Figure 6. *DNMT3A* deficiency modulates chromatin interacting proteins and nuclear proteins in FLT3(ITD)-positive leukemia cells. Cells expressing FLT3(ITD) (F) and 32D-FLT3(ITD) cells lacking DNMT3A (FD) were untreated or irradiated (2Gy). (**A**) *Left* – Representative Western blot of chromatin extracts obtained from F and FD 32Dcl3 cells after 2Gy irradiation (min.). *Right* – Mean expression levels of Polθ, PARP1 and PARG. (**B, C**) Western blots detecting histone H3 citrullinated on Arg26 (**C**) and RHINO1, PLK1, pPLK1 = PLK1 phosphorylated on Thr210 in the nuclear lysates analysed at 2 hrs after irradiation. Histone 3 (H3) served as loading control. (**D**) Cell cycle analysis of untreated cells. (**E**) Sensitivity to PLK1 inhibitor volasertib. Results represent mean % of colonies ± SD when compared to untreated cells. Error bars represent the SD; p values determined using one-way Anova.

**Supplemental Figure S8 related to Discussion.**
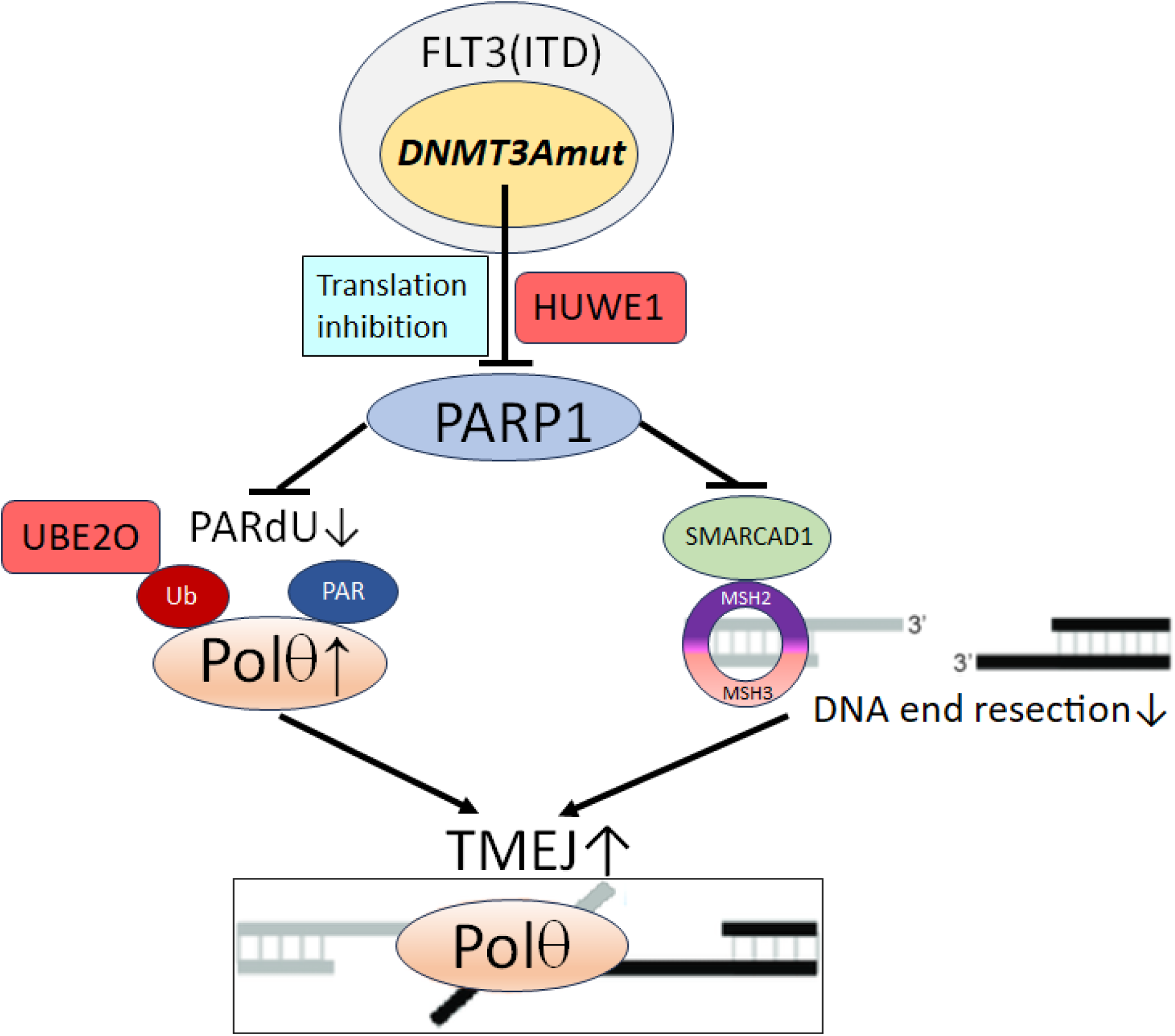
*DNMT3A* mutation enhances Polθ expression and limits DNA end resection to promote TMEJ in FLT3(ITD)-positive leukemia cells. *DNMT3Amut* inhibits PARP1 translation and enhances HUWE1 ubiquitin E3 ligase-mediated proteasomal degradation resulting in downregulation of PARP1 in FLT3(ITD)-positive leukemia cells. Inhibition of PARP1 enhances the expression of Polθ by abrogation of PARdU-dependent UBE2O ubiquitin E3 ligase-mediated ubiquitination and proteasomal degradation of Polθ. In addition, chromatin assembly of PARP1-SMARCAD1-MSH2/MSH3 complex is diminished, facilitating the loading of Polθ on DSBs resulting in limited DNA end resection and enhanced TMEJ.

**Supplemental Figure S9 related to STAR Methods.**
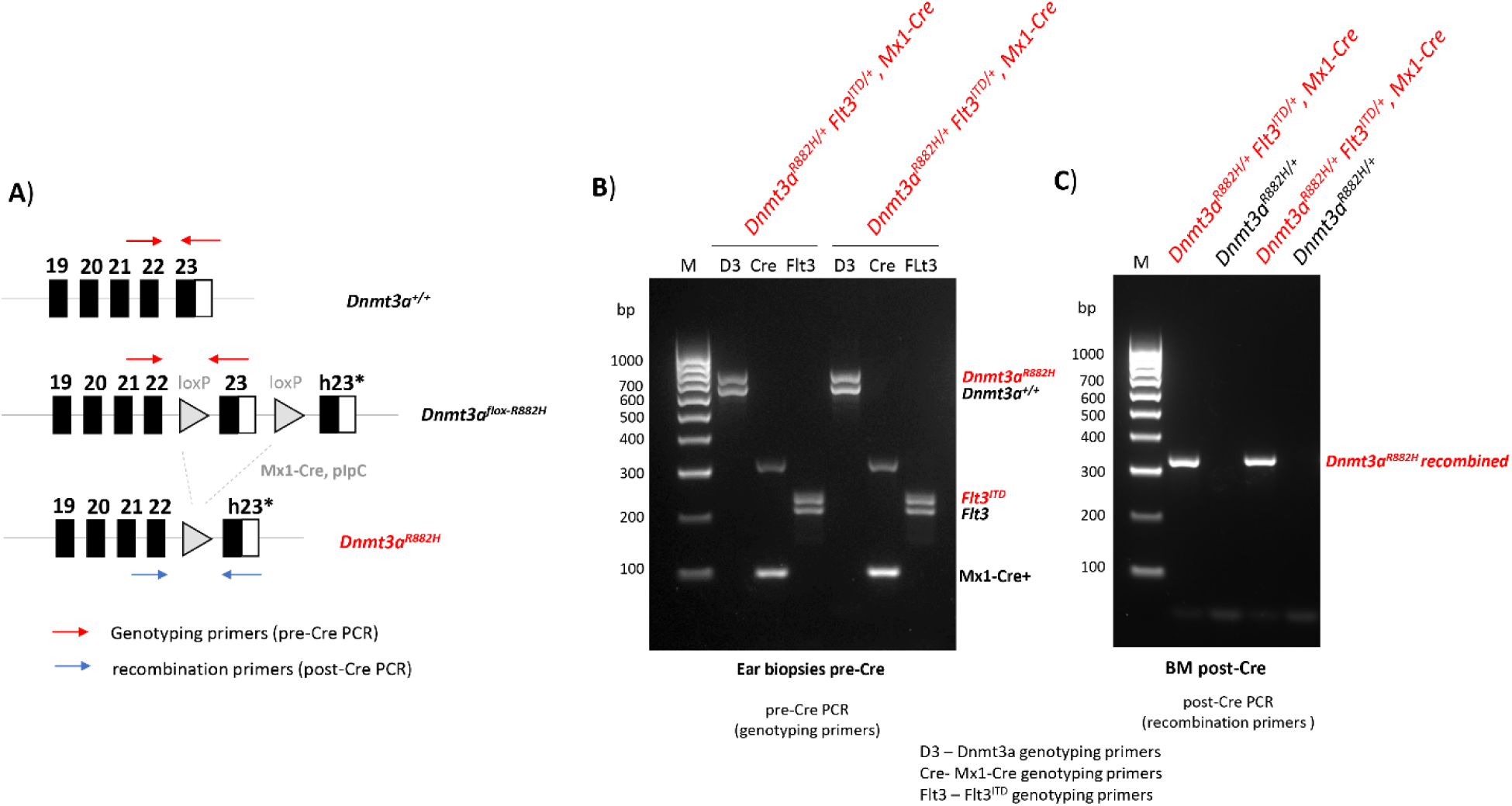
Genotyping validation of *Dnmt3a^R882H/+^ Flt3^ITD/+^*and *Mx1-Cre* alleles. (**A**) Schematic representation of *Dnmt3a^+/+^*(top), *Dnmt3a^flox-R882H^* (middle) and *Dnmt3a^R882H^* (bottom) alleles. Last exon of Dnmt3a, exon 23 is flanked with loxP sites. After stop codon human exon 23 with R882H mutation is introduced. *Dnmt3a^flox-R882H^*were crossed with pIpC-inducible Mx1-Cre reporter line. pIpC treatment results in Cre-mediated removal of mouse exon 23 and expression of human exon 23 containing R882H mutation. *Dnmt3a^flox-R882H^* genotyping primers are indicated in red; recombination primers are marked in blue. (**B**) Genotyping confirming presence of each allele in two *Dnmt3a^R882H/+^ Flt3^ITD/+^, Mx1-Cre* mice before pIpC treatment. D3 – Dnmt3a genotyping primers; Cre-Mx1-Cre genotyping primers; Flt3 – Flt3^ITD^ genotyping primers. (**C**) Validation of efficient *Dnmt3a^flox-R882H^*recombination in bone marrow (BM) extracted from *Dnmt3a^R882H/+^ Flt3^ITD/+^, Mx1-Cre* mice post pIpC treatment.

